# Activating p53^Y220C^ with a Mutant-Specific Small Molecule

**DOI:** 10.1101/2024.10.23.619961

**Authors:** Xijun Zhu, Woong Sub Byun, Dominika Ewa Pieńkowska, Kha The Nguyen, Jan Gerhartz, Qixiang Geng, Tian Qiu, Jianing Zhong, Zixuan Jiang, Mengxiong Wang, Roman C. Sarott, Stephen M. Hinshaw, Tinghu Zhang, Laura D. Attardi, Radosław P. Nowak, Nathanael S. Gray

## Abstract

*TP53* is the most commonly mutated gene in cancer, but it remains recalcitrant to clinically meaningful therapeutic reactivation. We present here the discovery and characterization of a small molecule chemical inducer of proximity that activates mutant p53. We named this compound <u>TR</u>anscriptional <u>A</u>ctivator of <u>p</u>53 (**TRAP-1**) due to its ability to engage mutant p53 and BRD4 in a ternary complex, which potently activates mutant p53 and triggers robust p53 target gene transcription. Treatment of p53^Y220C^ expressing pancreatic cell lines with **TRAP-1** results in rapid upregulation of p21 and other p53 target genes and inhibits the growth of p53^Y220C^-expressing cell lines. Negative control compounds that are unable to form a ternary complex do not have these effects, demonstrating the necessity of chemically induced proximity for the observed pharmacology. This approach to activating mutant p53 highlights how chemically induced proximity can be used to restore the functions of tumor suppressor proteins that have been inactivated by mutation in cancer.

## Main

The p53 protein is a transcription factor that activates multiple antiproliferative signaling pathways, including cell cycle arrest and apoptosis, in response to DNA damage, ribosomal stress, and oncogenic signaling^1,2^. In the absence of these stresses, the ubiquitin ligase Mouse double minute 2 homolog (MDM2) keeps p53 protein levels low^1^. When this negative regulation is released, p53 activates anti-neoplastic programs, such as cell cycle arrest, senescence, apoptosis and differentiation^3^. Due to these functions, *TP53* is the most mutated gene found in cancer with ∼50% of patients showing somatic mutation across a range of tumor types^4^. Approximately 80% of sequenced p53 lesions are missense mutations that cluster in the DNA-binding domain^5,6^. These mutations cause reduced thermal stability and/or impaired DNA binding, thus compromising the tumor suppressor function^7^.

For decades, restoring p53 function has been a goal in the development of cancer therapeutics. These efforts have stimulated the development of MDM2 inhibitors, such as nutlin-3a, which lead to stabilization of p53^WT^ ^8^. Severe thrombocytopenia limits the therapeutic use of nutlin-3a and related molecules^9,10^ (**Fig. 1A**). Another approach to restoring p53 function is to stabilize the mutated protein with directed small molecule ligands. Recently, several groups have developed small molecule p53 “correctors” that increase the thermal stability and partially restore the transcriptional activity of mutated p53^11^. These p53 corrector molecules target a structural cavity created by p53^Y220C^ ^12^, a hotspot mutation affecting ∼120,000 patients per year and accounting for approximately 1.6% of tumor p53 missense mutations^4,6,13,14^. Several successful medicinal chemistry campaigns have targeted the p53^Y220C^ pocket to improve the thermal stability of mutant p53 (**Fig. 1A**). An initial *in silico* screen followed by structural investigation identified a carbazole fragment PhiKan083 with weak binding affinity to p53^Y220C^ ^15^. Further optimization of this scaffold produced the lead compounds PK5196 and PK9328 with enhanced binding affinity, leading to increased expression of p53 target genes and improved antiproliferative activity in cells^16,17^. Using a similar approach, PMV Pharmaceuticals and Jacobio Pharmaceuticals have developed the mutant-specific small molecule p53 binders rezatapopt (PC14586) and JAB-30355^18,19^, respectively. In addition, the presence of a reactive cysteine in the p53^Y220C^ inspired synthesis of covalent-acting fragments such as KG13, which selectively targets mutant p53 via covalent modification of the sulfhydryl group of the cysteine 220 side chain using a methyl acrylamide warhead^20^. Despite these advances, it is not clear to what extent partial p53^Y220C^ functional restoration will provoke a clinically meaningful outcome. A major unresolved question is whether enhancing the thermal stability of mutant p53 to wild-type (WT) level is sufficient for cancer therapy, or if further activation is also required. Small molecule p53 activators are required to answer this question.

**Figure 1:**
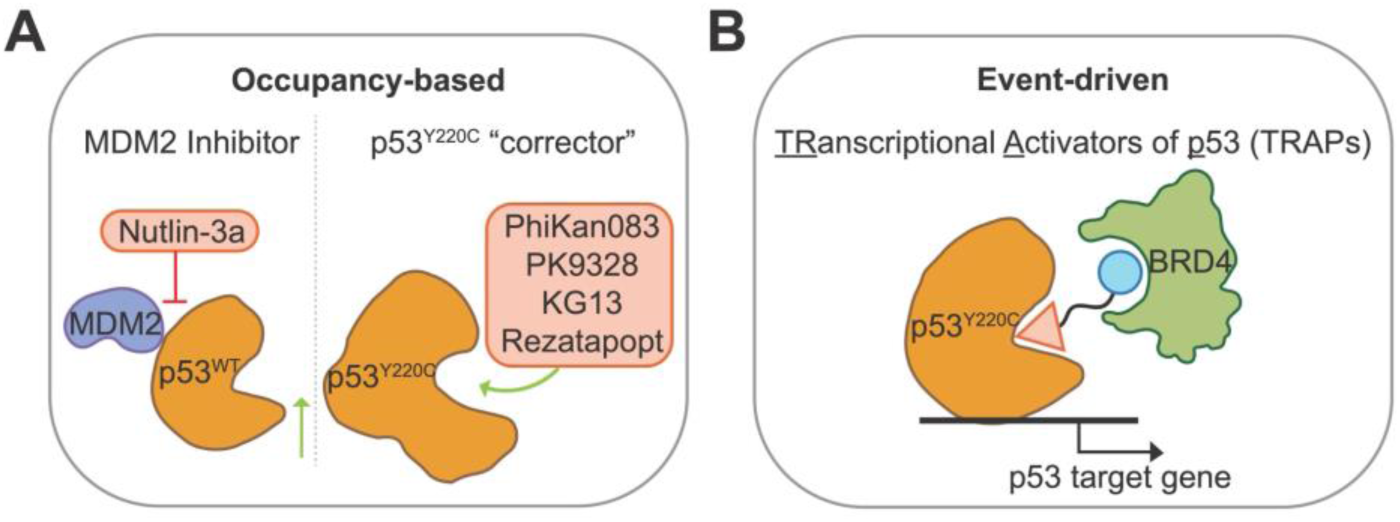
Chemical Targeting of p53 in Cancer Cells. **A.** Schematic of the traditional occupancy-based approach to develop MDM2 inhibitors that interrupt the MDM2-p53^WT^ interaction (left) and p53^Y220C^ correctors that bind to p53^Y220C^ and increase its thermal stability (right). **B.** Schematic of the event-driven transcriptional activator of p53 (TRAP) targeting p53^Y220C^ mutant through the recruitment of BRD4.

The advent of event-driven pharmacology, enabled by chemically induced proximity, has expanded the scope of targets addressable by small molecules. Leveraging molecular proximity as a unifying regulatory principle, small molecules that induce new protein-protein interactions have enabled precise control over cellular signaling, including rewiring of protein degradation pathways^21,22^. These strategies offer major advantages beyond traditional occupancy-based pharmacology: *i*) a single small molecule may catalyze multiple rounds of degradation, transcription, etc., *ii*) efficacy does not require complete target occupancy, and *iii*) the use of dual targets creates a logical “AND” gate, resulting in enhanced cellular and biochemical selectivity. Recently reported p53-targeting small molecules that purport to take advantage of these characteristics include those that induce mutant p53 degradation^23^, p53 acetylation^24,25^, p53 stabilization by deubiquitylation^26^, and selective mutant p53 cancer cell killing by inducing p53-dependent toxicity^27^. Here we seek to leverage the induced proximity strategy to directly activate the core function of mutant p53: transcriptional activity.

To activate p53, we took inspiration from an approach we have recently developed that uses small molecules to reprogram transcription factors. These small molecules, which are called transcriptional/epigenetic chemical inducers of proximity (TCIPs), recruit transcriptional coactivators (BRD4 or CDK9/pTEF-B) to gene promoters bound by the transcriptional repressor protein, BCL6 (B cell lymphoma 6)^28,29^. Treatment of BCL6-high B cell lymphomas with TCIPs rapidly activates a suite of cell death genes normally repressed by BCL6, resulting in tumor cell-specific cell death. Here we apply this logic to develop mutant-specific small molecules <u>TR</u>anscriptional <u>A</u>ctivators of <u>p</u>53 (TRAPs). We report proof-of-concept TRAPs that recruit the transcriptional coactivator protein BRD4 to p53^Y220C^. These small molecules activate p53 and produce exaggerated pharmacology when compared with existing small molecules that can only restore p53 thermal stability. Further development of TRAPs towards clinical candidates has the potential to provide clinical benefit to patients with p53^Y220C^ cancers.

## Results

### Small molecules that specifically activate a p53 mutant

To enable discovery of p53 activator compounds, we established a p53 luciferase reporter assay in BxPC-3 cells. In these cells, the sole remaining *TP53* allele codes for p53^Y220C^ ^30^. We first used this assay to search for small molecule p53 binders that could serve as starting points for bivalent compound synthesis. We screened a series of published p53^Y220C^ correctors^17,19,20^, the MDM2 inhibitor nutlin-3a^8^, and the acetylation targeting chimera (AceTAC) molecule MS78^25^. Among these, the B-1 binder^19^, which shares the same chemotype as the currently reported p53^Y220C^ selective clinical candidates, was the only compound that activated the reporter (**Fig. S1A**). Encouraged by these results, we next synthesized a tool compound, **B-1 linker** (**Fig. S1B**), where the methyl piperidine in B-1 was functionalized with a short alkyl linker through amide bond coupling^31^. **B-1 linker** produced enhanced transcriptional activation in the reporter assay relative to the parental B-1 molecule (**Fig. S1A**). This suggests that functionalization of B-1 with a linker did not impede p53 binding and possibly contributed additional favorable protein-ligand interactions, aiding in the restoration of p53 function.

Knowing that presence of a linker did not affect the functional response, we used **B-1 linker** as the basis for a library of bivalent molecules where the B-1 binder was linked to the bromodomain ligand JQ1^32^ through alkyl, PEG, and rigid diamine linkers (**Fig. 2A**). JQ1 recruits BRD4 without inhibiting its transcriptional activator function^28^. We next asked if the compounds in the library can induce a complex between BRD4 and p53^Y220C^ *in vitro*. To test this, we purified recombinant p53^Y220C^ DNA binding domain and established a time-resolved fluorescent energy transfer (TR-FRET) assay between BRD4_BD1_ and p53^Y220C^ (**Fig. 2B**). Due to the bifunctional nature of the molecules, we expected to observe a reduction of complex formation when excess bivalent molecules saturate individual protein binding sites rather than forming the ternary complex, a phenomenon commonly called the “hook effect”^33^. Among the tested compounds, those with methylene-connected (**462**, **463**) or spiro (**464**) six-membered heterocycle linkers showed bell-shaped curves, with the greatest TR-FRET ratios at the peak and a visible hook effect (**Fig. 2B**), while compounds with smaller rigid linkers, flexible alkyl, and PEG linkers produced much weaker signals (**Fig. S1C**). Additionally, minor structural changes in the linkers, such as removing the methylene spacer (**537**) or replacing one nitrogen atom on the six-membered ring (**540**), resulted in a rightward shift of the curve (**Fig. S1C**). Taken together, the biochemical data identified compounds **462** (**TRAP-1**), **463** (**TRAP-2**), and **464** (**TRAP-3**) as small molecules that induce proximity between the p53^Y220C^ DNA binding domain and BRD4_BD1_.

**Figure 2:**
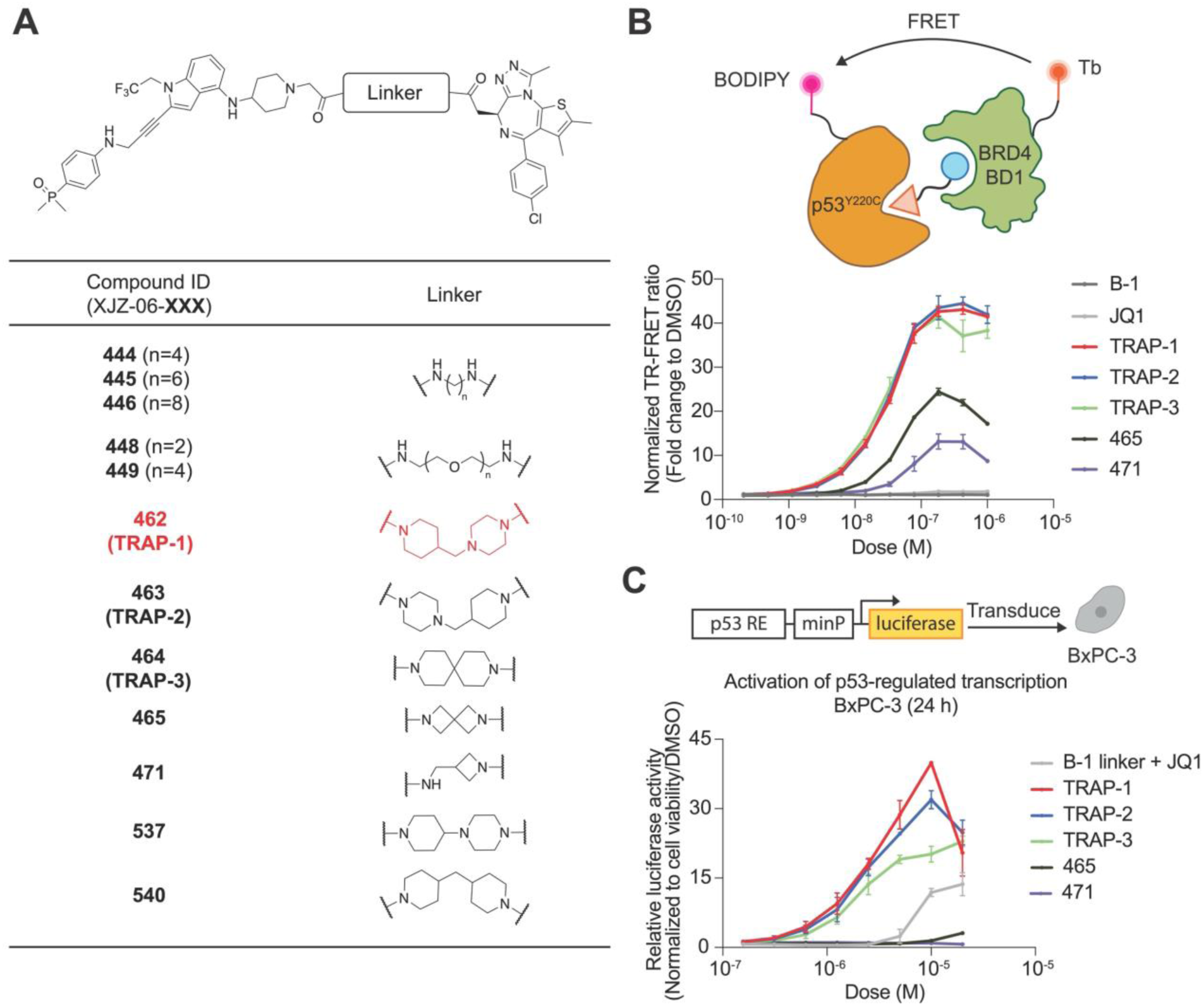
Development of B-1-based bivalent TRAPs. **A.** The structures of the B-1-based bivalent TRAP library. **B.** TR-FRET. The ternary complex formation measured by TR-FRET between purified p53^Y220C^ and BRD4_BD1_. The data is shown as means ± s.d. of n=2 independent experiments. **C.** Luciferase reporter. Activation of p53-regulated transcription in BxPC-3 (p53^Y220C^) cells after compound treatment for 24 h. The data is shown as means ± s.d. of n=2 independent experiments.

Using the p53 reporter assay, we proceeded to screen our bivalent compound library to assess p53-dependent transcriptional activity. We compared these results with co-treatment of cells with the individual binders as a control. JQ1 did not induce p53 target gene activation at concentrations up to 10 µM (**Fig. S1A**). Co-treatment with the **B-1 linker** and JQ1 produced modest reporter activation (12-fold versus DMSO) compared with the **B-1 linker** alone (**Fig. 2C**). In contrast, **TRAP-1**, **TRAP-2**, and **TRAP-3** each produced dose-dependent reporter activation, with **TRAP-1** inducing 40-fold target gene transcription upregulation at 10 µM (**Fig. 2C**). Further increasing the concentration beyond 10 µM resulted in a decrease in luciferase signal (**Fig. 2C**), recapitulating the "hook effect" observed in the TR-FRET dimerization assay. Compounds that showed lower dimerization ability in the TR-FRET assay induced lower or no transcription activation at the concentration range that we tested (**Fig. S1D**). This observation, along with strong overall correlation between reporter gene induction and biochemical binding as measured by TR-FRET (**Fig. S1E**), suggested that transcriptional activation of p53-dependent transcription depends on chemically induced BRD4-p53 proximity. Collectively, the data suggest that **TRAP-1**, **TRAP-2**, and **TRAP-3** are potent p53^Y220C^ transcriptional activators.

We next sought to explore bivalents featuring different p53^Y220C^ selective binders. For this, we focused on PK9328^17^ as the p53^Y220C^ binder which has been previously used for the development of MS78, an acetylation-inducing chimera^25^ (**Fig. S2A**). Treatment with PK9328 alone did not produce transcriptional activation in the reporter assay (**Fig. S1A**), and the synthesis of a similar small library of TRAP candidate compounds produced no active compounds in the TR-FRET dimerization (**Fig. S2B**) or luciferase reporter assays (**Fig. S2C**).

### Transcriptional activation requires ternary complex formation

To test whether transcriptional activation by our molecules depends on co-binding to p53^Y220C^ and BRD4, we designed and synthesized negative control compounds with minor chemical modifications that disrupt binding to either p53^Y220C^ or BRD4. Using **TRAP-1** as a template (**Fig. 3A**), **TRAP-1-Neg1** was synthesized with an opposite chiral configuration to reduce BRD4 binding^32^. Likewise, **TRAP-1-Neg2** was designed with two extra methylations on the B-1 nitrogen atoms to remove the H-bond interactions that were shown to be crucial for p53^Y220C^ binding^34^. Compared with the lead compounds, the negative controls showed negligible ternary complex formation in the TR-FRET dimerization assay (**Fig. 3B**) and no transcriptional activation in the reporter assay (**Fig. 3C**). A TR-FRET assay between p53^WT^ and BRD4 also resulted in no TRAP-induced ternary complex (**Fig. S3**). These data confirm that the enhanced transcriptional activation is associated with the ternary complex formation between p53^Y220C^, BRD4, and compound. Furthermore, chemically induced proximity is p53 mutant-specific.

**Figure 3:**
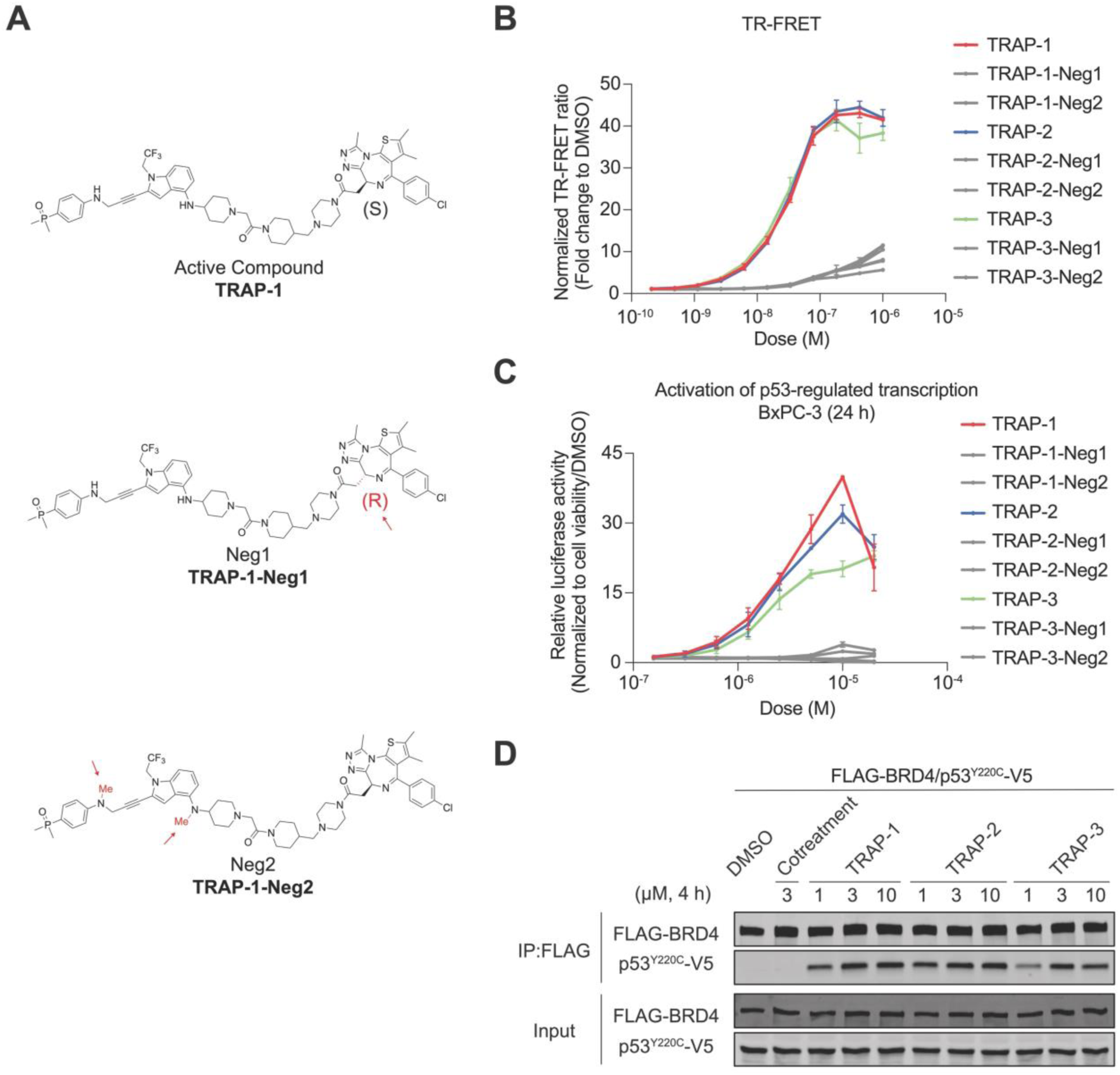
Ternary complex formation is required TRAP-mediated transcription activation. **A.** Structures of negative controls containing minor chemical modifications to remove BRD4 binding (**TRAP-1-Neg1**) or p53^Y220C^ binding (**TRAP-1-Neg2**). **B.** TR-FRET. The ternary complex formation of TRAPs and their negative controls measured by TR-FRET between p53^Y220C^ and BRD4_BD1_. The data is shown as means ± s.d. of n=2 independent experiments. **C.** Luciferase reporter. The p53-regulated transcriptional activation of TRAPs and their negative controls at 24 h in BxPC-3 cells. The data is shown as means ± s.d. of n=2 independent experiments. **D.** The co-immunoprecipitation of TRAPs compared with the cotreatment of p53^Y220C^ binder (B-1 linker) and BRD4 binder (JQ1) in HEK293T cells after 4 h of compound treatment.

To test the idea that TRAPs induce p53-BRD4 binding in cells, we performed co-immunoprecipitation experiments. HEK293T cells were transfected with plasmids coding for FLAG-BRD4 and p53^Y220C^-V5, and anti-FLAG beads were used to isolate BRD4-associated proteins from cell lysates after cellular compound treatment. p53^Y220C^ pulled down with BRD4 after cell treatment with TRAPs; **TRAP-1** and **TRAP-2** induced stronger interactions than **TRAP-3** (**Fig. 3D**). Co-treatment with the two parental compounds did not induce an interaction (**Fig. 3D**). Taken together, these data demonstrate that TRAPs induce a stable interaction between p53^Y220C^ and BRD4.

### **TRAP-1** activates p53 target genes

We next evaluated the immediate transcriptional consequences of treatment with TRAPs. To do so, we examined validated direct transcriptional targets of p53 with known functions in p53 regulation (*MDM2*), cell cycle arrest (*CDKN1A*), and apoptosis (*PMAIP1*, *BBC3*, and *BAX)*^17,18,20^. We used reverse transcription quantitative PCR (RT-qPCR) to measure mRNA levels in BxPC-3 cells. Time course experiments after treatment with 3 µM of **TRAP-1** showed time-dependent upregulation of *MDM2* and *CDKN1A* as early as 2 h after incubation, with maximum levels accumulating at approximately 8 h post-treatment (**Fig. S4**). The apoptosis-effector genes *PMAIP1* and *BBC3* showed modest mRNA increases. *TP53* mRNA levels were not affected by the molecules (**Fig. S4**). To compare TRAPs with the parental compounds, we measured p53 target gene levels by RT-qPCR after 8 h of treatment. We tested **TRAP-1**, the parental binders, and the two negative controls in BxPC-3 cells. **TRAP-1** induced mRNA expression for *MDM2*, *CDKN1A*, and *BBC3* (**Fig. 4A**). Induction was stronger for **TRAP-1** compared with B-1, JQ1, or **TRAP-1-Neg1** and **TRAP-1-Neg2**. These data match the results from reporter gene assays and the kinetics of gene activation we have observed for the best TCIP compounds^28,29^. Overall, these results indicate that TRAPs induce robust expression of direct p53 target genes, likely by recruiting BRD4 to the sites bound by chemically corrected p53^Y220C^.

**Figure 4:**
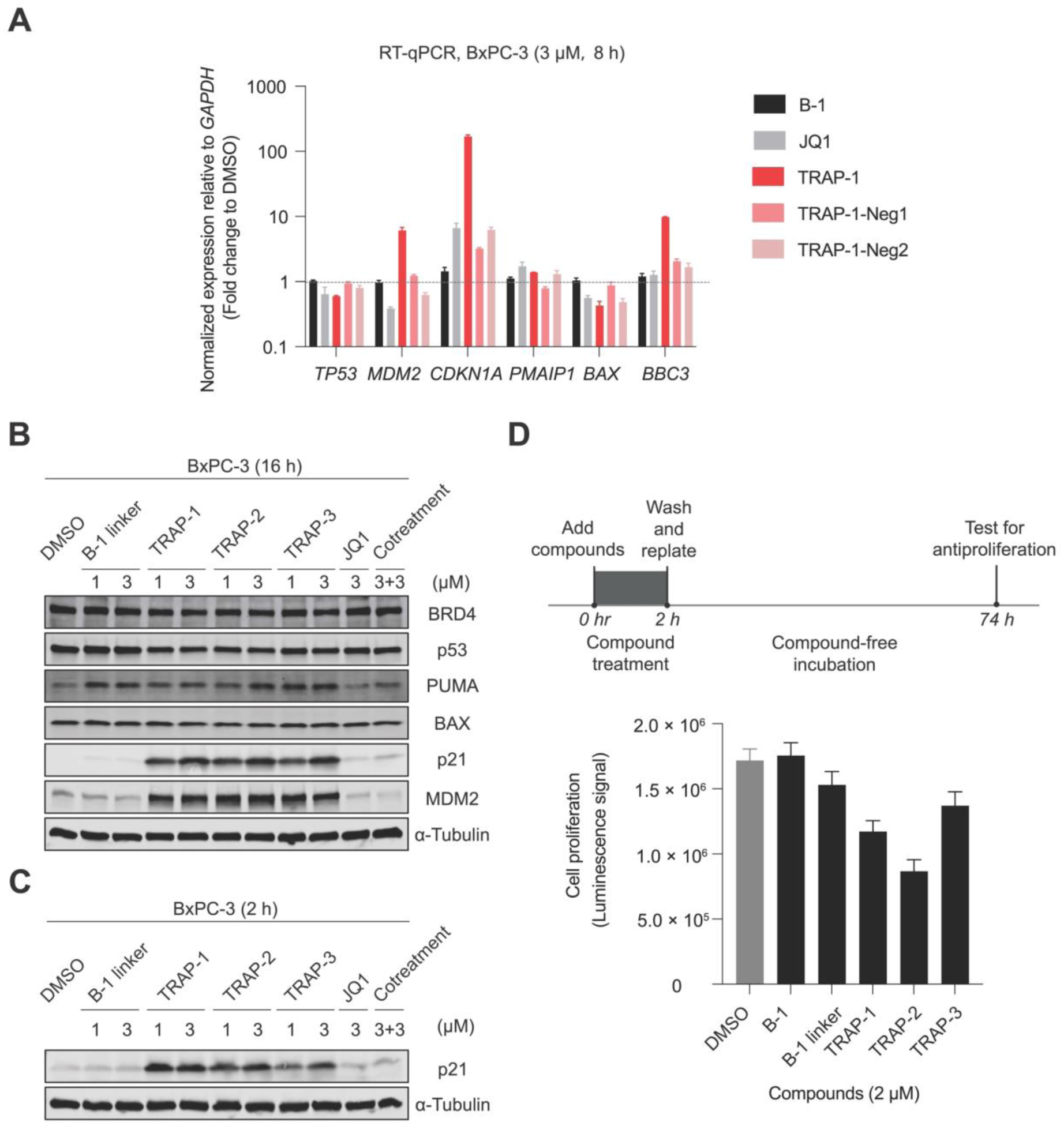
Activation of p53 target gene transcription. **A.** RT-qPCR. Changes of gene expression after 3 µM **TRAP-1** incubation for 8 h in BxPC-3 cells compared with the controls. A representative data is shown with mean ± s.d. of n=3 replicates. **B.** Changes in protein level after TRAPs treatment at 16 h in BxPC-3 cells compared with the controls. **C.** p21 protein level changes after TRAPs treatment at 2 h in BxPC-3 cells compared with the controls. **D.** Washout experiment. BxPC-3 cells were incubated with 2 µM TRAPs and controls for 2 h followed by a washout and measurement of cell proliferation at 74 h. A representative data is shown with mean ± s.d. of n=48 replicates.

We next used Western blotting to evaluate whether elevated mRNA levels for p53 target genes corresponds to elevated protein expression. Treatment with **TRAP-1**, **TRAP-2**, and **TRAP-3** for 16 h in BxPC-3 cells caused robust upregulation of p21 and MDM2 protein levels (**Fig. 4B**). The parental binders did not have this effect. We observed no significant changes in p53, PUMA, or BAX protein levels at this time point for any of the tested compounds. Potent upregulation of p21 was evident as early as 2 h after treatment with **TRAP-1**, highlighting the rapid and likely direct effect of the small molecules on p53 target genes (**Fig. 4C**).

Induction of cell cycle arrest factors, and p21 in particular, suggests that TRAPs may irreversibly reprogram cancer cell transcription. To test the idea that a pulsatile treatment with the most potent TRAPs might be sufficient to trigger an inescapable growth arrest, we conducted a washout experiment. BxPC-3 cells were treated with TRAPs for two hours – sufficient time to induce high levels of p21 protein – and the small molecules were subsequently washed away. After this, the cells were allowed to proliferate unperturbed for 72 h. Both **TRAP-1** and **TRAP-2** inhibited cell proliferation in this assay, while the parental B-1 binder did not (**Fig. 4D**). Collectively, these data demonstrate that the TRAPs can induce rapid and potent transcriptional activation in cells, and the washout result suggests that the effect can persist even after the compounds are removed.

### **TRAP-1** activity is p53 mutant-specific

Having explored the ability of the p53 transcriptional activator to upregulate cell-cycle arrest-related genes, we next investigated the antiproliferative effects of the compounds across a panel of cell lines with different p53 mutational statuses. Based on the biochemical p53 mutant selectivity of the TRAPs in the TR-FRET dimerization assay (**Fig. S3**), we compared the cell viability among BxPC-3 cells (p53^Y220C^), A549 cells (p53^WT^), and a non-tumorigenic colon epithelial cell line CCD 841 CoN (p53^WT^). Treatment with **TRAP-1**, **TRAP-2**, and **TRAP-3** for 72 h exhibited antiproliferative activity in BxPC-3 (p53^Y220C^) cells with submicromolar IC_50_ values, while compounds that induce weaker or no p53-BRD4 proximity had a weaker antiproliferative effects (**Fig. 5A**). **TRAP-1** exhibited 6.2-fold more potent antiproliferative activity in BxPC-3 cells than in A549 cells after a 3-day treatment (**Fig. 5B**; IC_50_ 3.94 µM A549/0.531 µM BxPC-3) and 22-fold more potent activity than in CCD 841 CoN cells after a 5-day treatment (**Fig. 5B**; IC_50_ 6.47 µM CCD 841 CoN/0.314 µM BxPC-3). We also observed modest antiproliferative activity with **TRAP-1** and **TRAP-1-Neg2** in A549 cells (**Fig. 5B**; IC_50_ 3.94 µM and 3.11 µM). **TRAP-1-Neg1**, which cannot bind and recruit BRD4, did not have antiproliferative activity in A549 cells (**Fig. 5B**; IC_50_ 7.48 µM). To further confirm the antiproliferation selectivity of **TRAP-1** in isogenic cells with different p53 statuses, we knocked out *TP53* in A549 cells and then reintroduced p53^WT^ or p53^Y220C^ to generate A549-p53^-/-^, A549-p53^WT^, and A549-p53^Y220C^ cell lines. While parental binders and binder co-treatment showed similar modest antiproliferative activities across the three cell lines, **TRAP-1** showed over 3-fold selectivity in A549-p53^Y220C^ cells (IC_50_ 1.03 µM) when compared with A549-p53^-/-^ (IC_50_ 3.83 µM) and A549-p53^WT^ (IC_50_ 3.45 µM) cells (**Fig. S5**). Taken together, these results suggest that **TRAP-1** induces a mutant-specific ternary complex formation, resulting in a potent and selective antiproliferative effect against p53^Y220C^ bearing cancer cells.

**Figure 5:**
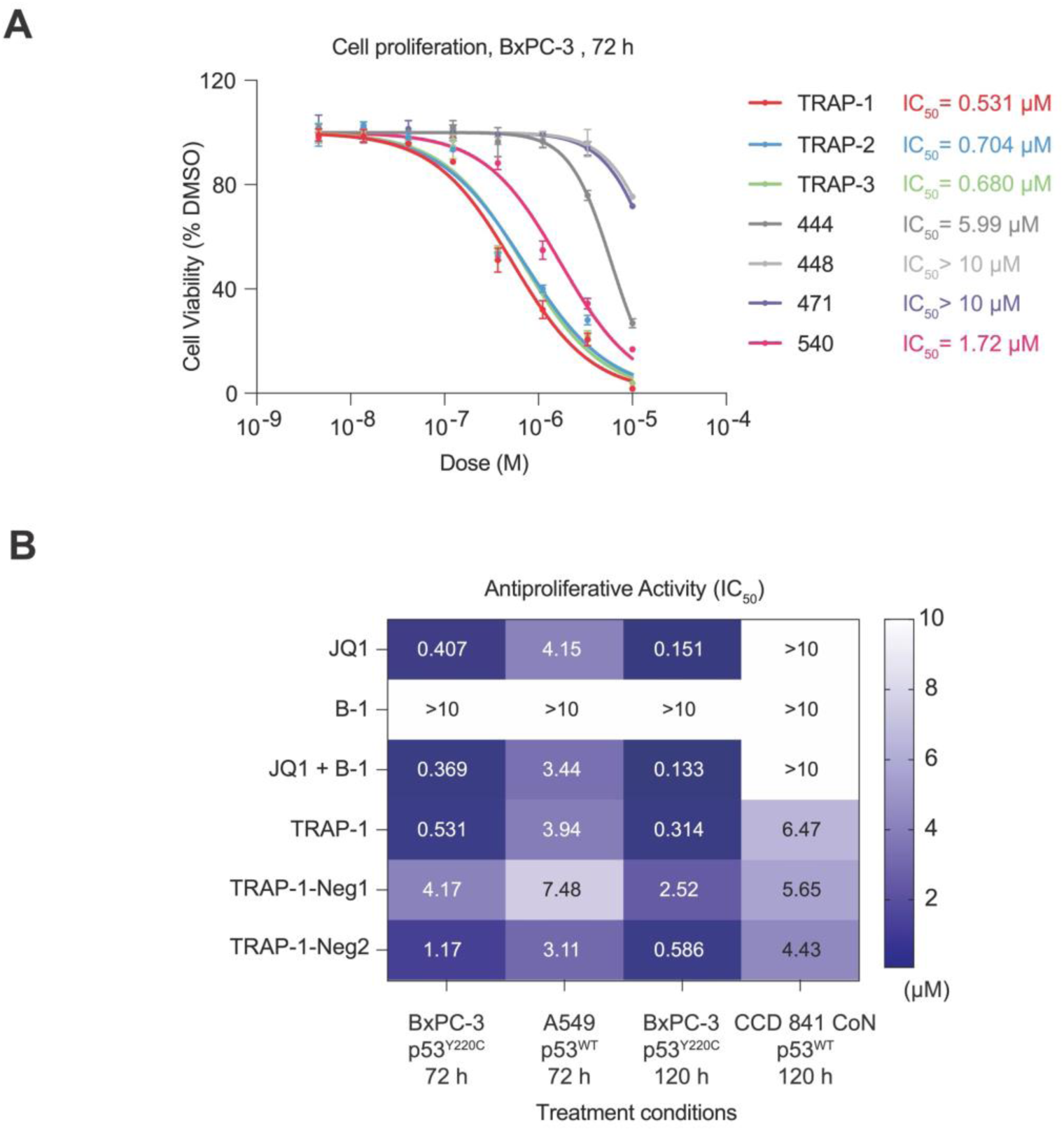
The antiproliferative activity of TRAPs in different cell lines. **A.** The antiproliferative activity of TRAPs compared with weaker or inactive compounds at 72 h in BxPC-3 cells. The data is shown as means ± s.d. of n=2 independent experiments. **B.** Heat map showing the antiproliferative potency of **TRAP-1** compared with controls in BxPC-3 (p53^Y220C^), A549 (p53^WT^), and CCD 841 CoN (p53^WT^). The data is shown as means of n=2 independent experiments.

## Discussion

Transcriptional dysregulation is a hallmark of cancer^35^. Overactivity of oncogenic transcription factors and loss of tumor suppressor transcription factors both enable cancer cells to escape normal control of cell proliferation^36^. While there have been successful approaches to inhibiting oncogenic transcription factors^37^, there have not been actionable strategies for activating tumor suppressor transcription factors like p53. Proof of concept experiments carried out in mouse models of disease demonstrate that p53 reactivation is a viable strategy to treat cancer^38^. Until now, chemical activators of mutant p53 have yet to deliver on the promise of this concept.

Here we developed a small molecule approach to generate p53 transcriptional activators. To do this, we chemically fused the transcriptional co-activator function of BRD4 to the restored DNA binding activity of p53^Y220C^. The resulting lead compound **TRAP-1** induced an interaction between p53^Y220C^ and BRD4. The ternary complex was p53 mutant-specific and activated p53 target gene transcription rapidly, potently, and dose-dependently in p53^Y220C^ bearing BxPC-3 cells. The transcriptional activation was significantly stronger than that observed upon treatment with parental p53^Y220C^ corrector, B-1, either given alone or in combination with the BRD4 binder JQ1.

Our experiments indicate that **TRAP-1** activates p53 by an induced proximity mechanism. Previous studies describe a synergistic relationship between BRD4 (with CPI203) and MDM2 inhibition (with nutlin-3). This was observed in acute myeloid leukemia cells^39^. Our biochemical and cellular data indicate that transcriptional activation induced by **TRAP-1** is fundamentally distinct at a mechanistic level. **TRAP-1** induced both p21 and MDM2, produced a modest increase in apoptosis-related genes at mRNA levels, and displayed antiproliferative activity in BxPC-3 cells. These and other activities correlated well with biochemical ternary complex formation and were obviated by minimal chemical modifications of **TRAP-1** to produce negative control compounds. Further efforts to find TRAP candidates using a different p53^Y220C^ ligand, PK9328, which was inactive in our p53 reporter assay, failed to produce active compounds, suggesting the activities of **TRAP-1** are not simply due to co-inhibition of BRD4 and p53, but rather reflect formation of a specific ternary complex.

This work demonstrates a new approach to activating p53 through the formation of a complex with BRD4 via chemically induced proximity. Further optimization is needed to enhance potency and mitigate cell cytotoxicity from BRD4 inhibition. Efforts to use different effector proteins are also currently underway. Besides BRD4, chromatin-modifying enzymes such as bromodomain and PHD finger containing 1 (BRPF1), histone acetyltransferase paralogues p300 and CREB-binding protein (CBP), and others have been recruited for endogenous gene activation^40,41^. Our group has also demonstrated the chemically induced proximity with transcriptional kinase CDK9 as a gain-of-function strategy to activate apoptosis^29^. In addition to these effectors, disease-specific driver proteins are potential candidates for the development of advanced TRAP molecules with improved selectivity.

Previous studies on bifunctional compounds that target p53 mainly focused on p53 post-translational modification^24,25^, protein abundance^23,26^, and cellular enrichment^27^. Direct potentiation of p53 transcriptional activity is heretofore unreported in the published literature. **TRAP-1** embodies an approach to drugging p53 that depends on protein-protein interaction induction by bivalent or monomeric molecular glue compounds. This strategy demonstrates the potential of chemically induced proximity to directly activate transcription factor function. While our study focuses on p53^Y220C^ specifically, this approach provides a potential solution to the challenge of developing activators for tumor suppressors that are dysregulated in cancer cells.

---

**Figure S1:**
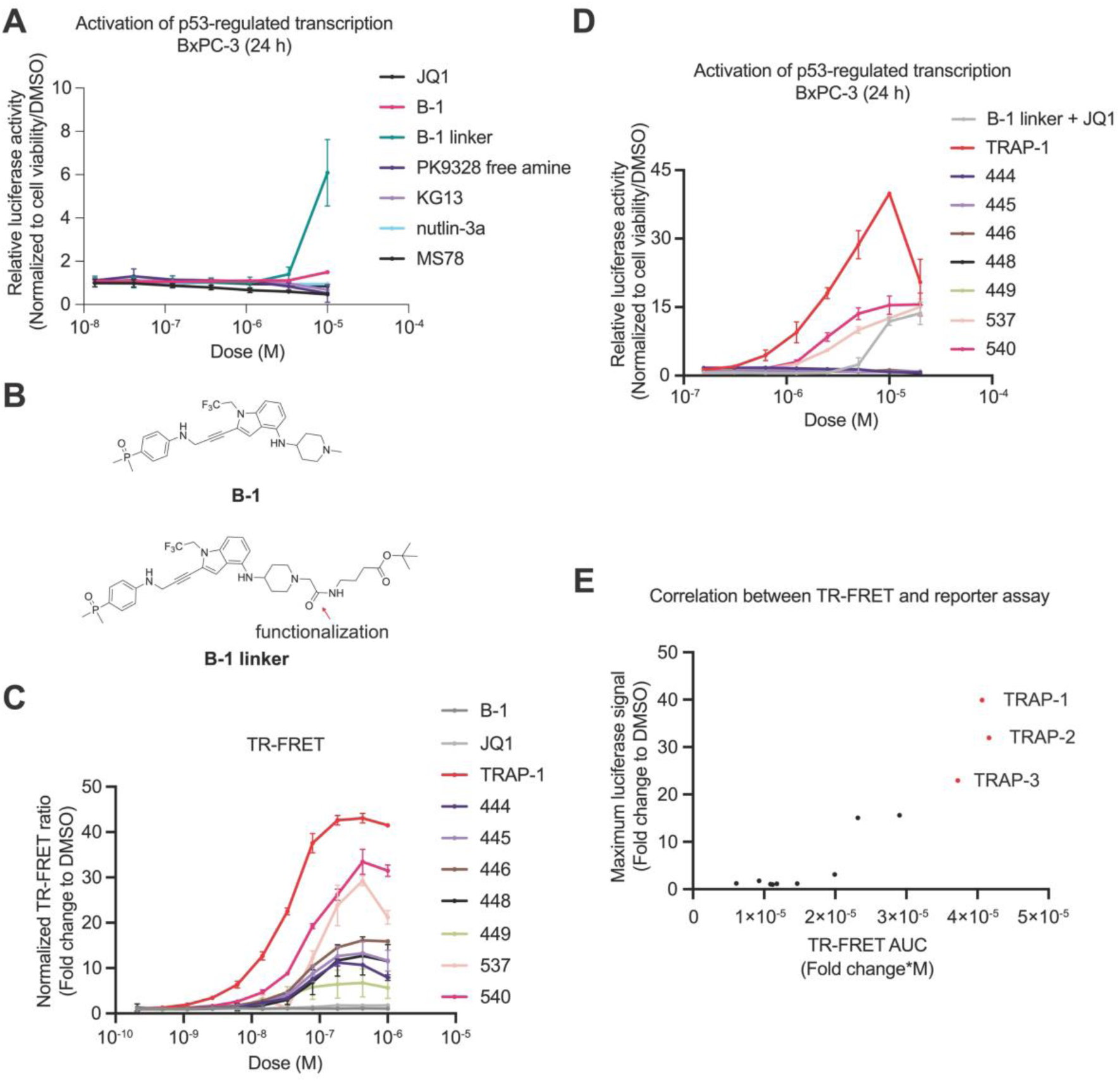
Development of B-1-based TRAPs. **A.** Activation of p53-regulated transcription in BxPC-3 cells with published p53^Y220C^ “corrector” (B-1, PK9328 free amine, KG13), p300/CBP-p53^Y220C^ acetylation-inducing chimera (MS78), MDM2 inhibitor (nutlin-3a), and BRD4 binder (JQ1) at 24 h. The data is shown as means ± s.d. of n=2 independent experiments. **B.** The structure of **B-1 linker** to validate the exit vector strategy. **C.** The ternary complex formation between p53^Y220C^ and BRD4_BD1_ mediated by weakly active or not active B-1-based compounds. The data is shown as means ± s.d. of n=2 independent experiments. **D.** Activation of p53-regulated transcription in BxPC-3 cells with weakly active or no active B-1-based compounds at 24 h. The data is shown as means ± s.d. of n=2 independent experiments. **E.** Scatter plot showing the relationship between the maximum luciferase signal and the TR-FRET area under the curve (AUC) of the B-1-based TRAPs.

**Figure S2:**
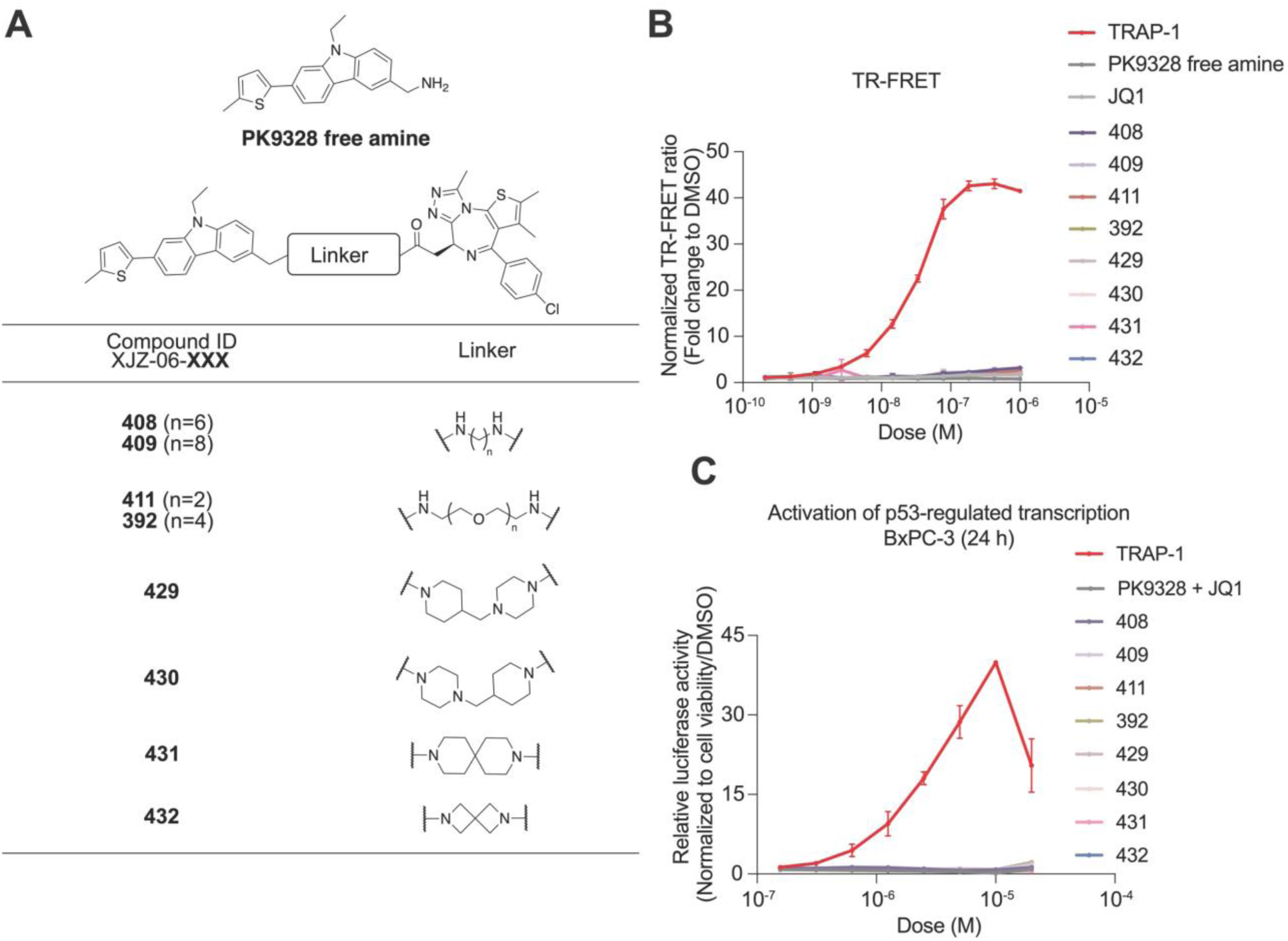
Development of PK9328-based TRAP candidates. **A.** The structure of the PK9328-based molecule library. **B.** The ternary complex formation measured by TR-FRET between purified p53^Y220C^ and BRD4_BD1_. The data is shown as means ± s.d. of n=2 independent experiments. **C.** Activation of p53-regulated transcription in BxPC-3 cells after compound treatment for 24 h. The data is shown as means ± s.d. of n=2 independent experiments.

**Figure S3:**
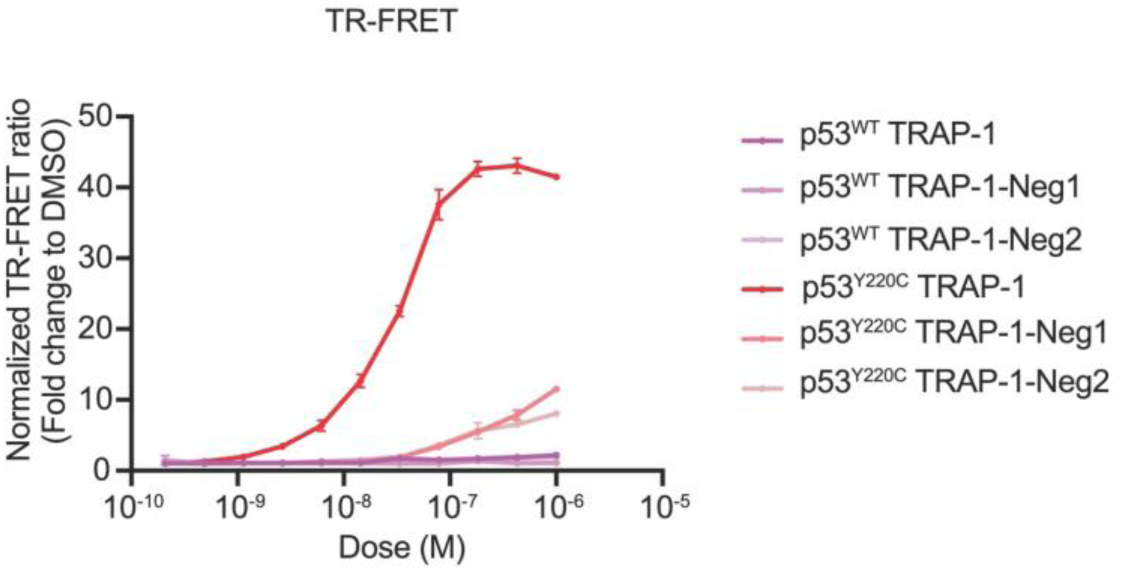
Selectivity of TRAP-1 on p53^Y220C^ over p53^WT^. The ternary complex formation of **TRAP-1** treatment measured by TR-FRET between purified p53^Y220C^ or p53^WT^ DNA binding domain and BRD4_BD1_. The data is shown as means ± s.d. of n=2 independent experiments.

**Figure S4:**
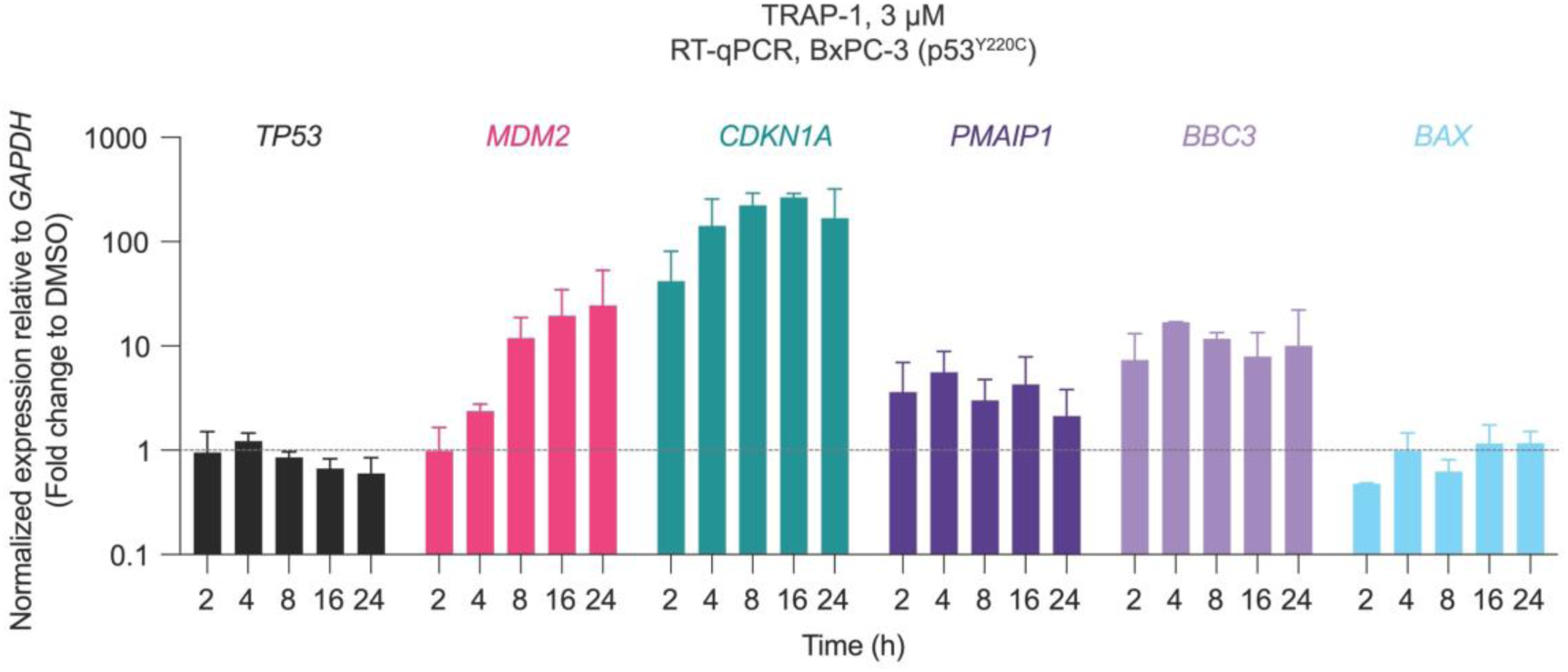
p53 target gene expression after TRAP-1 treatment. Gene expression of 3 µM **TRAP-1** in BxPC-3 cells after 2, 4, 8, 16, and 24 h treatment. The data is shown as means ± s.d. of n=2 independent experiments.

**Figure S5:**
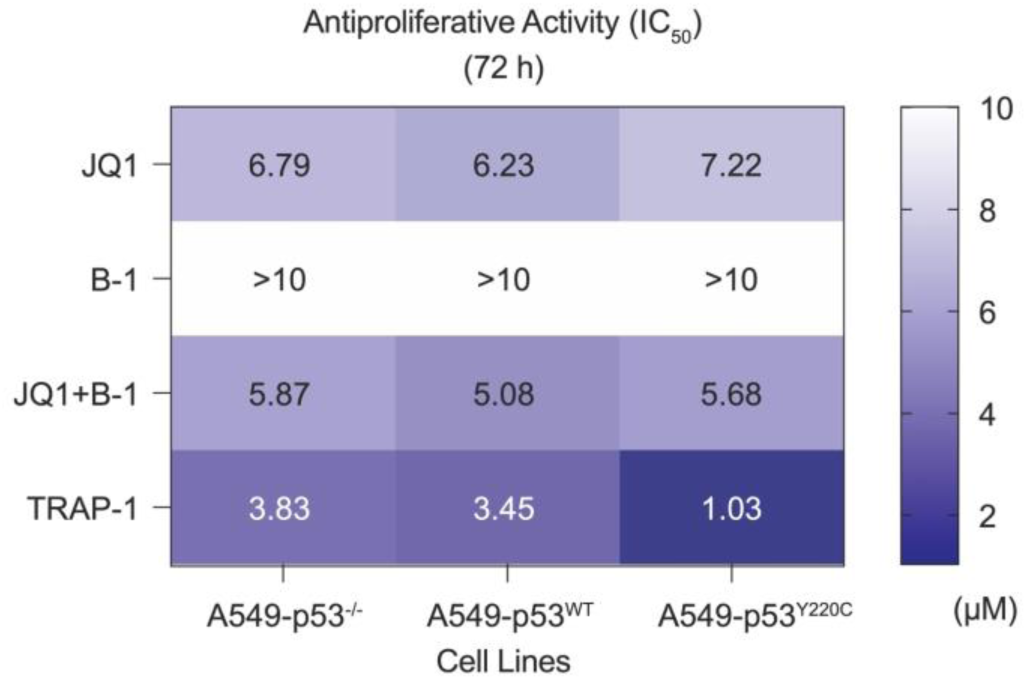
The antiproliferative activity of TRAP-1 and control compounds in isogenic A549 cell lines. Heat map showing the antiproliferative activity of **TRAP-1** compared with controls in A549-p53^-/-^, A549-p53^WT^, and A549-p53^Y220C^. The data is shown as means of n=2 independent experiments.

## Methods

### Chemical Synthesis

Additional details are provided in the supporting information.

### Cell Culture

Human pancreatic cancer (BxPC-3), kidney epithelial (HEK293T), lung cancer (A549), and colon epithelial (CCD 841 CoN) cell lines were obtained from the American Type Culture Collection (ATCC, Manassas, VA, USA). Cells were cultured in medium (RPMI 1640 for BxPC-3 cells; DMEM medium for HEK293T, A549 cells; EMEM for CCD 841 CoN) supplemented with 10% heat-inactivated FBS, 100 units/mL penicillin, 100 µg/mL streptomycin, and 0.25 µg/mL amphotericin B. Cells were incubated at 37°C with 5% CO_2_ in a humidified atmosphere. Mycoplasma testing was performed monthly using the MycoAlert mycoplasma detection kit (Lonza, Basel, Switzerland) and all lines were negative.

### p53^Y220C^ Reporter Assay

p53 luciferase reporter lentivirus (BPS Bioscience, San Diego, CA, USA) was used to transduce BxPC-3 cells cultured as described above, and tranductants were selected by growth in 1 µg/mL puromycin (Gibco Invitrogen Corp., Grand Island, NY, USA) added directly to the culture medium. Briefly, cells were seeded in 384-well plates (Corning, Corning, NY, USA, cat. no. 3570) and incubated overnight. Subsequently, the cells were treated with the indicated concentrations of compounds. After 24 h, the plates were subjected to Bright-Glo Luciferase Assay System (Promega, Madison, WI, USA) as described in the manufacturer’s manual. The proliferation assays were performed in biological triplicate.

### Cell Viability Assay (CellTiter-Glo Assay)

Cell viability was evaluated using the CellTiter-Glo assay (Promega). Briefly, cells were seeded in 384-well plates and incubated overnight. Subsequently, the cells were treated with the indicated concentrations of compounds. After 72 or 120 h, the plates were subjected to CellTiter-Glo as described in the manufacturer’s manual. The proliferation assays were performed in biological triplicate. IC_50_ values were determined using a non-linear regression curve fit in GraphPad Prism v10.3.0.

### Western Blotting Analysis

Total cell lysates were prepared in 2× sample loading buffer (i.e., 250 mM Tris-hydrochloride: pH 6.8, 4% sodium dodecyl sulfate, 10% glycerol, 0.006% bromophenol blue, 2% β-mercaptoethanol, 50 mM sodium fluoride, and 5 mM sodium orthovanadate). The samples with cell lysates were boiled for 5-8 min at 95°C. The protein concentrations of the cell lysates were quantified using the BCA method and a BCA Protein Assay Kit (Thermo Fisher Scientific, Waltham, MA, USA). Equal amounts of protein were subjected to 4-20% sodium dodecyl sulfate-polyacrylamide gel electrophoresis and transferred to polyvinylidene fluoride membranes (Millipore, Bedford, MA, USA) activated with 100% methanol. The membranes were blocked using Intercept (TBS) Blocking Buffer (LI-COR Biosciences, Lincoln, NE, USA), and subsequently probed with appropriate primary antibodies at 4°C overnight and then incubated with IRDye 800-labeled goat anti-rabbit IgG (LI-COR Biosciences, cat. no. 926-32211) or IRDye 680RD goat anti-Mouse IgG (LI-COR Biosciences, cat. no. 926-68070) secondary antibodies at room temperature for 1 h. After washing the membranes with PBS for 30 min, the membranes were detected on the Li-COR Odyssey CLx system.

### Co-Immunoprecipitation

HEK293T cells were seeded into a 6-well plate (8 × 10^5^ cells/well), cultured overnight, and transfected with 1.5 μg FLAG-tagged full-length BRD4 and 1.5 μg V5-tagged p53^Y220C^ plasmids using TransIT-LT1 transfection reagents (Mirus Bio, Madison, WI, USA). The transfected cells were cultured for another 48 h and treated with either compound or DMSO for 4 h before collection. The cells were collected and lysed in Pierce IP Lysis Buffer (Thermo Fisher Scientific) with cOmplete Mini Protease Inhibitor Cocktail (Roche, Basel, Switzerland) for 30 min on ice and centrifuged for 30 min at 4°C to remove the insoluble fraction. For immunoprecipitation, 20 μL of pre-cleaned anti-FLAG M2 magnetic beads (Sigma-Aldrich, St. Louis, MO, USA) were added to the lysates. The beads–lysate mix was incubated at 4°C overnight on a rotator. Beads were magnetically removed and washed three times with PBS, and the FLAG-Tagged protein was competitively eluted using 3X FLAG Peptide (APExBIO Technology, Houston, TX, USA). Immunoblotting was carried out as previously described.

### Reverse Transcription Quantitative PCR (RT-qPCR)

Total RNA was extracted using the Ditrect-zol RNA MicroPrep kit (Zymo Research, Irvine, CA, USA, cat. no. R2062). 1 μg of total RNA was used for synthesizing complementary DNAs (cDNAs) using RevertAid Reverse transcriptase (Thermo Fisher Scientific, cat. no. K1622). The synthesized cDNAs were then diluted 5 times in nuclease-free water and stored at –80°C until used. The diluted cDNAs were used for quantitative PCR in triplicate using PowerUP SYBR green master mix (Thermo Fisher Scientific, cat. no. A25743) and a 7900HT Fast Real-Time PCR machine (Applied Biosystems, Waltham, MA, USA). Expression analysis was performed using specific primers to amplify each target gene (Table 1). The housekeeping gene *GAPDH* used as an internal control to normalize the variability in each sample. All RT-qPCR reactions were conducted at 50°C for 2 min, 95°C for 2 min, and then 40 cycles of 95°C for 15 sec and 60°C for 1 min. Expression level of target genes were quantified using a standard curve.

**Table 1:**
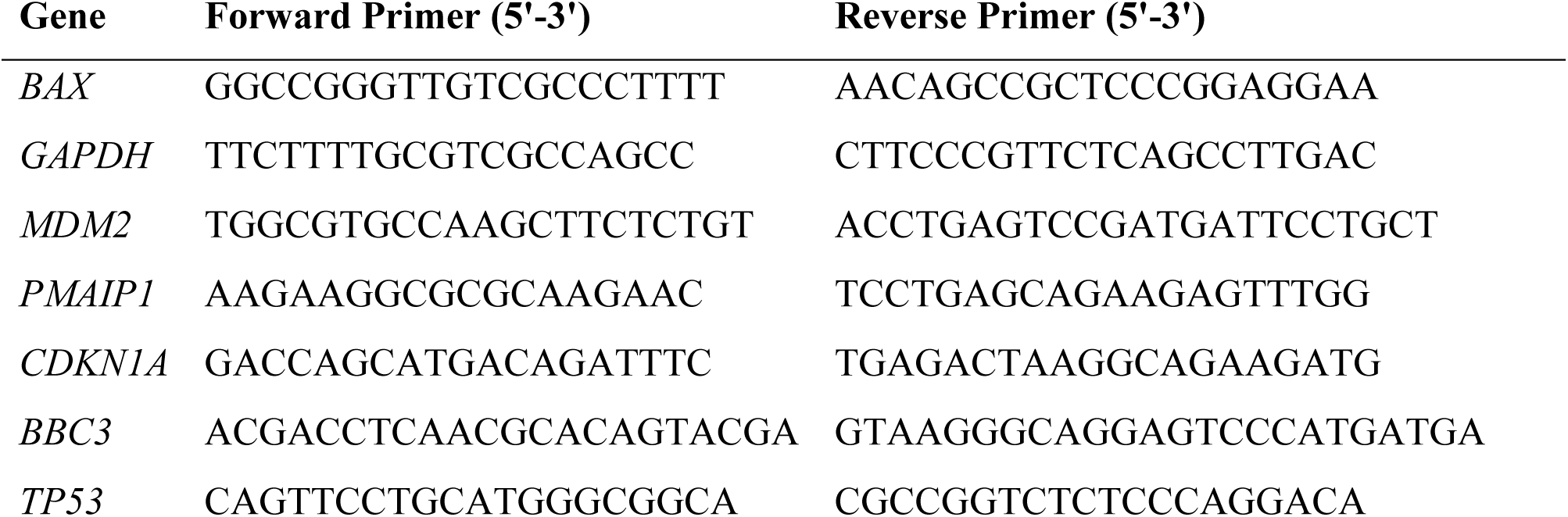
RT-qPCR primers.

### Expression and Purification of Human p53^WT^ and p53^Y220C^

Human p53^WT^ and p53^Y220C^ (residues 94–312) were expressed as 6X His-Spy-TEV fusion proteins in *Escherichia coli* Rosetta DE3 cells. Cells were cultured in Terrific Broth (TB) media supplemented with kanamycin (100 μg/mL) at 37°C with shaking at 130 rpm. Bacterial cultures were grown to an OD_600_ of 1.2 and induced with 1 mM isopropyl β-D-1-thiogalactopyranoside (IPTG) supplemented with 200 μM ZnSO_4_ at 18°C overnight. Cells were harvested by centrifugation at 4000 × g for 20 min at 4°C, and the cell pellet was resuspended 5 times w/v in lysis buffer containing 50 mM TRIS pH 8, 200 mM NaCl, 10% glycerol, 1 mM PMSF, and 2.5 mM DTT. After resuspension, cells were lysed by sonication (40% amplitude, 5 sec pulse, 5 sec pause, 5 min total, two times). The lysate was clarified by centrifugation at 75.000 × g for 40 min at 10°C and was then loaded onto a nickel-nitrilotriacetic acid (Ni-NTA) agarose column. Following incubation, the beads were washed with wash buffer containing 50 mM TRIS pH 8, 200 mM NaCl, 20 mM imidazole, and proteins were eluted from the resin in elution buffer containing 50 mM TRIS pH 8, 200 mM NaCl, 500 mM imidazole. The protein was then diluted in 50 mM HEPES pH 7 to lower salt concentration below 50 mM NaCl before being subjected to cation exchange chromatography and eluted in increased salt concentration buffer (50 mM HEPES, 50-1000 mM NaCl, 1 mM TCEP, pH 7). Eluted protein fractions were pooled and concentrated using an Amicon Ultra centrifugal filter (Molecular weight cut-off (MWCO) 10 kDa, Millipore). The concentrated protein was further purified by size-exclusion chromatography using a Superdex 75 16/600 GL column (Cytiva, Marlborough, MA, USA) in buffer containing 50 mM HEPES pH 7, 200 mM NaCl, and 1 mM TCEP. Protein-containing fractions were pooled, and the purity of the protein was assessed by SDS-PAGE. Protein concentration was determined using a Nanodrop. The purified protein was aliquoted, flash-frozen in liquid nitrogen, and stored at -80°C until further use.

### Expression and Purification of SpyCatcher S50C

The SpyCatcher S50C expression and purification were performed as previously described^42^. Briefly, SpyCatcher S50C was expressed in *Escherichia coli* Rosetta DE3. Cells were cultured in Luria-Bertani (LB) media supplemented with kanamycin and chloramphenicol (100 μg/mL) at 37°C with shaking at 130 rpm. Bacterial cultures were grown to an OD_600_ of 0.87 and induced with 1 mM IPTG at 18°C overnight. Cells were harvested by centrifugation at 4000 × g for 20 min at 4°C and lysed in the presence of 50 mM Tris-HCl pH 8.0, 200 mM NaCl, 1 mM TCEP, and 1 mM PMSF. Following ultracentrifugation, the soluble fraction was passed over the Ni-NTA agarose column and eluted with elution buffer containing 50 mM Tris-HCl pH 8.0, 200 mM NaCl, 400 mM imidazole, 1 mM TCEP. The affinity-purified protein was subjected to size exclusion chromatography (SEC), using a Superdex 75 16/600 GL column (Cytiva) equilibrated with SEC buffer (50 mM HEPES pH 7.5, 200 mM NaCl, 0.1 mM TCEP). Eluted protein was concentrated and flash-frozen in liquid nitrogen.

### Labeling of SpyCatcher S50C with BODIPY-FL-maleimide

Labeling of SpyCatcher with BODIPY-FL-maleimide was performed as previously described^42^. Briefly, purified SpyCatcher S50C protein was incubated with DTT (8 mM) at 4°C for 1 h. DTT was removed using a Superdex 75 16/600 GL size exclusion column in a buffer containing 50 mM TRIS pH 7.5, 150 mM NaCl, and 0.1 mM TCEP. BODIPY-FL-maleimide (Thermo Fisher) was dissolved in 100% DMSO and mixed with SpyCatcher S50C to achieve a 1.1 molar excess of BODIPY-FL-maleimide. SpyCatcher S50C labeling was carried out overnight at 4°C. The labeled SpyCatcher S50C was then purified on a Superdex 75 16/600 GL size exclusion column (Cytiva) in buffer containing 50 mM Tris pH 7.5, 150 mM NaCl, and 1 mM TCEP. The purified protein was concentrated using an Amicon Ultra centrifugal filter (Millipore), flash-frozen in liquid nitrogen, and stored at -80°C.

### Labeling of p53^WT^ and p53^Y220C^ with BODIPY-FL-SpyCatcher S50C

Purified p53^WT^ or p53^Y220C^ mutant proteins were incubated overnight at 4°C with BODIPY-FL labeled SpyCatcher S50C protein at a stoichiometric ratio. Protein was concentrated and loaded on the Superdex 75 10/300 size exclusion column (Cytiva). Labeling was monitored with absorption at 280 and 490 nm. The protein peak corresponding to the labeled protein was pooled, concentrated using an Amicon Ultra centrifugal filter (Millipore), flash-frozen (7.9 μM for p53-WT-SpyCatcher-S50C-BODIPY-FL and 2.4 μM for p53-Y220C-SpyCatcher-S50C-BODIPY-FL) in liquid nitrogen, and stored at -80°C.

### Expression, Purification and Biotinylation of BRD4_BD1_

BRD4_BD1_ was expressed in *Escherichia coli* Rosetta DE3. Cells were cultured in Luria-Bertani (LB) media supplemented with kanamycin and chloramphenicol (100 μg/mL) at 37°C with shaking at 130 rpm. Bacterial cultures were grown to an OD_600_ of 0.6 and induced with 1 mM IPTG at 18°C overnight. Cells were harvested by centrifugation at 4000 × g for 20 min at 4°C and lysed in the presence of 50 mM Tris-HCl pH 8.0, 200 mM NaCl, 10 mM imidazole, 1 mM TCEP, 1 mM PMSF, and 0.1% TritonX-100. Following ultracentrifugation, the soluble fraction was passed over the Ni-NTA agarose column and eluted with elution buffer containing 50 mM Tris-HCl pH 8.0, 200 mM NaCl, and 100 mM imidazole. The BRD4_BD1_ protein was biotinylated in vitro in the presence of 500 nM BirA enzyme, 10 mM MgCl_2_, 200 µM biotin, and 20 mM ATP in an SEC buffer containing 50 mM HEPES pH 7.5, 200 mM NaCl, and 1 mM TCEP for 1 h at room temperature, following incubation overnight at 4°C. The biotinylated protein was subjected to size exclusion chromatography, using a Superdex 75 16/600 GL column (Cytiva) equilibrated with the SEC buffer. Eluted protein was concentrated and flash-frozen in liquid nitrogen.

### Time-Resolved Fluorescence Resonance Energy Transfer (TR-FRET)

Compounds in dimerization assays were dispensed in a 384-well microplate (Corning, cat. no. 4514) using D300e Digital Dispenser (Tecan) normalized to 1% DMSO into 100 nM biotinylated BRD4_BD1_, 100 nM p53-Y220C-BODIPY-FL-SpyCatcher S50C (or 100 nM p53-WT-BODIPY-FL-SpyCatcher S50C) and 2 nM terbium-coupled streptavidin (Invitrogen) in a buffer containing 50 mM Tris pH 7.5, 200 mM NaCl, and 1 mM TCEP. Before the TR-FRET measurements were conducted, the reactions were incubated for 15 min at RT. The plate was then read on a Pherastar FSX (BMG Labtech) microplate reader. Following the excitation of terbium fluorescence at 337 nm, emission at 490 nm (terbium) and 520 nm (BODIPY-FL) were recorded and the TR-FRET signal of each data point was calculated as the 520/490 nm ratio. Data from two independent measurements (n=2), each calculated as an average of ten technical replicates per well per experiment, were plotted in GraphPad Prism v10.2.2.

### Construction of px459 TP53 Vector

pX459-TP53 vector was generated by cloning sgRNA targeting human *TP53* into the pX459 (Addgene #62988). *TP53*-targeted guide RNA was designed using the designing tool at https://chopchop.cbu.uib.no/. The backbone pX459 plasmid was cut with BbsI digestion enzyme and ligated to complementary annealed oligos containing the BbsI overhang site and *TP53* sgRNA sequence. The oligo sequences which targeting *TP53* were: 5’-CACCGCGACGCTAGGATCTGACTG-3’ and AAACCAGTCAGATCCTAGCGTCGC-3’.

### Generation of A549 TP53 Knockout (A549-p53^-/-^) Cell Line

A549 cells were transfected with pX459-*TP53* using lipofectamine 2000 (Invitrogen, cat. no. 11668027) according to manufacturer’s instructions. After 48 h of transfection, the transfected cells were passaged and cultured in medium containing 2 µg/mL puromycin (Gibco, cat. no. A11138-03) for three days. The surviving cells were grown for an additional 10 days to form cell colonies. Individual cell colonies were picked and expanded into different clones. Knockout efficiency of p53 in each clone was confirmed using immunoblotting with anti-p53 antibody.

### Construction of pLenti6/V5-p53_p53^Y220C^

The pLenti6/V5-p53_p53^WT^ (Addgene #22945) as a backbone plasmid was used as a template for site-directed mutagenesis to generate pLenti6/V5-p53_p53Y220C. KOD Xtreme Hot Start DNA Polymerase (Sigma-Aldrich, cat. no. 71975-M) was used to amplify the template with a pair of primers (Forward primer 5’-AGTGTGGTGGGTCCTGTGAGCCGCCTGAGGTT-3’; and Reverse primer: 5’-AACCTCAGGCGGCTCACAGGGCACCACACACT-3’), and the mutation was verified by Sanger sequencing.

### Generation of A549-p53^WT^ and -p53^Y220C^ Cell Lines

HEK293T cells were transfected with either pLenti6/V5-p53_p53^WT^ (Addgene #22945) or pLenti6/V5-p53_p53^Y220C^, along with pMD2.G (Addgene #12259)/psPAX2 (Addgene #12260), using TransIT-LT1 transfection reagents (Mirus Bio) to produce lentivirus. After 48 h of transfection, the supernatant was collected, filtered, and added to A549-p53^-/-^ cells in the presence of 5 µg/mL polybrene (APExBIO, cat. No. K2701). Following 48 h of infection, the infected cells were passaged and cultured in medium containing 10 µg/mL blasticidin (Gibco, cat. no. A11139-03) for five days. The surviving cells were grown for an additional 7 days to form cell colonies. Individual cell colonies were picked and expanded into different clones. The expression p53 in each clone was confirmed using immunoblotting with anti-V5 antibody.

## Acknowledgements

This work was supported by the National Institutes of Health (NIH) grant NIH High End Instrumentation grant (1S10OD028697-01) (to N.S.G.), R35 grant CA197591 (to L.D.A.), departmental funds from Stanford Chemical and Systems Biology and Stanford Cancer Institute (to N.S.G.), Ludwig Institute, Stanford’s SPARK Translational Research Program, and Bio-X. W.S.B. is supported by Basic Science Research Program through the National Research Foundation of Korea (NRF) funded by the Ministry of Education (grant no. RS-2024-00410290). R.P.N. is a member of the excellence cluster ImmunoSensation2 funded by the Deutsche Forschungsgemeinschaft (DFG) under Germany’s Excellence Strategy (EXC2151– 390873048). R.C.S. acknowledges the Swiss National Science Foundation for a postdoctoral fellowship (SNF Mobility grant P500PN_206898). M.W. acknowledges the Tobacco Related Diseases Research Program for a postdoctoral fellowship award (TRDRP T31FT1610).

## Author contributions

X.Z. and W.S.B. contributed equally to this work. X.Z., W.S.B., and N.S.G. conceived and initiated the study; X.Z. designed and synthesized bivalent compounds with the help of J.Z., Q.G., R.C.S., and T.Z.; W.S.B. designed and conducted cell biological experiments with the help of K.T.N., T.Q., Z.J., and M.W.; D.E.P. carried out biochemical studies with the help of J.G.; N.S.G., R.P.N., and L.D.A. supervised the project; The manuscript was written by X.Z., W.S.B., S.M.H, and N.S.G. with input from all authors.

## Data availability

Data generated in this study are provided in the manuscript, supplementary information, and source data files. Additional data supporting the findings of this study are available from the corresponding author upon reasonable request.

