## Supporting Information for "Activating p53^Y220C^ with a Mutant-Specific Small Molecule"

TABLE OF CONTENTS

1. Chemical Syntheses
2. NMR Spectra
3. Uncropped Western Blots

### 1. Chemical Syntheses

General synthetic methods

Unless otherwise noted, all reagents were purchased from commercial suppliers and used without further purification. Reactions were monitored using a Waters Acquity UPLC/MS system (Waters PDA eλ Detector, QDa Detector, Sample manager - FL, Binary Solvent Manager) using Acquity UPLC BEH C18 column (2.1 x 50 mm, 1.7 µm particle size): solvent gradient = 85% A at 0 min, 1% A at 1.7 min; solvent A = 0.1% formic acid in water; solvent B = 0.1% formic acid in Acetonitrile; flow rate: 0.6 mL/min. Analytical thin layer chromatography (TLC) was performed on Merck silica gel 60 F254 TLC glass plates and analytes were visualized by fluorescence quenching (using 254 nm light). Purification of reaction products was carried out by flash column chromatography using CombiFlash Rf with Teledyne Isco RediSep normal-phase silica flash columns (4 g, 12 g, 24 g, 40 g or 80 g), with Teledyne RediSep Gold C18 reversed-phase column (5.5 g, 15.5 g, 30 g, or 50 g), with preparative RP-HPLC using Waters SunFire Prep C18 column (19 x 100 mm, 5 µm particle size) with a gradient of 10-90% methanol in water containing 0.035% trifluoroacetic acid (TFA) over 40 min (45 min run time) at a flow of 40 mL/min, or with preparative TLC (prep-TLC) on Merck silica gel 60 F254 TLC glass plates. Assayed compounds were isolated and tested as TFA salts with assayed compounds purities in all cases greater than 95%, as determined by reverse-phase UPLC analysis. NMR spectra were acquired on a 500 MHz Bruker Avance III spectrometer, operating at the denoted spectrometer frequency given in MHz for the specified nucleus. All experiments were acquired at 298.0 K with a calibrated Bruker Variable Temperature Controller unless otherwise noted. The chemical shifts are reported in parts per million (ppm) and coupling constants (J) are given in Hertz (Hz). 1H NMR spectra are reported with the solvent resonance as the reference unless noted otherwise ($d_6$-DMSO at 2.50 ppm, $CDCl_3$ at 7.26 ppm). Peaks are reported as (s = singlet, d = doublet, t = triplet, q = quartet, m = multiplet or unresolved, br = broad signal, coupling constant(s) in Hz, integration).

**Procedure A:** Synthesis of JQ1 amine linker

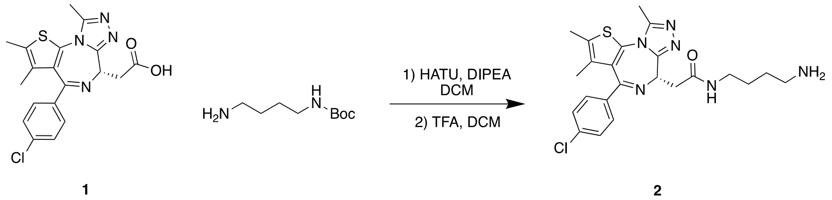


To a solution of (*S*)-2-(4-(4-chlorophenyl)-2,3,9-trimethyl-6*H*-thieno[3,2*f*][1,2,4]triazolo[4,3-*a*][1,4]diazepin-6-yl)acetic acid **1** (50 mg, 0.12 mmol, 1 eq.) and *tert*-butyl (4-aminobutyl)carbamate (23 mg, 0.12 mmol, 1 eq.) in DCM (1 mL) was added HATU (52 mg, 0.14 mmol, 1.1 eq.) and DIPEA (65 µL, 0.37 mmol, 3 eq.), and the reaction mixture was stirred at room temperature for 1 hour. The crude mixture was purified by flash chromatography with 0-10% MeOH/DCM to afford the coupling product as a colorless oil. The product was then dissolved in DCM (5 mL) and subjected to TFA (1 mL) at room temperature for 1 hour. The solvent was then evaporated to afford compound **2** as a yellow oil (55 mg, 94% in two steps) which was used directly in the next step. LC/MS(ESI) for $C_{23}H_{28}ClN_6OS$ $[M+H]^+$: m/z calcd, 471.17; found, 471.12.

**Procedure B:** Preparation of the 2-(4-((2-(3-((4-(dimethylphosphoryl)phenyl)amino)prop-1-yn-1-yl)-1-(2,2,2-trifluoroethyl)-1*H*-indol-4-yl)amino)piperidin-1-yl)acetic acid (Compound **10**)

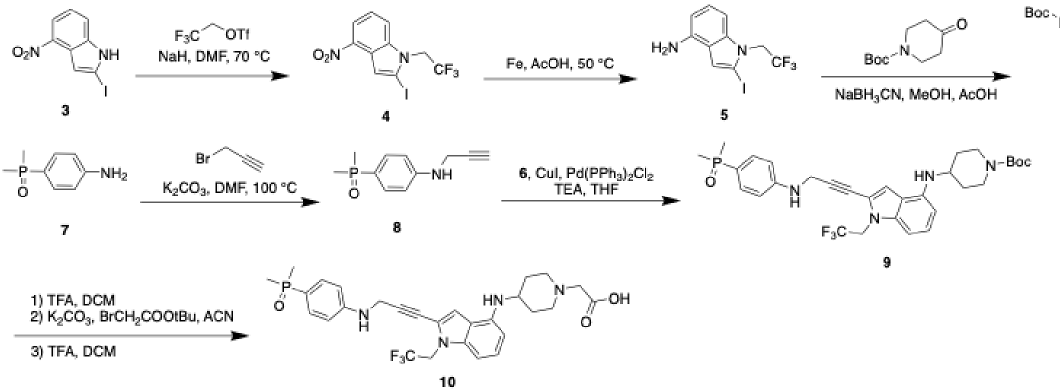


A solution of 2-Iodo-4-nitro-1H-indole **3** (403 mg, 1.74 mmol, 1 eq.) in DMF (3 mL) was added NaH (60% in oil, 208 mg, 5.21 mmol, 3 eq.) at 0 °C in portions. The solution was then stirred at the same temperature for 1 hour before adding 2,2,2-trifluoroethyl trifluoromethylsulfonate (750 µL, 5.22 mmol, 3 eq.). The temperature was then raised to 70 °C and the reaction was stirred for 2 hours. The reaction was quenched with water and extracted with ethyl acetate three times. The combined organic phase was washed with water, dried with anhydrous magnesium sulfate, and concentrated under reduced pressure. The crude residue was purified by flash chromatography column with EA/Hex to afford intermediate **4** as a brown solid (470 mg, 73.2%). $^1$H NMR (500 MHz, $CDCl_3$) δ 8.15 (dd, *J* = 8.0, 0.8 Hz, 1H), 7.71 – 7.63 (m, 2H), 7.32 (t, *J* = 8.1 Hz, 1H), 4.84 (q, *J* = 8.2 Hz, 2H).

Intermediate **4** (470 mg, 1.27 mmol, 1 eq.) and iron (426 mg, 7.26 mmol, 6 eq.) were dissolved in acetic acid (7 mL), and the reaction mixture was stirred for 4 hours at 50 °C. The mixture was then filtered, and the filtrate was concentrated under vacuum and purified by C18 chromatography column with 10-100% ACN/ $H_2O$ to afford intermediate **5** a brown solid (410 mg, 95%). LC/MS(ESI) for $C_{10}H_9F_3IN_2$ $[M+H]^+$: m/z calcd, 340.98; found, 340.96.

To a solution of intermediate **5** (400 mg, 1.18 mmol, 1 eq.) and tert-butyl 4-oxopiperidine-1-carboxylate (1.17 g, 5.88 mmol, 5 eq.) in ethanol (12 mL) was added titanium ethoxide (1.34 g, 5.88 mmol, 5eq), and the solution was stirred at 50 °C overnight. The mixture was then cooled to room temperature, and sodium cyanoborohydride (370 mg, 5.88 mmol, 5 eq.) was added. The resulting solution was stirred at room temperature overnight before purification through a flash chromatography column with EA/Hex to afford intermediate **6** (277 mg, 45%) as a pink solid. LC/MS(ESI) for $C_{20}H_{26}F_3IN_3O_2$ $[M+H]^+$: m/z calcd, 524.10; found, 524.04.

To a solution of (4-aminophenyl)dimethylphosphine oxide **7** (100 mg, 591 µmol, 1 eq.) and 3-bromoprop-1-yne (80% in toluene, 72.4 µL, 650 µmol, 1.1 eq.) in DMF (3 mL) was added potassium carbonate (163 mg, 1.18 mmol, 2 eq.), and the reaction mixture was stirred at 100 °C for 1 hour. The crude was then purified by C18 chromatography column with 10-100% ACN/$H_2O$ to afford intermediate **8** (65.6 mg, 53.5%) as a yellow oil. LC/MS(ESI) for $C_{11}H_{15}NOP$ $[M+H]^+$: m/z calcd, 208.09; found, 208.03.

A solution of intermediate **6** (277 mg, 529 µmol, 1 eq.), intermediate **8** (110 mg, 529 µmol, 1 eq.), bis(triphenylphosphine)palladium(II) dichloride (37.2 mg, 52.9 µmol, 0.1 eq.), and copper(I) iodide (5.04 mg, 26.5 µmol, 0.05 eq.) in THF (5 mL) under nitrogen was added triethylamine (5 mL), and the reaction was stirred at 50 °C overnight. The crude was then filtered through celite and purified by flash chromatography with MeOH/DCM to give the intermediate **9** as a brown solid (175.6 mg, 55.1%). LC/MS(ESI) for $C_{31}H_{39}F_3N_4O_3P$ $[M+H]^+$: m/z calcd, 603.27; found, 603.15.

The intermediate **9** (200 mg, 332 µmol, 1 eq.) was dissolved in DCM (5 mL) and subjected to TFA (1 mL), and the reaction mixture was stirred for 2 hours before concentrating under reduced pressure. The crude was then dissolved in ACN (3 mL) followed by the addition of potassium carbonate (138 mg, 997 µmol, 3 eq.) and tert-butyl 2-bromoacetate (146 µL, 997 µmol, 3 eq.). After stirring for 2 hours at room temperature, the reaction mixture was purified by C18 chromatography column with 10-100% ACN/$H_2O$ to afford the reaction product. The reaction product was then dissolved in DCM (5 mL) with TFA (1 mL), and the solution was stirred for 2 hours before concentrating under reduced pressure to afford compound **10** in crude (130 mg, 70% in three steps). LC/MS(ESI) for $C_{28}H_{33}F_3N_4O_3P$ $[M+H]^+$: m/z calcd, 561.22; found, 561.24.

Synthesis of *tert*-butyl 4-(2-(4-((2-(3-((4-(dimethylphosphoryl)phenyl)amino)prop-1-yn-1-yl)-1-(2,2,2-trifluoroethyl)-1*H*-indol-4-yl)amino)piperidin-1-yl)acetamido)butanoate (**B-1 linker**)

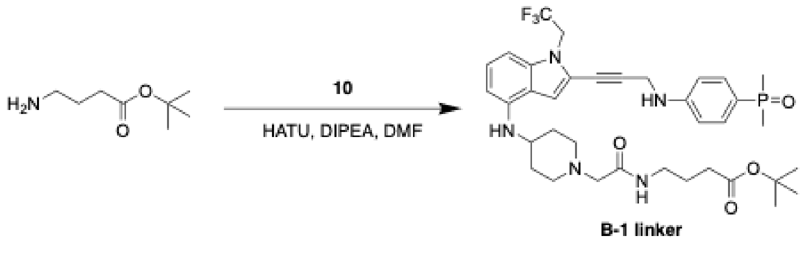

To a solution of *tert*-butyl 4-aminobutanoate (1.4 mg, 8.9 µmol, 1 eq.), compound **10** (5.0 mg, 8.9 µmol, 1 eq.), HATU (3.7 mg, 9.8 µmol, 1.1 eq.) in DMF (200 µL) was added DIPEA (6.2 µL, 45 µmol, 5 eq.), and the reaction was stirred at room temperature overnight. The crude was then purified by HPLC with MeOH/$H_2O$ (0.035% TFA) to give compound **B-1 linker** as a white solid (4.17 mg, 67%). LC/MS(ESI) for $C_{36}H_{48}F_3N_5O_4P$ $[M+H]^+$: m/z calcd, 702.34; found, 702.57. $^1$H NMR (500 MHz, d$^6$-DMSO) δ 9.75 (br, 1H), 8.56 (t, *J* = 5.7 Hz, 1H), 7.53 – 7.46 (m, 2H), 7.12 (d, *J* = 14.1 Hz, 1H), 7.03 (t, *J* = 8.0 Hz, 1H), 6.86 – 6.78 (m, 2H), 6.75 (d, *J* = 8.2 Hz, 1H), 6.22 (d, *J* = 7.8 Hz, 1H), 4.94 (q, *J* = 9.2 Hz, 2H), 4.29 (s, 2H), 3.96 – 3.86 (m, 2H), 3.65 – 3.56 (m, 1H), 3.52 (d, *J* = 11.7 Hz, 2H), 3.33 (s, 1H), 3.24 – 3.10 (m, 4H), 2.24 (t, *J* = 7.4 Hz, 2H), 2.15 (d, *J* = 13.8 Hz, 2H), 1.96 – 1.71 (m, 2H), 1.66 (p, *J* = 7.3 Hz, 2H), 1.55 (d, *J* = 13.1 Hz, 6H), 1.40 (s, 9H).

**<u>Procedure C:</u>** Synthesis of (*S*)-2-(4-(4-chlorophenyl)-2,3,9-trimethyl-6*H*-thieno[3,2-*f*][1,2,4]triazolo[4,3-*a*][1,4]diazepin-6-yl)-*N*-(4-(2-(4-((2-(3-((4-(dimethylphosphoryl)phenyl)amino)prop-1-yn-1-yl)-1-(2,2,2-trifluoroethyl)-1*H*-indol-4-yl)amino)piperidin-1-yl)acetamido)butyl)acetamide (**XJZ-06-444**)

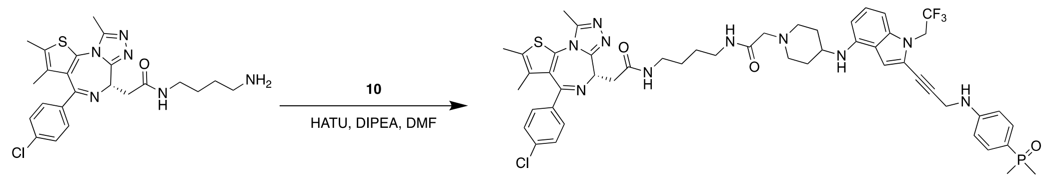


2

XJZ-06-444

To a solution of compound **2** (5.0 mg, 11 µmol, 1 eq.), compound **10** (6.0 mg, 11 µmol, 1 eq.), HATU (4.8 mg, 13 µmol, 1.2 eq.) in DMF (200 µL) was added DIPEA (7.4 µL, 53 µmol, 5 eq.), and the reaction was stirred at room temperature overnight. The crude was then purified by HPLC with MeOH/$H_2O$ (0.035% TFA) to give compound **XJZ-06-444** as a white solid (3.53 mg, 33%). LC/MS(ESI) for $C_{51}H_{58}ClF_3N_{10}O_3PS$ $[M+H]^+$: m/z calcd, 1013.38; found,1013.37. $^1$H NMR (500 MHz, d$^6$-DMSO) δ 9.76 (s, 1H), 8.55 (d, *J* = 5.9 Hz, 1H), 8.21 (t, *J* = 5.7 Hz, 1H), 7.49 (m, 4H), 7.45 – 7.39 (m, 2H), 7.17 – 7.07 (m, 1H), 7.02 (t, *J* = 8.0 Hz, 1H), 6.80 (dd, *J* = 8.6, 2.2 Hz, 2H), 6.74 (d, *J* = 8.3 Hz, 1H), 6.21 (d, *J* = 7.9 Hz, 1H), 4.94 (q, *J* = 9.4 Hz, 2H), 4.50 (t, *J* = 7.1 Hz, 1H), 4.29 (s, 2H), 3.96 – 3.87 (m, 2H), 3.63 – 3.55 (m, 1H), 3.55 – 3.49 (m, 2H), 3.32 (s, 1H), 3.32 – 3.07 (m, 8H), 2.59 (s, 3H), 2.40 (s, 3H), 2.17 – 2.10 (m, 2H), 1.83 – 1.74 (m, 2H), 1.62 (s, 3H), 1.55 (d, *J* = 13.1 Hz, 6H), 1.48 (br, 4H).

Synthesis of (*S*)-2-(4-(4-chlorophenyl)-2,3,9-trimethyl-6*H*-thieno[3,2-*f*][1,2,4]triazolo[4,3-*a*][1,4]diazepin-6-yl)-*N*-(6-(2-(4-((2-(3-((4-(dimethylphosphoryl)phenyl)amino)prop-1-yn-1-yl)-1-(2,2,2-trifluoroethyl)-1*H*-indol-4-yl)amino)piperidin-1-yl)acetamido)hexyl)acetamide (**XJZ-06-445**)

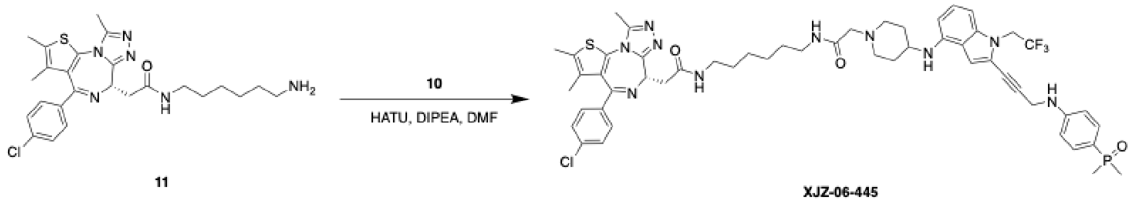


11

XJZ-06-445

The corresponding compound was prepared following **Procedure C** using intermediate **11** (5 mg, 10 µmol, 1 eq.) prepared through **Procedure A** (88% in two steps, LC/MS(ESI) for $C_{25}H_{32}ClN_6OS$ $[M+H]^+$: m/z calcd, 499.20; found, 499.13.). Purification by HPLC with $MeOH/H_2O$ (0.035% TFA) to afford the product as a white solid (4.23 mg, 41%). LC/MS(ESI) for $C_{53}H_{62}ClF_3N_{10}O_3PS$ $[M+H]^+$: m/z calcd, 1041.41; found, 1041.42 $[M+1]^+$. $^1H$ NMR (500 MHz, $d^6$-DMSO) δ 9.74 (s, 1H), 8.53 (t, *J* = 5.7 Hz, 1H), 8.18 (t, *J* = 5.7 Hz, 1H), 7.53 – 7.46 (m, 4H), 7.42 (d, *J* = 8.5 Hz, 2H), 7.12 (d, *J* = 11.7 Hz, 1H), 7.02 (t, *J* = 8.0 Hz, 1H), 6.80 (dd, *J* = 8.7, 2.2 Hz, 2H), 6.74 (d, *J* = 8.2 Hz, 1H), 6.21 (d, *J* = 7.9 Hz, 1H), 4.94 (q, *J* = 9.3 Hz, 2H), 4.50 (dd, *J* = 7.8, 6.4 Hz, 1H), 4.29 (s, 2H), 3.98 – 3.84 (m, 2H), 3.63 – 3.55 (m, 1H), 3.51 (d, *J* = 11.4 Hz, 2H), 3.32 (s, 1H), 3.29 – 3.08 (m, 8H), 2.59 (s, 3H), 2.40 (s, 3H), 2.20 – 2.06 (m, 2H), 1.84 – 1.66 (m, 2H), 1.61 (s, 3H), 1.56 (d, *J* = 13.1 Hz, 6H), 1.48 – 1.39 (m, 4H), 1.33 – 1.27 (m, 4H).

Synthesis of (S)-2-(4-(4-chlorophenyl)-2,3,9-trimethyl-6H-thieno[3,2-f][1,2,4]triazolo[4,3-a][1,4]diazepin-6-yl)-N-(8-(2-(4-((2-(3-((4-(dimethylphosphoryl)phenyl)amino)prop-1-yn-1-yl)-1-(2,2,2-trifluoroethyl)-1H-indol-4-yl)amino)piperidin-1-yl)acetamido)octyl)acetamide (**XJZ-06-446**)

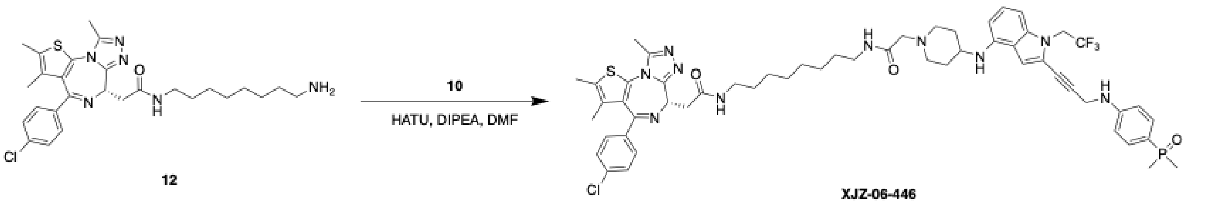


The corresponding compound was prepared following **Procedure C** using intermediate **12** (5 mg, 9.5 µmol, 1 eq.) prepared through **Procedure A** (91% in two steps, LC/MS(ESI) for $C_{27}H_{36}ClN_6OS$ $[M+H]^+$: m/z calcd, 527.24; found, 527.19.). Purification by HPLC with $MeOH/H_2O$ (0.035% TFA) to afford the product as a white solid (3.33 mg, 33%). LC/MS(ESI) for $C_{55}H_{66}ClF_3N_{10}O_3PS$ $[M+H]^+$: m/z calcd, 1069.44; found, 1069.53. $^1H$ NMR (500 MHz, $d^6$-DMSO) δ 9.73 (s, 1H), 8.53 (t, *J* = 5.5 Hz, 1H), 8.17 (t, *J* = 5.7 Hz, 1H), 7.54 – 7.46 (m, 4H), 7.45 – 7.41 (m, 2H), 7.13 (d, *J* = 12.5 Hz, 1H), 7.03 (t, *J* = 7.9 Hz, 1H), 6.81 (dd, *J* = 8.7, 2.2 Hz, 2H), 6.75 (d, *J* = 8.2 Hz, 1H), 6.22 (d, *J* = 7.9 Hz, 1H), 4.95 (q, *J* = 9.3 Hz, 2H), 4.51 (dd, *J* = 8.0, 6.1 Hz, 1H), 4.30 (s, 2H), 3.94 – 3.87 (m, 2H), 3.65 – 3.56 (m, 1H), 3.52 (d, *J* = 11.5 Hz, 2H), 3.33 (s, 1H), 3.29 – 3.05 (m, 8H), 2.60 (s, 3H), 2.41 (s, 3H), 2.22 – 2.09 (m, 2H), 1.83 – 1.68 (m, 2H), 1.63 (s, 3H), 1.56 (d, *J* = 13.1 Hz, 6H), 1.50 – 1.38 (m, 4H), 1.36 – 1.21 (m, 8H).

Synthesis of (S)-2-(4-(4-chlorophenyl)-2,3,9-trimethyl-6H-thieno[3,2-f][1,2,4]triazolo[4,3-a][1,4]diazepin-6-yl)-N-(2-(2-(2-(2-(4-((2-(3-((4-(dimethylphosphoryl)phenyl)amino)prop-1-yn-1-yl)-1-(2,2,2-trifluoroethyl)-1H-indol-4-yl)amino)piperidin-1-yl)acetamido)ethoxy)ethoxy)ethyl)acetamide (**XJZ-06-448**)

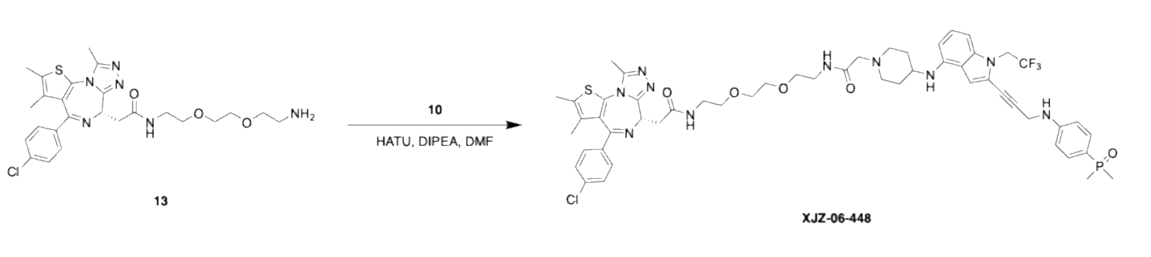

The corresponding compound was prepared following **Procedure C** using intermediate **13** (5 mg, 9.4 µmol, 1 eq.) prepared through **Procedure A** (91% in two steps, LC/MS(ESI) for $C_{25}H_{32}ClN_6O_3S$ $[M+H]^+$: m/z calcd, 531.19; found, 531.19.). Purification by HPLC with MeOH/$H_2O$ (0.035% TFA) to afford the product as a white solid (3.25 mg, 32%). LC/MS(ESI) for $C_{53}H_{62}ClF_3N_{10}O_5PS$ $[M+H]^+$: m/z calcd, 1073.40; found, 1073.44. $^1H$ NMR (500 MHz, $d^6$-DMSO) δ 9.75 (s, 1H), 8.66 (t, *J* = 5.7 Hz, 1H), 8.28 (t, *J* = 5.8 Hz, 1H), 7.54 – 7.46 (m, 4H), 7.43 (d, *J* = 8.3 Hz, 2H), 7.12 (d, *J* = 9.8 Hz, 1H), 7.02 (t, *J* = 8.1 Hz, 1H), 6.81 (dd, *J* = 8.6, 2.2 Hz, 2H), 6.75 (d, *J* = 8.2 Hz, 1H), 6.21 (d, *J* = 7.9 Hz, 1H), 4.95 (q, *J* = 9.2 Hz, 2H), 4.52 (dd, *J* = 7.9, 6.2 Hz, 1H), 4.29 (s, 2H), 3.96 – 3.90 (m, 2H), 3.63 – 3.53 (m, 5H), 3.53 – 3.44 (m, 5H), 3.38 – 3.13 (m, 10H), 2.59 (s, 3H), 2.40 (s, 3H), 2.14 (d, *J* = 14.2 Hz, 2H), 1.95 – 1.72 (m, 2H), 1.62 (s, 3H), 1.56 (d, *J* = 13.1 Hz, 6H).

Synthesis of (*S*)-2-(4-(4-chlorophenyl)-2,3,9-trimethyl-6*H*-thieno[3,2-*f*][1,2,4]triazolo[4,3-*a*][1,4]diazepin-6-yl)-*N*-(1-(4-((2-(3-((4-(dimethylphosphoryl)phenyl)amino)prop-1-yn-1-yl)-1-(2,2,2-trifluoroethyl)-1*H*-indol-4-yl)amino)piperidin-1-yl)-2-oxo-6,9,12,15-tetraoxa-3-azaheptadecan-17-yl)acetamide (**XJZ-06-449**)

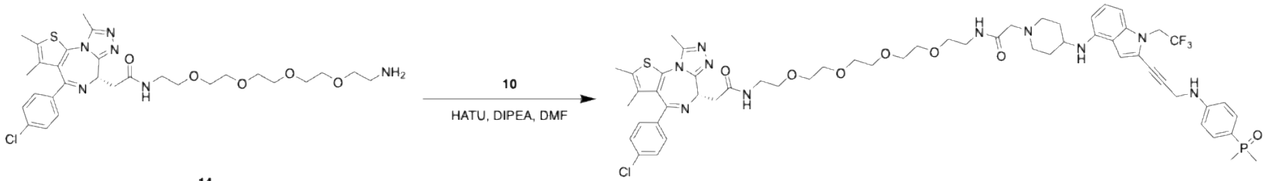


14

XJZ-06-449

The corresponding compound was prepared following **Procedure C** using intermediate **14** (5 mg, 8.1 µmol, 1 eq.) prepared through **Procedure A** (84% in two steps, LC/MS(ESI) for $C_{29}H_{40}ClN_6O_5S$ $[M+H]^+$: m/z calcd, 619.25; found, 619.30.). Purification by HPLC with MeOH/$H_2O$ (0.035% TFA) to afford the product as a white solid (1.45 mg, 15%). LC/MS(ESI) for $C_{57}H_{70}ClF_3N_{10}O_7PS$ $[M+H]^+$: m/z calcd, 1161.45; found,1161.33. $^1H$ NMR (500 MHz, $d^6$-DMSO) δ 9.78 (s, 1H), 8.73 – 8.57 (m, 1H), 8.31 – 8.26 (m, 1H), 7.54 – 7.45 (m, 4H), 7.42 (d, *J* = 8.5 Hz, 2H), 7.18 – 7.09 (m, 1H), 7.01 (t, *J* = 7.9 Hz, 1H), 6.83 – 6.77 (m, 2H), 6.74 (d, *J* = 8.3 Hz, 1H), 6.21 (d, *J* = 7.9 Hz, 1H), 4.93 (q, *J* = 9.0 Hz, 2H), 4.50 (dd, *J* = 8.0, 6.0 Hz, 1H), 4.29 (s, 2H), 3.99 – 3.88 (m, 2H), 3.55 – 3.45 (m, 18H), 3.35 – 3.15 (m, 10H), 2.59 (s, 3H), 2.40 (s, 3H), 2.20 – 2.08 (m, 2H), 1.82 – 1.74 (m, 2H), 1.61 (s, 3H), 1.55 (d, *J* = 13.0 Hz, 6H).

Synthesis of (S)-2-(4-(4-chlorophenyl)-2,3,9-trimethyl-6H-thieno[3,2-f][1,2,4]triazolo[4,3-a][1,4]diazepin-6-yl)-1-(4-((1-(2-(4-((2-(3-((4-(dimethylphosphoryl)phenyl)amino)prop-1-yn-1-yl)-1-(2,2,2-trifluoroethyl)-1H-indol-4-yl)amino)piperidin-1-yl)acetyl)piperidin-4-yl)methyl)piperazin-1-yl)ethan-1-one (**TRAP-1** or **XJZ-06-462**)

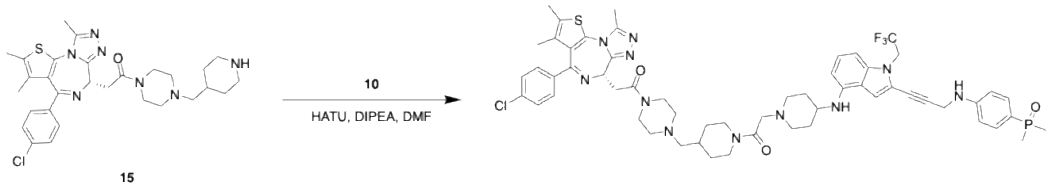


15

XJZ-06-462

The corresponding compound was prepared following **Procedure C** using intermediate **15** (5.1 mg, 8.9 µmol, 1 eq.) prepared through **Procedure A** (92% in two steps, LC/MS(ESI) for $C_{29}H_{37}ClN_7OS$ $[M+H]^+$: m/z calcd, 566.25; found, 566.30.). Purification by HPLC with MeOH/$H_2O$ (0.035% TFA) to afford the product as a white solid (3.99 mg, 40%). LC/MS(ESI) for $C_{57}H_{67}ClF_3N_{11}O_3PS$ $[M+H]^+$: m/z calcd, 1108.45; found, 1108.45. $^1$H NMR (500 MHz, $d^6$-DMSO) δ 9.53 (br, 1H), 7.50 (dt, *J* = 8.2, 5.4 Hz, 4H), 7.44 (d, *J* = 8.3 Hz, 2H), 7.15 – 7.10 (m, 1H), 7.07 – 7.00 (m, 1H), 6.82 – 6.79 (m, 2H), 6.76 (d, *J* = 8.2 Hz, 1H), 6.28 – 6.20 (m, 1H), 4.95 (q, *J* = 9.4 Hz, 2H), 4.58 (t, *J* = 6.7 Hz, 1H), 4.47 – 4.32 (m, 3H), 4.29 (m, 3H), 3.81 – 3.54 (m, 6H), 3.49 (br, 1H), 3.40 – 3.24 (m, 2H), 3.23 – 3.30 (m, 8H), 2.75 (t, *J* = 11.0 Hz, 2H), 2.61 (s, 3H), 2.42 (s, 3H), 2.24 – 2.09 (m, 4H), 1.98 (br, 1H), 1.91 – 1.75 (m, 4H), 1.63 (s, 3H), 1.56 (d, *J* = 13.0 Hz, 6H), 1.30 – 1.16 (m, 1H), 1.15 – 1.03 (m, 1H).

Synthesis of (S)-2-(4-(4-chlorophenyl)-2,3,9-trimethyl-6H-thieno[3,2-f][1,2,4]triazolo[4,3-a][1,4]diazepin-6-yl)-1-(4-((4-(2-(4-((2-(3-((4-(dimethylphosphoryl)phenyl)amino)prop-1-yn-1-yl)-1-(2,2,2-trifluoroethyl)-1H-indol-4-yl)amino)piperidin-1-yl)acetyl)piperazin-1-yl)methyl)piperidin-1-yl)ethan-1-one (**TRAP-2** or **XJZ-06-463**)

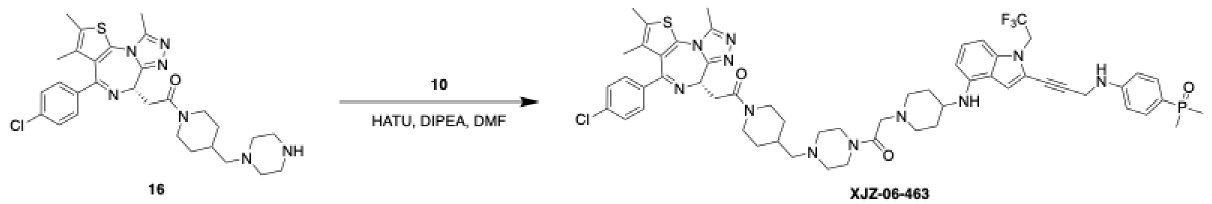


The corresponding compound was prepared following **Procedure C** using intermediate **16** (5.1 mg, 8.9 µmol, 1 eq.) prepared through **Procedure A** (92% in two steps, LC/MS(ESI) for $C_{29}H_{37}ClN_7OS$ $[M+H]^+$: m/z calcd, 566.25; found, 566.35). Purification by HPLC with MeOH/$H_2O$ (0.035% TFA) to afford the product as a white solid (3.76 mg, 38%). LC/MS(ESI) for $C_{57}H_{67}ClF_3N_{11}O_3PS$ $[M+H]^+$: m/z calcd, 1108.45; found, 1108.35. $^1$H NMR (500 MHz, $d^6$-DMSO) δ 9.74 (br, 1H), 7.50 (ddd, *J* = 8.6, 6.9, 4.0 Hz, 4H), 7.44 (dd, *J* = 8.7, 2.6 Hz, 2H), 7.17 – 7.11 (m, 1H), 7.08 – 7.00 (m, 1H), 6.81 (dd, *J* = 8.6, 2.2 Hz, 2H), 6.76 (d, *J* = 8.5 Hz, 1H), 6.25 (t, *J* = 9.2 Hz, 1H), 4.95 (q, *J* = 9.0 Hz, 1H), 4.59 (t, *J* = 6.7 Hz, 1H), 4.49 – 4.32 (m, 4H), 4.29 (s, 2H), 4.26 – 4.14 (m, 1H), 3.69 – 3.60 (m, 3H), 3.59 – 3.52 (m, 2H), 3.42 – 3.35 (m, 2H), 3.32 (br, 1H), 3.13 (m, 8H), 2.68 – 2.61 (m, 1H), 2.60 (s, 3H), 2.42 (s, 3H), 2.30 – 2.05 (m, 4H), 1.98 (br, 1H), 1.93 – 1.74 (m, 4H), 1.63 (s, 3H), 1.56 (d, *J* = 13.2 Hz, 6H), 1.41 – 1.17 (m, 1H), 1. 16 – 0.94 (m, 1H).

Synthesis of (S)-2-(4-(4-chlorophenyl)-2,3,9-trimethyl-6H-thieno[3,2-f][1,2,4]triazolo[4,3-a][1,4]diazepin-6-yl)-1-(9-(2-(4-((2-(3-((4-(dimethylphosphoryl)phenyl)amino)prop-1-yn-1-yl)-1-(2,2,2-trifluoroethyl)-1H-indol-4-yl)amino)piperidin-1-yl)acetyl)-3,9-diazaspiro[5.5]undecan-3-yl)ethan-1-one (**TRAP-3** or **XJZ-06-464**)

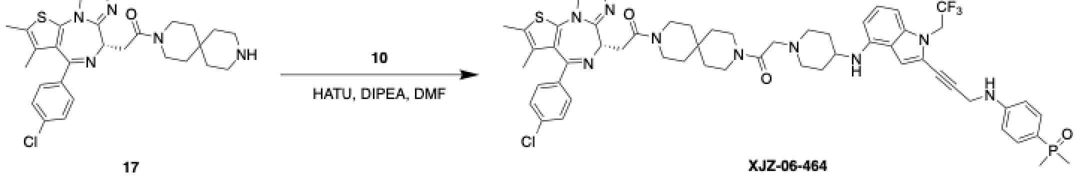

The corresponding compound was prepared following **Procedure C** using intermediate **17** (4.8 mg, 8.9 µmol, 1 eq.) prepared through **Procedure A** (90% in two steps, LC/MS: m/z 537.24). Purification by HPLC with MeOH/$H_2O$ (0.035% TFA) to afford the product as a white solid (3.24 mg, 34%). LC/MS(ESI) for $C_{56}H_{64}ClF_3N_{10}O_3PS$ $[M+H]^+$: m/z calcd, 1079.43; found, 1079.44. $^1H$ NMR (500 MHz, $d^6$-DMSO) δ 9.50 (br, 1H), 7.55 – 7.46 (m, 4H), 7.46 – 7.41 (m, 2H), 7.13 (d, *J* = 4.6 Hz, 1H), 7.08 – 6.98 (m, 1H), 6.84 – 6.78 (m, 2H), 6.76 (d, *J* = 8.4 Hz, 1H), 6.24 (dd, *J* = 13.2, 7.9 Hz, 1H), 4.94 (q, *J* = 9.1 Hz, 2H), 4.59 (t, *J* = 6.6 Hz, 1H), 4.34 – 4.27 (m, 4H), 3.69 – 3.45 (m, 10H), 3.43 – 3.28 (m, 4H), 3.21 – 3.06 (m, 2H), 2.60 (s, 3H), 2.42 (s, 3H), 2.25 – 2.08 (m, 2H), 1.92 – 1.72 (m, 2H), 1.63 (s, 3H), 1.61 – 1.52 (m, 10H), 1.50 – 1.47 (m, 2H), 1.45 – 1.41 (m, 2H).

Synthesis of (*S*)-2-(4-(4-chlorophenyl)-2,3,9-trimethyl-6*H*-thieno[3,2-*f*][1,2,4]triazolo[4,3-*a*][1,4]diazepin-6-yl)-1-(6-(2-(4-((2-(3-((4-(dimethylphosphoryl)phenyl)amino)prop-1-yn-1-yl)-1-(2,2,2-trifluoroethyl)-1*H*-indol-4-yl)amino)piperidin-1-yl)acetyl)-2,6-diazaspiro[3.3]heptan-2-yl)ethan-1-one (**XJZ-06-465**)

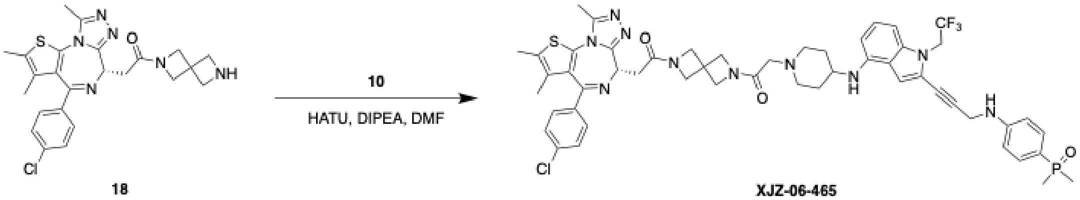


The corresponding compound was prepared following **Procedure C** using intermediate **18** (4.3 mg, 8.9 µmol, 1 eq.) prepared through **Procedure A** (83% in two steps, LC/MS(ESI) for $C_{24}H_{26}ClN_6O_3S$ $[M+H]^+$: m/z calcd, 481.16; found, 481.17). Purification by HPLC with MeOH/$H_2O$ (0.035% TFA) to afford the product as a white solid (3.32 mg, 36%). LC/MS(ESI) for $C_{52}H_{56}ClF_3N_{10}O_3PS$ $[M+H]^+$: m/z calcd, 1023.36; found, 1023.42. $^1H$ NMR (500 MHz, $d^6$-DMSO) δ 9.86 – 9.68 (m, 1H), 7.53 – 7.46 (m, 4H), 7.44 (d, *J* = 1.3 Hz, 2H), 7.12 (s, 1H), 7.03 (t, *J* = 7.9 Hz, 1H), 6.80 (dd, *J* = 8.7, 2.1 Hz, 2H), 6.75 (d, *J* = 8.3 Hz, 1H), 6.23 (t, *J* = 8.0 Hz, 1H), 4.94 (q, *J* = 9.3 Hz, 2H), 4.58 (d, *J* = 8.9 Hz, 1H), 4.54 – 4.44 (m, 2H), 4.42 – 4.31 (m, 2H), 4.29 (s, 2H), 4.23 – 4.16 (m, 2H), 4.14 – 4.01 (m, 4H), 3.98 (s, 2H), 3.62 – 3.56 (m, 1H), 3.54 (d, *J* = 11.0 Hz, 2H), 3.33 (s, 1H), 3.25 (dd, *J* = 15.5, 7.0 Hz, 1H), 3.19 – 3.09 (m, 3H), 2.60 (s, 3H), 2.41 (s, 3H), 2.17 (d, *J* = 13.9 Hz, 1H), 1.78 (d, *J* = 12.4 Hz, 1H), 1.62 (s, 3H), 1.56 (d, *J* = 13.3 Hz, 6H).

Synthesis of (*S*)-*N*-((1-(2-(4-(4-chlorophenyl)-2,3,9-trimethyl-6*H*-thieno[3,2-*f*][1,2,4]triazolo[4,3-*a*][1,4]diazepin-6-yl)acetyl)azetidin-3-yl)methyl)-2-(4-((2-(3-((4-(dimethylphosphoryl)phenyl)amino)prop-1-yn-1-yl)-1-(2,2,2-trifluoroethyl)-1*H*-indol-4-yl)amino)piperidin-1-yl)acetamide (**XJZ-06-471**)

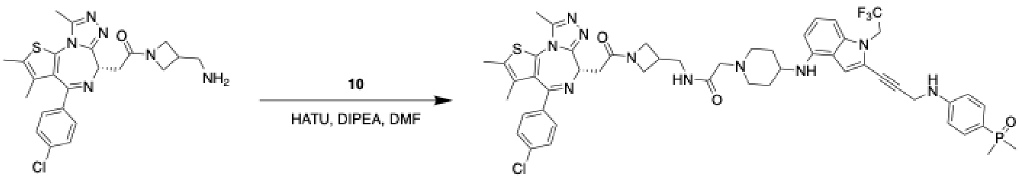

19 XJZ-06-471

The corresponding compound was prepared following **Procedure C** using intermediate **19** (4.2 mg, 8.9 µmol, 1 eq.) prepared through **Procedure A** (87% in two steps, LC/MS(ESI) for $C_{23}H_{26}ClN_6OS$ $[M+H]^+$: m/z calcd, 469.16; found, 469.17). Purification by prep-TLC with MeOH/DCM to afford the product as a white solid (2.38 mg, 26%). LC/MS(ESI) for $C_{51}H_{56}ClF_3N_{10}O_3PS$ $[M+H]^+$: m/z calcd, 1011.36; found, 1011.42. $^1$H NMR (500 MHz, d$^6$-DMSO) δ 8.05 – 7.97 (m, 1H), 7.52 – 7.45 (m, 4H), 7.42 (dd, *J* = 8.6, 3.3 Hz, 2H), 7.11 (s, 1H), 6.97 (dt, *J* = 11.1, 8.0 Hz, 1H), 6.80 (dd, *J* = 8.7, 2.2 Hz, 2H), 6.67 (t, *J* = 6.8 Hz, 2H), 6.13 (dd, *J* = 11.8, 7.9 Hz, 1H), 5.48 (dd, *J* = 7.5, 4.0 Hz, 1H), 4.90 (q, *J* = 7.0 Hz, 2H), 4.46 (td, *J* = 7.0, 3.0 Hz, 1H), 4.36 (dt, *J* = 29.3, 8.5 Hz, 1H), 4.28 (d, *J* = 6.2 Hz, 2H), 4.05 (dt, *J* = 31.1, 7.0 Hz, 1H), 3.93 – 3.81 (m, 1H), 3.68 – 3.58 (m, 1H), 3.51 (s, 2H), 3.44 – 3.37 (m, 1H), 3.20 (dt, *J* = 14.0, 6.7 Hz, 1H), 3.16 – 3.04 (m, 1H), 3.01 – 2.94 (m, 2H), 2.89 – 2.72 (m, 3H), 2.58 (d, *J* = 3.9 Hz, 3H), 2.41 – 2.37 (m, 3H), 2.29 – 2.15 (m, 2H), 2.01 – 1.85 (m, 2H), 1.61 (d, *J* = 6.4 Hz, 3H), 1.58 – 1.49 (m, 8H).

Synthesis of (*S*)-2-(4-(4-chlorophenyl)-2,3,9-trimethyl-6*H*-thieno[3,2-*f*][1,2,4]triazolo[4,3-*a*][1,4]diazepin-6-yl)-1-(4-(1-(2-(4-((2-(3-((4-(dimethylphosphoryl)phenyl)amino)prop-1-yn-1-yl)-1-(2,2,2-trifluoroethyl)-1*H*-indol-4-yl)amino)piperidin-1-yl)acetyl)piperidin-4-yl)piperazin-1-yl)ethan-1-one (**XJZ-06-537**)

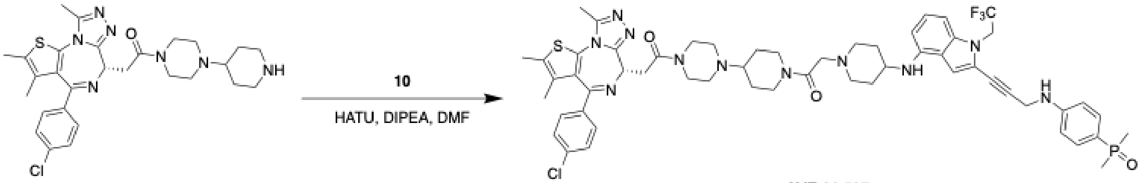

20 XJZ-06-537

The corresponding compound was prepared following **Procedure C** using intermediate **20** (4.9 mg, 8.9 µmol, 1 eq.) prepared through **Procedure A** (91% in two steps, LC/MS(ESI) for $C_{28}H_{35}ClN_7OS$ $[M+H]^+$: m/z calcd, 552.23; found, 552.29). Purification by HPLC with MeOH/$H_2O$ (0.035% TFA) to afford the product as a white solid (1.78 mg, 18%). LC/MS(ESI) for $C_{56}H_{65}ClF_3N_{11}O_3PS$ $[M+H]^+$: m/z calcd, 1094.44; found, 1094.44. $^1$H NMR (500 MHz, d$^6$-DMSO) δ 9.67 – 9.46 (m, 1H), 7.50 (dt, *J* = 8.6, 5.6 Hz, 4H), 7.44 (d, *J* = 8.3 Hz, 2H), 7.16 – 7.10 (m, 1H), 7.08 – 7.00 (m, 1H), 6.83 – 6.79 (m, 2H), 6.75 (d, *J* = 8.6 Hz, 1H), 6.28 – 6.21 (m, 1H), 4.95 (q, *J* = 9.3 Hz, 2H), 4.58 (t, *J* = 6.8 Hz, 1H), 4.56 – 4.34 (m, 5H), 4.34 – 4.23 (m, 3H), 3.91 – 3.76 (m, 1H), 3.72 (dd, *J* = 16.3, 7.4 Hz, 1H), 3.66 – 3.41 (m, 7H), 3.41 – 3.20 (m, 1H), 3.20 – 3.04 (m, 2H), 3.02 (br, 1H), 2.72 (t, *J* = 12.6 Hz, 1H), 2.61 (s, 3H), 2.42 (s, 3H), 2.28 – 2.05 (m, 5H), 2.05 – 1.88 (m, 1H), 1.89 – 1.75 (m, 1H), 1.75 – 1.61 (m, 5H), 1.59 – 1.49 (m, 8H).

Synthesis of (*S*)-2-(4-(4-chlorophenyl)-2,3,9-trimethyl-6*H*-thieno[3,2-*f*][1,2,4]triazolo[4,3-*a*][1,4]diazepin-6-yl)-1-(4-((1-(2-(4-((2-(3-((4-(dimethylphosphoryl)phenyl)amino)prop-1-yn-1-

yl)-1-(2,2,2-trifluoroethyl)-1*H*-indol-4-yl)amino)piperidin-1-yl)acetyl)piperidin-4-yl)methyl)piperidin-1-yl)ethan-1-one (**XJZ-06-540**)

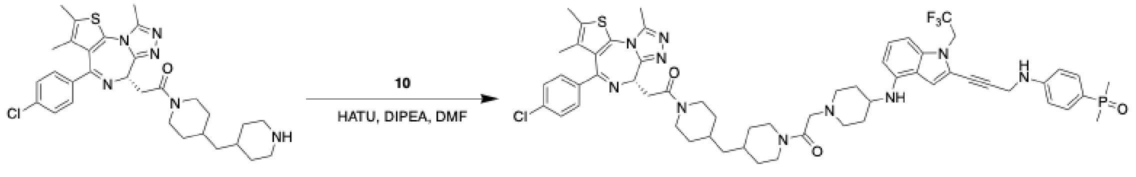


21

XJZ-06-540

The corresponding compound was prepared following **Procedure C** using intermediate **21** (5.0 mg, 8.9 µmol, 1 eq.) prepared through **Procedure A** (89% in two steps, LC/MS(ESI) for $C_{30}H_{38}ClN_6OS$ $[M+H]^+$: m/z calcd, 565.25; found, 565.35). Purification by HPLC with MeOH/$H_2O$ (0.035% TFA) to afford the product as a white solid (3.27 mg, 33%). LC/MS(ESI) for $C_{58}H_{68}ClF_3N_{10}O_3PS$ $[M+H]^+$: m/z calcd, 1107.46; found, 1107.50. $^1H$ NMR (500 MHz, $d^6$-DMSO) δ 9.47 (br, 1H), 7.53 – 7.46 (m, 4H), 7.44 (dd, *J* = 8.5, 5.0 Hz, 2H), 7.12 (s, 1H), 7.04 (td, *J* = 8.0, 3.4 Hz, 1H), 6.80 (dd, *J* = 8.7, 2.2 Hz, 2H), 6.75 (d, *J* = 8.3 Hz, 1H), 6.24 (dd, *J* = 12.2, 7.8 Hz, 1H), 4.94 (q, *J* = 9.1 Hz, 2H), 4.58 (t, *J* = 6.7 Hz, 1H), 4.37 (d, *J* = 13.5 Hz, 2H), 4.33 – 4.22 (m, 4H), 4.13 (d, *J* = 13.1 Hz, 2H), 3.66 – 3.53 (m, 4H), 3.42 – 3.33 (m, 2H), 3.19 – 3.00 (m, 4H), 2.69 (t, *J* = 12.4 Hz, 1H), 2.63 – 2.56 (m, 4H), 2.42 (s, 3H), 2.18 (d, *J* = 13.4 Hz, 2H), 2.13 – 2.05 (m, 1H), 2.02 – 1.92 (m, 1H), 1.83 – 1.65 (m, 6H), 1.63 (s, 3H), 1.56 (d, *J* = 13.2 Hz, 6H), 1.25 – 1.06 (m, 4H), 1.01 – 0.87 (m, 2H).

**Procedure D:** Preparation of 9-ethyl-7-(4-methylthiophen-2-yl)-9*H*-carbazole-3-carbaldehyde (Compound **25**)

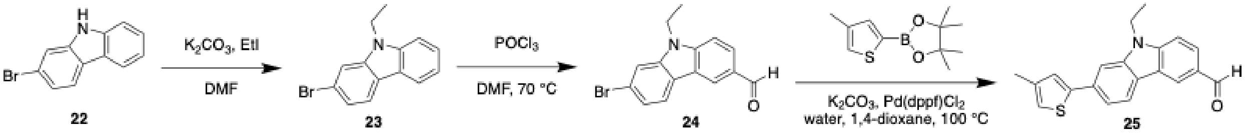


22 23 24 25

A solution containing 2-bromo-9H-carbazole **22** (2.00 g, 9.13 mmol, 1 eq.) and NaOH (1.40 g, 35 mmol, 4.3 eq.) in DMF (20 mL) was stirred for 30 minutes under room temperature before the addition of iodoethane (1.13 mL, 14.0 mmol, 1.72 eq.). The reaction was stirred overnight at room temperature. The resulting solution was then diluted in water and extracted with ethyl acetate 3 times. The combined organic phase was washed with 5% LiCl aqueous solution 3 times, dried with anhydrous sodium sulfate, and concentrated under reduced pressure. The crude was purified with flash chromatography with 0-10% EA/Hex to afford intermediate **23** as a white solid (2.1 g, 94%). LC/MS(ESI) for $C_{14}H_{13}BrN$ $[M+H]^+$: m/z calcd, 274.02; found, 275.19.

The intermediate **23** (2.4 g, 8.8 mmol, 1 eq.) was dissolved in DMF (16 mL), and the solution was cooled to 0 °C before the addition of phosphoryl chloride (2.4 mL, 26 mmol, 3 eq.) dropwise. The reaction was then warmed to room temperature and then heated at 70 °C overnight. After cooling to room temperature, the reaction was quenched with ice and neutralized with 5% aqueous sodium hydroxide. The resulting solution was extracted with ethyl acetate 3 times, and the combined organic layer was washed with water and brine, dried with anhydrous sodium sulfate, and concentrated under reduced pressure. The crude was then purified with flash chromatography with

0-10% EA/Hex to afford intermediate **24** as a white solid (1.90 g, 72%). LC/MS(ESI) for $C_{15}H_{13}BrNO$ [M+H]$^+$: m/z calcd, 302.02; found, 303.10.

A solution of intermediate **24** (800 mg, 2.65 mmol, 1 eq.), 4,4,5,5-tetramethyl-2-(4-methylthiophen-2-yl)-1,3,2-dioxaborolane (890 mg, 3.97 mmol, 1.5 eq.), potassium carbonate (732 mg, 5.30 mmol, 2 eq.), and [1,1′-bis(diphenylphosphino)ferrocene] dichloropalladium(II) (194 mg, 265 µmol, 0.1 eq.) in 1,4-dioxane (30 mL) and water (2.65 mL) was stirred at 100 ℃ overnight under nitrogen. The resulting crude was concentrated under reduced pressure and then purified by flash chromatography 0-10% EA/Hex to afford compound **25** as a white solid (731 mg, 86.4%). LC/MS(ESI) for $C_{20}H_{18}NOS$ [M+H]$^+$: m/z calcd, 320.11; found, 320.05.

Synthesis of (9-ethyl-7-(4-methylthiophen-2-yl)-9*H*-carbazol-3-yl)methanamine (**PK9328 free amine**)

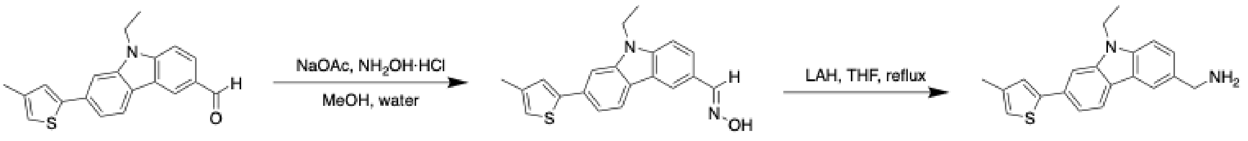


25 31 PK9328 free amine

The compound **25** (731 mg, 2.29 mmol, 1 eq.) was dissolved in MeOH (30 mL) followed by the addition of hydroxylamine hydrochloride (318 mg, 4.58 mmol, 2 eq.), sodium acetate (375 mg, 4.58 mmol, 2 eq.), and water (3.3 mL). The solution was stirred at room temperature for 2 hours and was then diluted with ethyl acetate. The resulting solution was washed with water, and the aqueous phase was extracted twice with ethyl acetate. The combined organic phase was then dried over anhydrous sodium sulfate and concentrated under reduced pressure to afford intermediate **31** (557 mg, 72.8%) which was used directly for the next step. LC/MS(ESI) for $C_{20}H_{19}N_2OS$ [M+H]$^+$: m/z calcd, 335.12; found, 355.26.

To a solution of compound **15** (557 mg, 1.73 mmol, 1 eq.) in dry THF (6 mL) was added a suspension of $LiAlH_4$ (228 mg, 6.00 mmol, 3.5 eq.) in dry THF (6 mL) dropwise. The resulting solution was refluxed for 1 hour before cooling to room temperature and quenched with MeOH. The solution was filtered and evaporated under reduced pressure, and the residue was then purified by flash chromatography with 0-20% MeOH/DCM to afford compound **PK9328 free amine** as a brown solid (250 mg, 45%). LC/MS(ESI) for $C_{20}H_{21}N_2S$ [M+H]$^+$: m/z calcd, 321.14; found, 321.26. $^1$H NMR (500 MHz, d$^6$-DMSO) δ 8.24 – 8.19 (m, 3H), 8.10 (dd, *J* = 8.1, 0.6 Hz, 1H), 7.89 (d, *J* = 1.5 Hz, 1H), 7.68 (d, *J* = 8.4 Hz, 1H), 7.56 (dd, *J* = 8.4, 1.8 Hz, 1H), 7.51 (d, *J* = 1.4 Hz, 1H), 7.48 (dd, *J* = 8.1, 1.5 Hz, 1H), 7.14 (t, *J* = 1.3 Hz, 1H), 4.52 (q, *J* = 7.1 Hz, 2H), 4.18 (s, 2H), 2.28 (d, *J* = 1.1 Hz, 3H), 1.33 (t, *J* = 7.1 Hz, 3H).

**Procedure E:** Synthesis of (*S*)-2-(4-(4-chlorophenyl)-2,3,9-trimethyl-6*H*-thieno[3,2-*f*][1,2,4]triazolo[4,3-*a*][1,4]diazepin-6-yl)-*N*-(6-(((9-ethyl-7-(5-methylthiophen-2-yl)-9*H*-carbazol-3-yl)methyl)amino)hexyl)acetamide (**XJZ-06-408**)

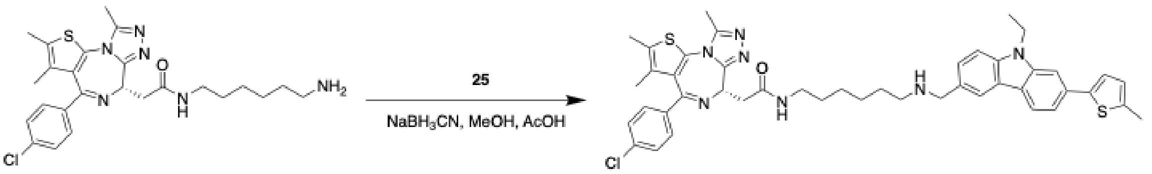


11

XJZ-06-408

To a solution of intermediate **11** (7.8 mg, 16 µmol, 1 eq.) and compound **25** (5.0 mg, 16 µmol, 1 eq.) in MeOH (300 µL) was acetic acid to tune the pH to 5 before the addition of sodium cyanoborohydride (3.0 mg, 47 µmol, 3 eq.). The reaction was stirred overnight at room temperature and was then purified by HPLC with MeOH/$H_2O$ (0.035% TFA) to give compound **XJZ-06-408** as a white solid (6.76 mg, 54%). LC/MS(ESI) for $C_{45}H_{49}ClN_7OS_2$ $[M+H]^+$: m/z calcd, 802.31; found, 802.30. $^1H$ NMR (500 MHz, $d^6$-DMSO) δ 8.75 (s, 2H), 8.24 (d, *J* = 1.7 Hz, 1H), 8.19 (t, *J* = 5.7 Hz, 1H), 8.11 (d, *J* = 8.1 Hz, 1H), 7.91 (d, *J* = 1.5 Hz, 1H), 7.70 (d, *J* = 8.4 Hz, 1H), 7.57 (dd, *J* = 8.4, 1.7 Hz, 1H), 7.52 (d, *J* = 1.4 Hz, 1H), 7.51 – 7.47 (m, 3H), 7.42 (d, *J* = 8.7 Hz, 2H), 7.15 (t, *J* = 1.3 Hz, 1H), 4.57 – 4.47 (m, 3H), 4.30 (t, *J* = 5.8 Hz, 2H), 3.24 (d, *J* = 7.2 Hz, 2H), 3.12 (hept, *J* = 6.4 Hz, 2H), 3.04 – 2.93 (m, 2H), 2.59 (s, 3H), 2.40 (s, 3H), 2.29 (d, *J* = 1.1 Hz, 3H), 1.68 – 1.62 (m, 2H), 1.61 (s, 3H), 1.46 (t, *J* = 6.9 Hz, 2H), 1.37 – 1.31 (m, 7H).

Synthesis of (*S*)-2-(4-(4-chlorophenyl)-2,3,9-trimethyl-6*H*-thieno[3,2-*f*][1,2,4]triazolo[4,3-*a*][1,4]diazepin-6-yl)-*N*-(8-(((9-ethyl-7-(5-methylthiophen-2-yl)-9*H*-carbazol-3-yl)methyl)amino)octyl)acetamide (**XJZ-06-409**)

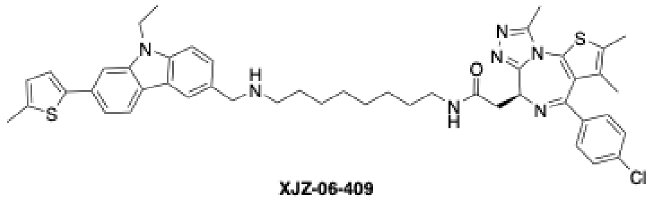

XJZ-06-409

The corresponding compound was prepared following **Procedure E** using compound **25** (5.0 mg, 16 µmol, 1 eq.) and intermediate **12**. Purification by HPLC with MeOH/$H_2O$ (0.035% TFA) to afford the product as a white solid (6.56 mg, 50%). LC/MS(ESI) for $C_{47}H_{53}ClN_7OS_2$ $[M+H]^+$: m/z calcd, 830.34; found, 830.30. $^1H$ NMR (500 MHz, $d^6$-DMSO) δ 8.72 (br, 2H), 8.23 (d, *J* = 1.6 Hz, 1H), 8.16 (t, *J* = 5.7 Hz, 1H), 8.11 (d, *J* = 8.1 Hz, 1H), 7.90 (d, *J* = 1.5 Hz, 1H), 7.69 (d, *J* = 8.4 Hz, 1H), 7.56 (dd, *J* = 8.4, 1.7 Hz, 1H), 7.52 (d, *J* = 1.4 Hz, 1H), 7.48 (dq, *J* = 9.2, 1.8 Hz, 3H), 7.44 – 7.39 (m, 2H), 7.15 (p, *J* = 1.1 Hz, 1H), 4.56 – 4.45 (m, 3H), 4.29 (t, *J* = 5.8 Hz, 2H), 3.28 – 3.15 (m, 2H), 3.09 (qd, *J* = 6.8, 2.9 Hz, 2H), 2.99 – 2.91 (m, 2H), 2.58 (s, 3H), 2.40 (d, *J* = 1.0 Hz, 3H), 2.28 (d, *J* = 1.1 Hz, 3H), 1.66 – 1.58 (m, 5H), 1.47 – 1.39 (m, 2H), 1.36 – 1.25 (m, 11H).

Synthesis of (*S*)-2-(4-(4-chlorophenyl)-2,3,9-trimethyl-6*H*-thieno[3,2-*f*][1,2,4]triazolo[4,3-*a*][1,4]diazepin-6-yl)-*N*-(2-(2-(2-(((9-ethyl-7-(5-methylthiophen-2-yl)-9*H*-carbazol-3-yl)methyl)amino)ethoxy)ethoxy)ethyl)acetamide (**XJZ-06-411**)

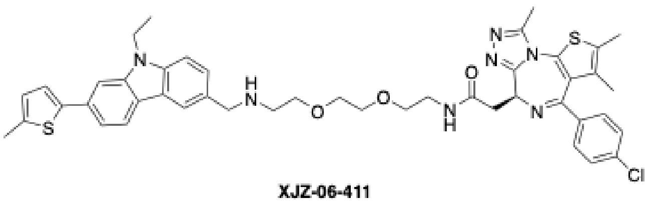

XJZ-06-411

The corresponding compound was prepared following **Procedure E** using compound **25** (5.0 mg, 16 µmol, 1 eq.) and intermediate **13**. Purification by HPLC with MeOH/$H_2O$ (0.035% TFA) to afford the product as a white solid (7.86 mg, 60%). LC/MS(ESI) for $C_{45}H_{49}ClN_7O_3S_2$ $[M+H]^+$: m/z calcd, 834.30; found, 834.25. $^1H$ NMR (500 MHz, $d^6$-DMSO) δ 8.88 (br, 2H), 8.26 (t, *J* = 5.7 Hz, 1H), 8.23 (d, *J* = 1.7 Hz, 1H), 8.12 – 8.06 (m, 1H), 7.89 (d, *J* = 1.6 Hz, 1H), 7.67 (d, *J* = 8.5 Hz, 1H), 7.57 (dd, *J* = 8.4, 1.7 Hz, 1H), 7.51 (d, *J* = 1.5 Hz, 1H), 7.48 – 7.43 (m, 3H), 7.42 – 7.37 (m, 2H), 7.14 (t, *J* = 1.3 Hz, 1H), 4.50 (q, *J* = 7.0 Hz, 3H), 4.33 (t, *J* = 5.6 Hz, 2H), 3.71 (t, *J* = 5.2 Hz, 2H), 3.63 – 3.59 (m, 4H), 3.48 – 3.47 (m, 2H), 3.32 – 3.20 (m, 4H), 3.15 (t, *J* = 5.7 Hz, 2H), 2.57 (s, 3H), 2.38 (s, 3H), 2.28 (d, *J* = 1.1 Hz, 3H), 1.59 (s, 3H), 1.32 (t, *J* = 7.1 Hz, 3H).

Synthesis of (*S*)-2-(4-(4-chlorophenyl)-2,3,9-trimethyl-6*H*-thieno[3,2-*f*][1,2,4]triazolo[4,3-*a*][1,4]diazepin-6-yl)-*N*-(1-(9-ethyl-7-(5-methylthiophen-2-yl)-9*H*-carbazol-3-yl)-5,8,11,14-tetraoxa-2-azahexadecan-16-yl)acetamide (**XJZ-06-392**)

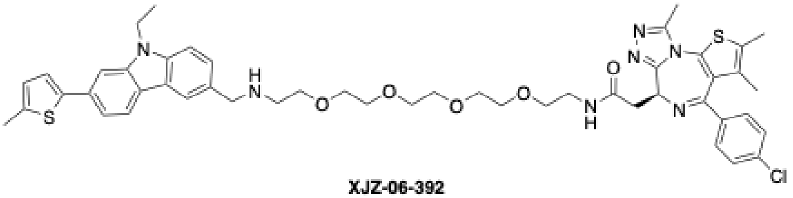

XJZ-06-392

The corresponding compound was prepared following **Procedure E** using compound **25** (5.0 mg, 16 µmol, 1 eq.) and intermediate **14**. Purification by prep-TLC with MeOH/DCM to afford the product as a white solid (5.80 mg, 40%). LC/MS(ESI) for $C_{49}H_{57}ClN_7O_5S_2$ $[M+H]^+$: m/z calcd, 922.35; found, 922.36. $^1H$ NMR (500 MHz, $d^6$-DMSO) δ 8.27 (t, *J* = 5.7 Hz, 1H), 8.12 – 8.06 (m, 2H), 7.83 (s, 1H), 7.54 (d, *J* = 8.3 Hz, 1H), 7.51 – 7.38 (m, 7H), 7.12 (s, 1H), 4.53 – 4.43 (m, 3H), 3.92 (s, 2H), 3.50 (d, *J* = 6.6 Hz, 14H), 3.43 (t, *J* = 5.9 Hz, 2H), 3.27 – 3.16 (m, 4H), 2.75 (q, *J* = 6.3 Hz, 2H), 2.58 (s, 3H), 2.38 (s, 3H), 2.27 (s, 3H), 1.60 (s, 3H), 1.32 (t, *J* = 7.1 Hz, 3H).

Synthesis of (*S*)-2-(4-(4-chlorophenyl)-2,3,9-trimethyl-6*H*-thieno[3,2-*f*][1,2,4]triazolo[4,3-*a*][1,4]diazepin-6-yl)-1-(4-((1-((9-ethyl-7-(5-methylthiophen-2-yl)-9*H*-carbazol-3-yl)methyl)piperidin-4-yl)methyl)piperazin-1-yl)ethan-1-one (**XJZ-06-429**)

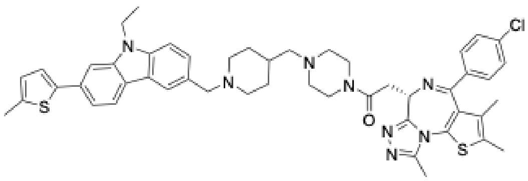

XJZ-06-429

The corresponding compound was prepared following **Procedure E** using compound **25** (5.0 mg, 16 µmol, 1 eq.) and intermediate **15**. Purification by prep-TLC with MeOH/DCM to afford the product as a white solid (3.47 mg, 25%). LC/MS(ESI) for $C_{49}H_{54}ClN_{8}OS_{2}$ $[M+H]^{+}$: m/z calcd, 869.35; found 869.31. $^{1}$H NMR (500 MHz, d$^{6}$-DMSO) δ 8.18 – 8.04 (m, 2H), 7.85 (s, 1H), 7.61 – 7.56 (m, 1H), 7.50 (s, 1H), 7.44 (d, *J* = 8.1 Hz, 3H), 7.18 (d, *J* = 8.2 Hz, 2H), 7.13 (s, 1H), 7.05 (d, *J* = 8.1 Hz, 2H), 6.55 (s, 1H), 5.28 (s, 1H), 4.49 (q, *J* = 7.4 Hz, 2H), 4.20 (t, *J* = 7.6 Hz, 1H), 3.58 – 3.50 (m, 5H), 3.49 – 3.41 (m, 4H), 3.21 (dd, *J* = 16.4, 6.1 Hz, 2H), 2.85 (dd, *J* = 16.5, 7.0 Hz, 2H), 2.43 – 2.34 (m, 7H), 2.30 – 2.23 (m, 10H), 2.00 (s, 3H), 1.62 (s, 1H), 1.35 (t, *J* = 7.0 Hz, 3H).

Synthesis of (*S*)-2-(4-(4-chlorophenyl)-2,3,9-trimethyl-6*H*-thieno[3,2-*f*][1,2,4]triazolo[4,3-*a*][1,4]diazepin-6-yl)-1-(4-((4-((9-ethyl-7-(5-methylthiophen-2-yl)-9*H*-carbazol-3-yl)methyl)piperazin-1-yl)methyl)piperidin-1-yl)ethan-1-one (**XJZ-06-430**)

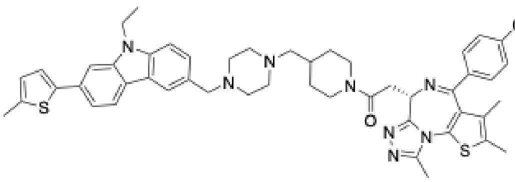

XJZ-06-430

The corresponding compound was prepared following **Procedure E** using compound **25** (5.0 mg, 16 µmol, 1 eq.) and intermediate **16**. Purification by prep-TLC with MeOH/DCM to afford the product as a white solid (4.40 mg, 32%). LC/MS(ESI) for $C_{49}H_{54}ClN_{8}OS_{2}$ $[M+H]^{+}$: m/z calcd, 869.35; found 869.21 $[M+1]^{+}$. $^{1}$H NMR (500 MHz, d$^{6}$-DMSO) δ 8.13 (d, *J* = 8.1 Hz, 1H), 8.00 (d, *J* = 1.6 Hz, 1H), 7.83 (d, *J* = 1.5 Hz, 1H), 7.53 (d, *J* = 8.4 Hz, 1H), 7.49 (d, *J* = 1.5 Hz, 1H), 7.43 (s, 1H), 7.18 (d, *J* = 8.2 Hz, 2H), 7.12 (s, 1H), 7.10 – 7.02 (m, 2H), 5.28 (d, *J* = 4.4 Hz, 1H), 4.47 (q, *J* = 7.1 Hz, 2H), 4.35 – 4.31 (m, 1H), 4.19 (q, *J* = 5.9 Hz, 1H), 3.95 (d, *J* = 13.3 Hz, 1H), 3.60 (s, 2H), 3.57 – 3.47 (m, 1H), 3.19 (dt, *J* = 15.5, 7.3 Hz, 1H), 3.03 (t, *J* = 12.8 Hz, 1H), 2.85 (dt, *J* = 15.3, 7.3 Hz, 1H), 2.65 – 2.53 (m, 1H), 2.47 – 2.31 (m, 9H), 2.30 – 2.23 (m, 6H), 2.11 (d, *J* = 7.0 Hz, 2H), 2.01 (d, *J* = 5.7 Hz, 3H), 1.79 – 1.71 (m, 2H), 1.70 – 1.62 (m, 1H), 1.33 (t, *J* = 7.1 Hz, 3H), 1.16 – 1.00 (m, 1H), 0.90 – 0.83 (m, 1H).

Synthesis of (*S*)-2-(4-(4-chlorophenyl)-2,3,9-trimethyl-6*H*-thieno[3,2-*f*][1,2,4]triazolo[4,3-*a*][1,4]diazepin-6-yl)-1-(9-((9-ethyl-7-(5-methylthiophen-2-yl)-9*H*-carbazol-3-yl)methyl)-3,9-diazaspiro[5.5]undecan-3-yl)ethan-1-one (**XJZ-06-431**)

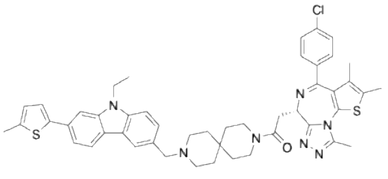

XJZ-06-431

The corresponding compound was prepared following **Procedure E** using compound **25** (5.0 mg, 16 µmol, 1 eq.) and intermediate **17**. Purification by prep-TLC with MeOH/DCM to afford the product as a white solid (1.74 mg, 13%). LC/MS(ESI) for $C_{48}H_{51}ClN_{7}OS_{2}$ $[M+H]^{+}$: m/z calcd, 840.33; found 840.20. $^{1}$H NMR (500 MHz, d$^{6}$-DMSO) δ 8.14 (d, *J* = 8.1 Hz, 1H), 8.01 (s, 1H), 7.83 (s, 1H), 7.53 (d, *J* = 8.4 Hz, 1H), 7.51 – 7.45 (m, 3H), 7.47 – 7.36 (m, 4H), 7.12 (s, 1H), 4.56

(t, *J* = 6.7 Hz, 1H), 4.47 (q, *J* = 7.3 Hz, 2H), 3.62 (s, 2H), 3.61 – 3.55 (m, 3H), 3.47 – 3.41 (m, 2H), 2.59 (s, 3H), 2.43 – 2.36 (m, 8H), 2.28 (s, 3H), 1.62 (s, 3H), 1.51 (br, 6H), 1.37 – 1.30 (m, 5H).

Synthesis of (*S*)-2-(4-(4-chlorophenyl)-2,3,9-trimethyl-6*H*-thieno[3,2-*f*][1,2,4]triazolo[4,3-*a*][1,4]diazepin-6-yl)-1-(6-((9-ethyl-7-(5-methylthiophen-2-yl)-9*H*-carbazol-3-yl)methyl)-2,6-diazaspiro[3.3]heptan-2-yl)ethan-1-one (**XJZ-06-432**)

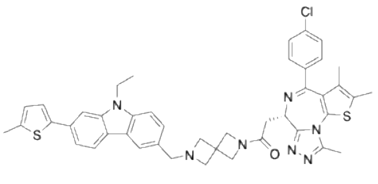

XJZ-06-432

The corresponding compound was prepared following **Procedure E** using compound **25** (5.0 mg, 16 µmol, 1 eq.) and intermediate **18**. Purification by prep-TLC with MeOH/DCM to afford the product as a white solid (3.41 mg, 28%). LC/MS(ESI) for $C_{44}H_{43}ClN_7OS_2$ $[M+H]^+$: m/z calcd, 784.27; found, 784.25 $[M+1]^+$. $^1$H NMR (500 MHz, d$^6$-DMSO) δ 10.27 – 10.11 (m, 1H), 8.22 (d, *J* = 8.2 Hz, 1H), 8.12 (dd, *J* = 8.1, 4.1 Hz, 1H), 7.91 (d, *J* = 1.7 Hz, 1H), 7.71 (dd, *J* = 8.5, 4.0 Hz, 1H), 7.57 – 7.43 (m, 4H), 7.44 – 7.37 (m, 1H), 7.22 (dd, *J* = 12.3, 8.4 Hz, 1H), 7.15 (t, *J* = 1.3 Hz, 1H), 7.06 (t, *J* = 7.9 Hz, 1H), 4.56 – 4.36 (m, 7H), 4.23 (d, *J* = 6.9 Hz, 2H), 4.15 – 3.98 (m, 2H), 3.22 (dd, *J* = 15.5, 6.8 Hz, 1H), 3.13 (dd, *J* = 7.4, 3.0 Hz, 1H), 2.59 (d, *J* = 5.0 Hz, 2H), 2.39 (d, *J* = 15.6 Hz, 3H), 2.35 (d, *J* = 5.5 Hz, 1H), 2.28 (s, 3H), 1.95 (s, 1H), 1.61 (s, 2H), 1.34 (td, *J* = 7.1, 2.4 Hz, 3H).

Synthesis of (R)-2-(4-(4-chlorophenyl)-2,3,9-trimethyl-6H-thieno[3,2-f][1,2,4]triazolo[4,3-a][1,4]diazepin-6-yl)-1-(4-((1-(2-(4-((2-(3-((4-(dimethylphosphoryl)phenyl)amino)prop-1-yn-1-yl)-1-(2,2,2-trifluoroethyl)-1H-indol-4-yl)amino)piperidin-1-yl)acetyl)piperidin-4-yl)methyl)piperazin-1-yl)ethan-1-one (**TRAP-1-Neg1** or **XJZ-06-483**)

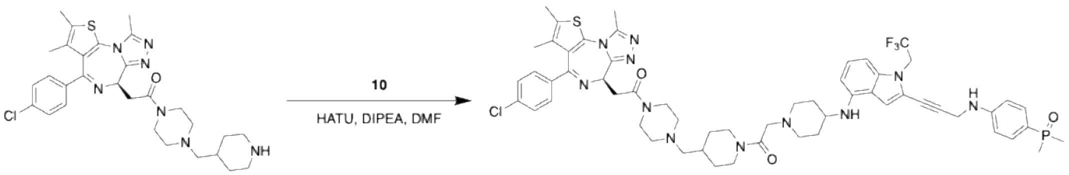


26 XJZ-06-483

The corresponding compound was prepared following **Procedure C** using intermediate **26** (prepared following **Procedure A** with (*R*)-2-(4-(4-chlorophenyl)-2,3,9-trimethyl-6*H*-thieno[3,2-*f*][1,2,4]triazolo[4,3-*a*][1,4]diazepin-6-yl)acetic acid, 89% in two steps) and compound **10** (10 mg, 17.8 µmol, 1 eq.). Purification by HPLC with MeOH/$H_2O$ (0.035% TFA) to afford the product as a white solid (4.41 mg, 22%). LC/MS(ESI) for $C_{57}H_{67}ClF_3N_{11}O_3PS$ $[M+H]^+$: m/z calcd, 1108.45; found, 1108.35. $^1$H NMR (500 MHz, d$^6$-DMSO) δ 9.53 (s, 1H), 7.54 – 7.46 (m, 4H), 7.44 (d, *J* = 8.6 Hz, 2H), 7.12 (s, 1H), 7.04 (s, 1H), 6.81 (dd, *J* = 8.6, 2.2 Hz, 2H), 6.76 (d, *J* = 8.2 Hz, 1H), 6.25 (d, *J* = 7.9 Hz, 1H), 4.95 (q, *J* = 9.5 Hz, 2H), 4.58 (t, *J* = 6.7 Hz, 1H), 4.46 – 4.32 (m, 3H), 4.32 – 4.22 (m, 3H), 3.81 – 3.53 (m, 8H), 3.52 – 3.41 (m, 1H), 3.40 – 3.25 (m, 1H), 3.23 – 3.03 (m, 7H), 2.95 (br, 1H), 2.74 (t, *J* = 11.5 Hz, 1H), 2.61 (s, 3H), 2.42 (s, 3H), 2.25 – 2.09 (m, 3H),

2.04 – 1.74 (m, 4H), 1.63 (s, 3H), 1.56 (dd, *J* = 13.2, 2.0 Hz, 7H), 1.27 – 1.16 (m, 1H), 1.15 – 1.03 (m, 1H).

Synthesis of (*R*)-2-(4-(4-chlorophenyl)-2,3,9-trimethyl-6*H*-thieno[3,2-*f*][1,2,4]triazolo[4,3-*a*][1,4]diazepin-6-yl)-1-(4-((4-(2-(4-((2-(3-((4-(dimethylphosphoryl)phenyl)amino)prop-1-yn-1-yl)-1-(2,2,2-trifluoroethyl)-1*H*-indol-4-yl)amino)piperidin-1-yl)acetyl)piperazin-1-yl)methyl)piperidin-1-yl)ethan-1-one (**TRAP-2-Neg1** or **XJZ-06-484**)

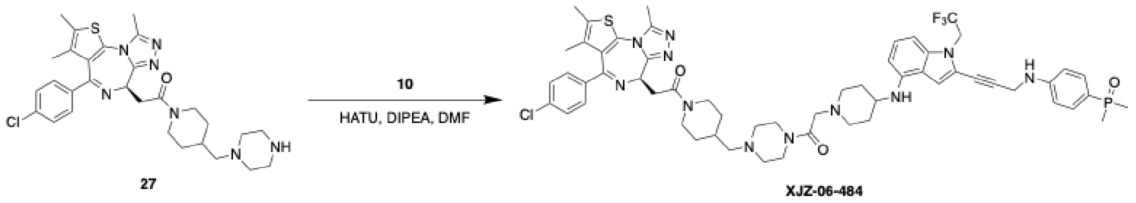

The corresponding compound was prepared following **Procedure C** using intermediate **27** (prepared following **Procedure A** with (*R*)-2-(4-(4-chlorophenyl)-2,3,9-trimethyl-6*H*-thieno[3,2-*f*][1,2,4]triazolo[4,3-*a*][1,4]diazepin-6-yl)acetic acid, 89% in two steps) and compound **10** (10 mg, 17.8 µmol, 1 eq.). Purification by HPLC with MeOH/$H_2O$ (0.035% TFA) to afford the product as a white solid (1.79 mg, 9%). LC/MS(ESI) for $C_{57}H_{67}ClF_3N_{11}O_3PS$ $[M+H]^+$: m/z calcd, 1108.45; found, 1108.35. $^1H$ NMR (500 MHz, $d^6$-DMSO) δ 9.72 (br, 1H), 7.50 (ddd, *J* = 8.6, 6.9, 4.1 Hz, 4H), 7.44 (dd, *J* = 8.6, 2.7 Hz, 2H), 7.13 (d, *J* = 2.8 Hz, 1H), 7.04 (t, *J* = 7.9 Hz, 1H), 6.84 – 6.73 (m, 3H), 6.25 (t, *J* = 9.2 Hz, 1H), 4.95 (q, *J* = 9.1 Hz, 2H), 4.59 (dd, *J* = 7.4, 6.1 Hz, 1H), 4.48 – 4.32 (m, 4H), 4.29 (s, 2H), 4.24 – 4.15 (m, 1H), 3.71 – 3.59 (m, 3H), 3.55 (d, *J* = 11.2 Hz, 2H), 3.39 (dt, *J* = 16.2, 5.5 Hz, 2H), 3.31 (br, 1H), 3.24 – 2.91 (m, 8H), 2.70 – 2.62 (m, 1H), 2.60 (s, 3H), 2.42 (s, 3H), 2.25 – 2.17 (m, 2H), 2.17 – 2.07 (m, 2H), 2.00 (br, 1H), 1.93 – 1.74 (m, 4H), 1.63 (s, 3H), 1.56 (d, *J* = 13.1 Hz, 6H), 1.36 – 1.23 (m, 1H), 1.12 – 1.01 (m, 1H).

Synthesis of (*R*)-2-(4-(4-chlorophenyl)-2,3,9-trimethyl-6*H*-thieno[3,2-*f*][1,2,4]triazolo[4,3-*a*][1,4]diazepin-6-yl)-1-(9-(2-(4-((2-(3-((4-(dimethylphosphoryl)phenyl)amino)prop-1-yn-1-yl)-1-(2,2,2-trifluoroethyl)-1*H*-indol-4-yl)amino)piperidin-1-yl)acetyl)-3,9-diazaspiro[5.5]undecan-3-yl)ethan-1-one (**TRAP-3-Neg1** or **XJZ-06-485**)

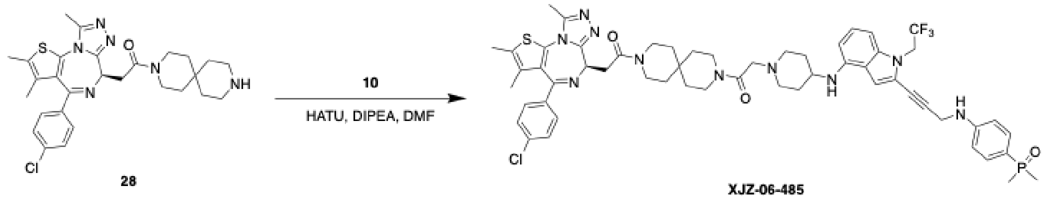

The corresponding compound was prepared following **Procedure C** using intermediate **28** (prepared following **Procedure A** with (*R*)-2-(4-(4-chlorophenyl)-2,3,9-trimethyl-6*H*-thieno[3,2-*f*][1,2,4]triazolo[4,3-*a*][1,4]diazepin-6-yl)acetic acid, 93% in two steps) and compound **10** (10 mg, 17.8 µmol, 1 eq.). Purification by HPLC with MeOH/$H_2O$ (0.035% TFA) to afford the product as a white solid (3.95 mg, 21%). LC/MS(ESI) for $C_{56}H_{64}ClF_3N_{10}O_3PS$ $[M+H]^+$: m/z calcd, 1079.43; found, 1079.34. $^1H$ NMR (500 MHz, $d^6$-DMSO) δ 9.49 (br, 1H), 7.50 (dt, *J* = 8.5, 5.6 Hz, 4H), 7.44 (d, *J* = 8.6 Hz, 2H), 7.12 (s, 1H), 7.04 (td, *J* = 8.0, 3.3 Hz, 1H), 6.84 – 6.71 (m, 3H), 6.24 (dd, *J* = 12.9, 7.9 Hz, 1H), 4.95 (q, *J* = 9.1 Hz, 2H), 4.59 (t, *J* = 6.7 Hz, 1H), 4.34 – 4.27 (m, 4H), 3.68

– 3.45 (m, 10H), 3.43 – 3.29 (m, 4H), 3.23 – 3.05 (m, 2H), 2.60 (s, 3H), 2.42 (s, 3H), 2.23 – 2.08 (m, 2H), 2.03 – 1.74 (m, 2H), 1.63 (s, 3H), 1.61 – 1.52 (m, 10H), 1.51 – 1.47 (m, 2H), 1.45 – 1.41 (m, 2H).

**Procedure F:** Preparation of 2-(4-((2-(3-((4-(dimethylphosphoryl)phenyl)(methyl)amino)prop-1-yn-1-yl)-1-(2,2,2-trifluoroethyl)-1*H*-indol-4-yl)(methyl)amino)piperidin-1-yl)acetic acid (Compound **30**)

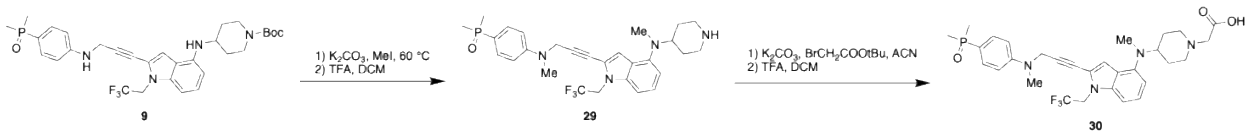


To a solution of compound **9** (100 mg, 166 µmol, 1 eq.) and potassium carbonate (45.9 mg, 332 µmol, 2 eq.) in DMF (2 mL) was added iodomethane (20.8 µL, 332 µmol, 2 eq.). The resulting mixture was stirred at 60 ℃ for overnight. The crude mixture was then purified by C18 chromatography column with 10-100% ACN/$H_2O$ to give the resulting product was dissolved in DCM (5 mL) and subjected to TFA (1 mL). After 2 h, the solution was concentrated under reduced pressure to afford intermediate **29** (55 mg, 62% in two steps) which was used directly as a crude. LC/MS(ESI) for $C_{28}H_{35}F_3N_4OP$ $[M+H]^+$: m/z calcd, 531.25; found, 531.32 $[M+1]^+$.

The intermediate **29** (55 mg, 100 µmol, 1 eq.) was dissolved in ACN (3 mL) followed by the addition of potassium carbonate (43 mg, 300 µmol, 3 eq.) and tert-butyl 2-bromoacetate (44 µL, 300 µmol, 3 eq.). After stirring for 2 hours at room temperature, the reaction mixture was purified by C18 chromatography column with 10-100% ACN/$H_2O$ to afford the reaction product. The reaction product was then dissolved in DCM (5 mL) with TFA (1 mL), and the solution was stirred for 2 hours before concentrating under reduced pressure to afford compound **30** in crude (15 mg, 25% in three steps). LC/MS(ESI) for $C_{30}H_{37}F_3N_4O_3P$ $[M+H]^+$: m/z calcd, 589.25; found, 589.39.

Synthesis of (*S*)-2-(4-(4-chlorophenyl)-2,3,9-trimethyl-6*H*-thieno[3,2-*f*][1,2,4]triazolo[4,3-*a*][1,4]diazepin-6-yl)-1-(4-((1-(2-(4-((2-(3-((4-(dimethylphosphoryl)phenyl)(methyl)amino)prop-1-yn-1-yl)-1-(2,2,2-trifluoroethyl)-1*H*-indol-4-yl)(methyl)amino)piperidin-1-yl)acetyl)piperidin-4-yl)methyl)piperazin-1-yl)ethan-1-one (**TRAP-1-Neg2** or **XJZ-06-488**)

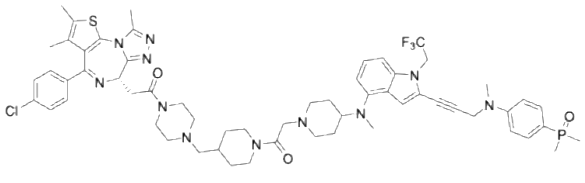

**XJZ-06-488**

The corresponding compound was prepared following **Procedure C** using intermediate **15** and compound **30** (5 mg, 8.5 µmol, 1 eq.). Purification by HPLC with MeOH/$H_2O$ (0.035% TFA) to afford the product as a white solid (3.53 mg, 37%). LC/MS(ESI) for $C_{59}H_{71}ClF_3N_{11}O_3PS$ $[M+H]^+$: m/z calcd, 1136.48; found, 1136.37. $^1H$ NMR (500 MHz, $d^6$-DMSO) δ 9.40 (s, 1H), 7.63 – 7.54 (m, 2H), 7.50 (d, *J* = 8.4 Hz, 2H), 7.44 (d, *J* = 8.6 Hz, 2H), 7.14 (td, *J* = 9.5, 7.4 Hz, 2H), 7.00 (dd, *J* = 8.9, 2.0 Hz, 2H), 6.83 (s, 1H), 6.62 (d, *J* = 7.0 Hz, 1H), 4.97 (q, *J* = 9.0 Hz, 2H), 4.58 (d, *J* = 5.1 Hz, 3H), 4.48 – 4.31 (m, 3H), 4.27 (d, *J* = 16.3 Hz, 1H), 4.17 (d, *J* = 16.0 Hz, 1H), 3.83 – 3.75

(s, 1H), 3.72 (dd, $J$ = 16.5, 7.3 Hz, 1H), 3.60 (d, $J$ = 13.7 Hz, 4H), 3.48 (d, $J$ = 11.3 Hz, 3H), 3.32 – 3.06 (m, 6H), 3.05 (s, 3H), 2.80 – 2.68 (m, 4H), 2.61 (s, 3H), 2.42 (s, 3H), 2.22 – 2.09 (m, 3H), 1.92 – 1.80 (m, 4H), 1.63 (s, 3H), 1.59 (dd, $J$ = 13.2, 5.0 Hz, 6H), 1.25 – 1.16 (m, 1H), 1.11 – 1.05 (m, 1H).

Synthesis of (*S*)-2-(4-(4-chlorophenyl)-2,3,9-trimethyl-6*H*-thieno[3,2-*f*][1,2,4]triazolo[4,3-*a*][1,4]diazepin-6-yl)-1-(4-((4-(2-(4-((2-(3-((4-(dimethylphosphoryl)phenyl)(methyl)amino)prop-1-yn-1-yl)-1-(2,2,2-trifluoroethyl)-1*H*-indol-4-yl)(methyl)amino)piperidin-1-yl)acetyl)piperazin-1-yl)methyl)piperidin-1-yl)ethan-1-one (**TRAP-2-Neg2** or **XJZ-06-489**)

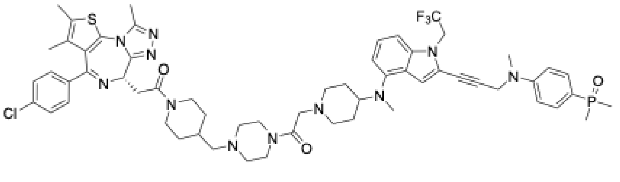

XJZ-06-489

The corresponding compound was prepared following **Procedure C** using intermediate **16** and compound **30** (5 mg, 8.5 µmol, 1 eq.). Purification by HPLC with MeOH/$H_2O$ (0.035% TFA) to afford the product as a white solid (4.33 mg, 45%). LC/MS(ESI) for $C_{59}H_{71}ClF_3N_{11}O_3PS$ $[M+H]^+$: m/z calcd, 1136.48; found, 1136.37 $[M+1]^+$. $^1$H NMR (500 MHz, d$^6$-DMSO) δ 9.74 – 9.44 (m, 1H), 7.63 – 7.54 (m, 2H), 7.50 (dd, $J$ = 8.8, 2.6 Hz, 2H), 7.44 (dd, $J$ = 8.6, 2.5 Hz, 2H), 7.18 – 7.10 (m, 2H), 7.00 (dd, $J$ = 8.9, 2.0 Hz, 2H), 6.94 – 6.82 (m, 1H), 6.74 – 6.59 (m, 1H), 4.96 (q, $J$ = 9.0 Hz, 2H), 4.61 – 4.55 (m, 3H), 4.45 – 4.32 (m, 2H), 4.31 – 4.10 (m, 2H), 3.85 – 3.76 (m, 1H), 3.69 – 3.56 (m, 3H), 3.52 – 3.45 (m, 3H), 3.43 – 3.34 (m, 1H), 3.33 – 3.21 (m, 1H), 3.22 – 2.90 (m, 11H), 2.81 – 2.71 (m, 3H), 2.69 – 2.61 (m, 1H), 2.60 (s, 3H), 2.42 (s, 3H), 2.23 – 2.08 (m, 3H), 1.93 – 1.83 (m, 3H), 1.81 – 1.73 (m, 1H), 1.63 (s, 3H), 1.58 (d, $J$ = 13.2 Hz, 6H), 1.37 – 1.19 (m, 1H), 1.14 – 0.98 (m, 1H).

Synthesis of (S)-2-(4-(4-chlorophenyl)-2,3,9-trimethyl-6H-thieno[3,2-f][1,2,4]triazolo[4,3-a][1,4]diazepin-6-yl)-1-(9-(2-(4-((2-(3-((4-(dimethylphosphoryl)phenyl)(methyl)amino)prop-1-yn-1-yl)-1-(2,2,2-trifluoroethyl)-1H-indol-4-yl)(methyl)amino)piperidin-1-yl)acetyl)-3,9-diazaspiro[5.5]undecan-3-yl)ethan-1-one (**TRAP-3-Neg2** or **XJZ-06-490**)

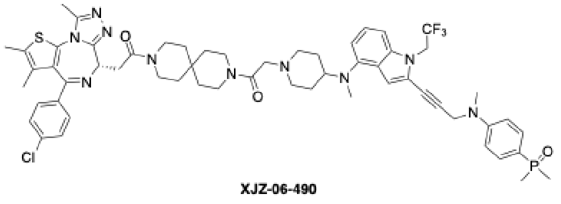

XJZ-06-490

The corresponding compound was prepared following **Procedure C** using intermediate **17** and compound **30** (5 mg, 8.5 µmol, 1 eq.). Purification by HPLC with MeOH/$H_2O$ (0.035% TFA) to afford the product as a white solid (4.25 mg, 45%). LC/MS(ESI) for $C_{58}H_{68}ClF_3N_{10}O_3PS$ $[M+H]^+$: m/z calcd, 1107.46; found, 1107.45. $^1$H NMR (500 MHz, d$^6$-DMSO) δ 9.41 (d, $J$ = 46.2 Hz, 1H), 7.63 – 7.54 (m, 2H), 7.53 – 7.48 (m, 2H), 7.46 – 7.41 (m, 2H), 7.18 – 7.09 (m, 2H), 7.00 (dd, $J$ = 8.9, 2.0 Hz, 2H), 6.88 (d, $J$ = 47.5 Hz, 1H), 6.62 (d, $J$ = 7.1 Hz, 1H), 4.96 (q, $J$ = 9.0 Hz, 2H), 4.62 – 4.55 (m, 3H), 4.37 – 4.18 (m, 2H), 3.85 – 3.74 (m, 1H), 3.69 – 3.59 (m, 3H), 3.56 – 3.46 (m,

5H), 3.38 (dd, *J* = 16.3, 6.1 Hz, 1H), 3.34 – 3.24 (m, 2H), 3.13 – 3.05 (m, 2H), 3.04 (s, 3H), 2.80 – 2.73 (m, 3H), 2.60 (s, 3H), 2.42 (s, 3H), 2.22 – 2.09 (m, 2H), 1.91 – 1.85 (m, 2H), 1.63 (s, 3H), 1.61 – 1.55 (m, 8H), 1.55 – 1.50 (m, 2H), 1.50 – 1.45 (m, 2H), 1.44 – 1.39 (m, 2H).

### 2. NMR Spectra

$^{1}$H NMR of intermediate **4** (500 MHz, $CDCl_3$)

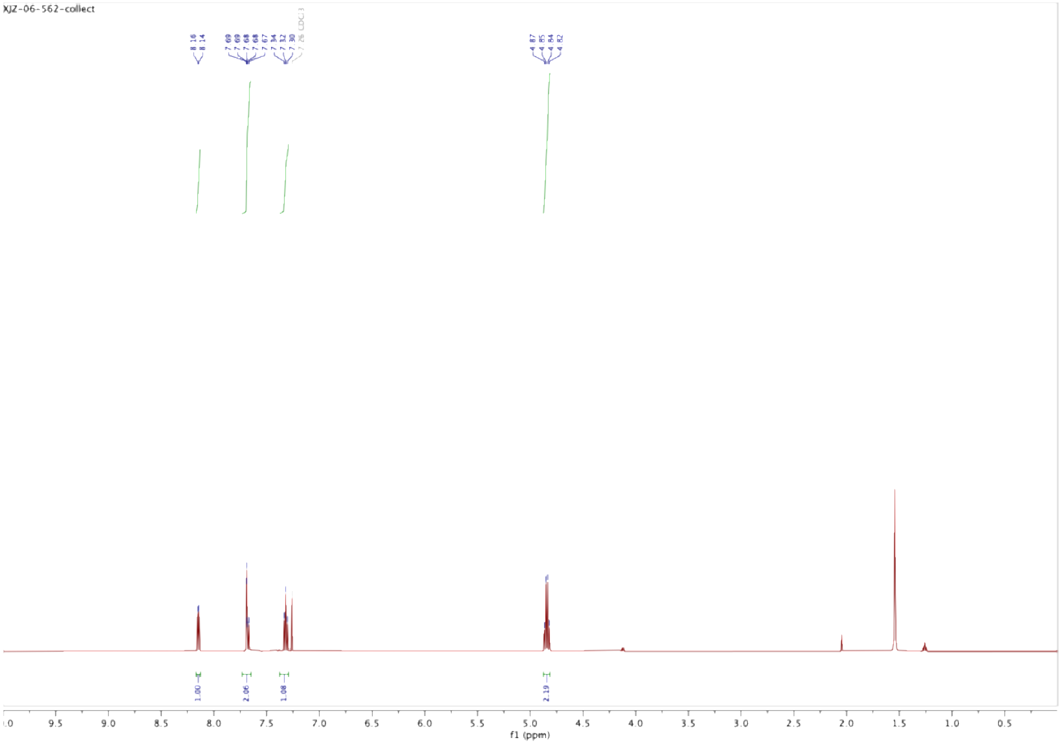

$^{1}$H NMR of **B-1 linker** (500 MHz, $d^6$-DMSO)

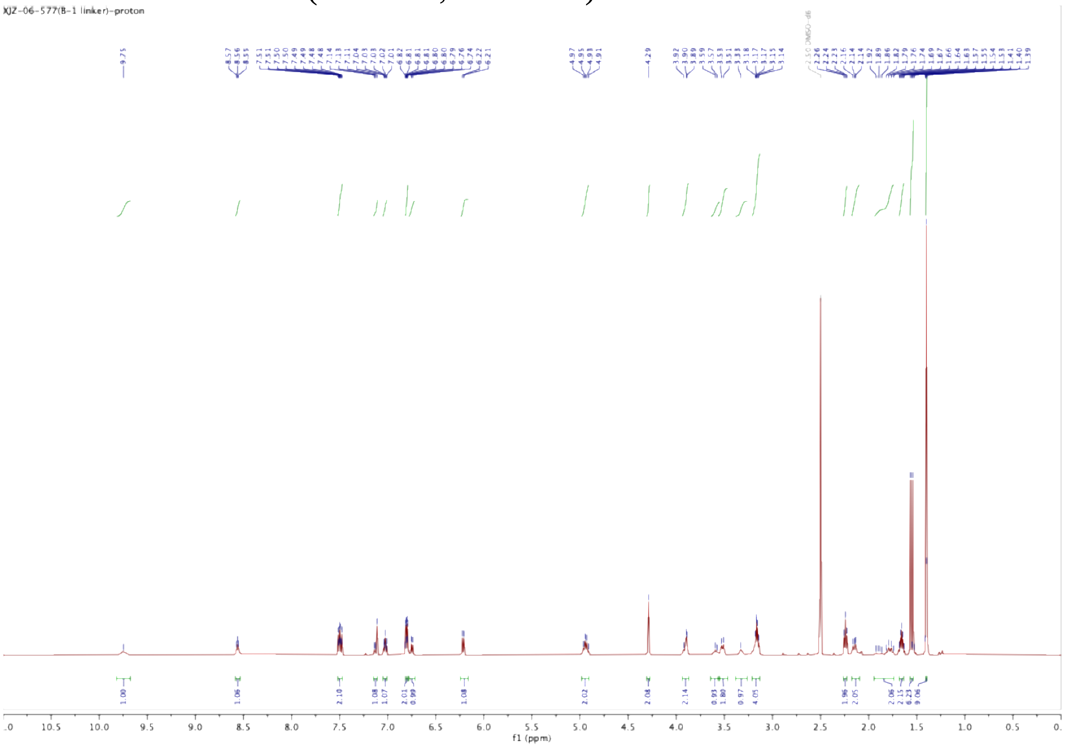

$^1$H NMR of **XJZ-06-444** (500 MHz, d$^6$-DMSO)

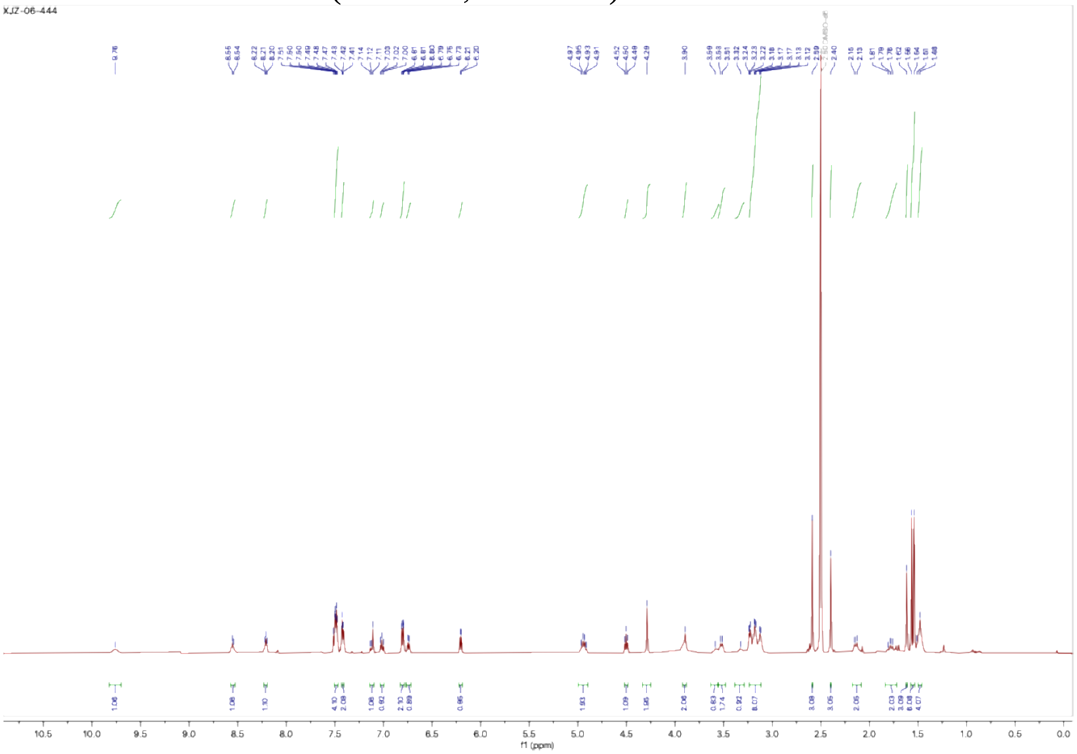

$^1$H NMR of **XJZ-06-445** (500 MHz, d$^6$-DMSO)

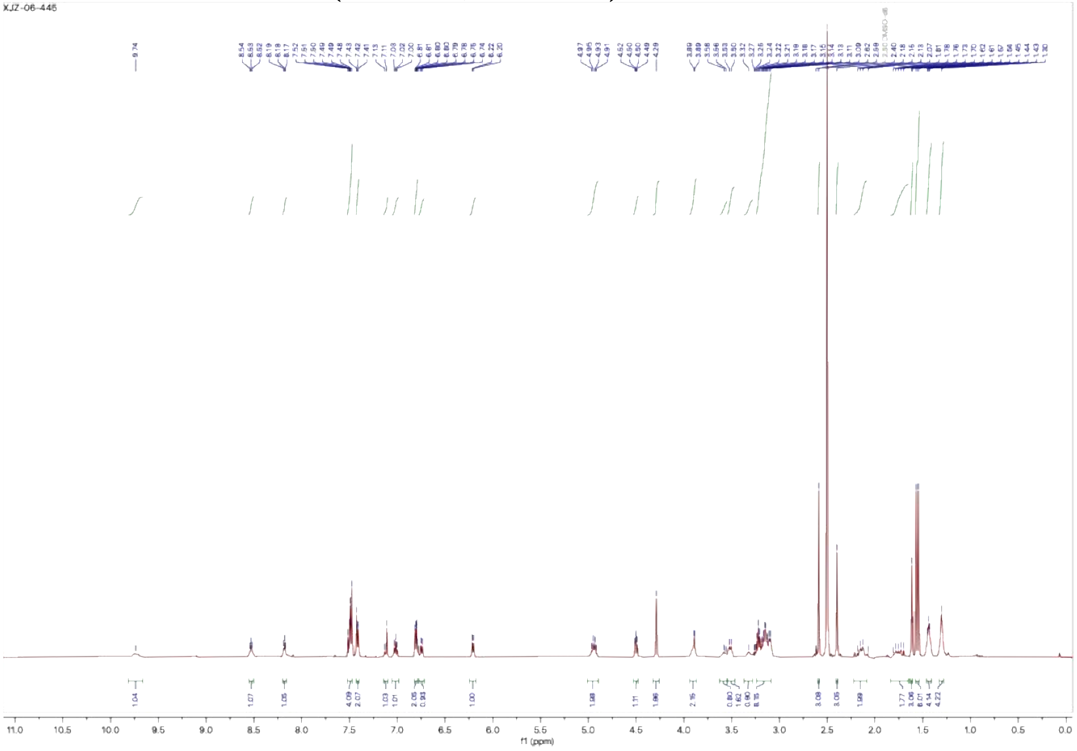

$^1$H NMR of **XJZ-06-446** (500 MHz, $d^6$-DMSO)

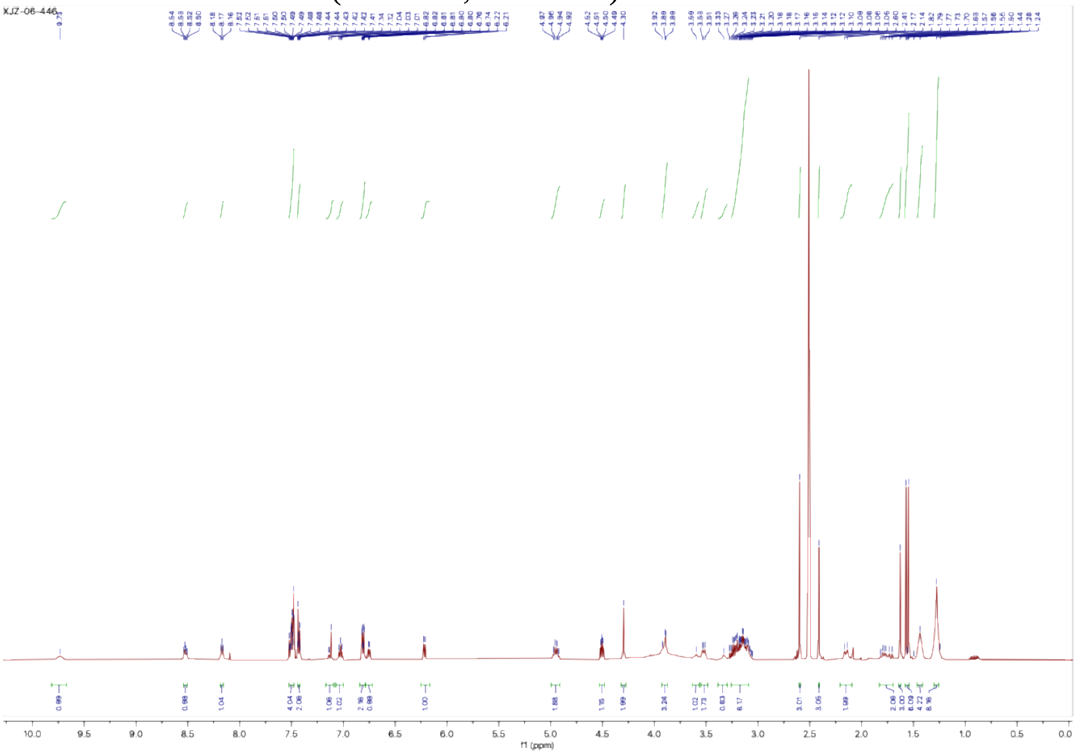

$^1$H NMR of **XJZ-06-448** (500 MHz, $d^6$-DMSO)

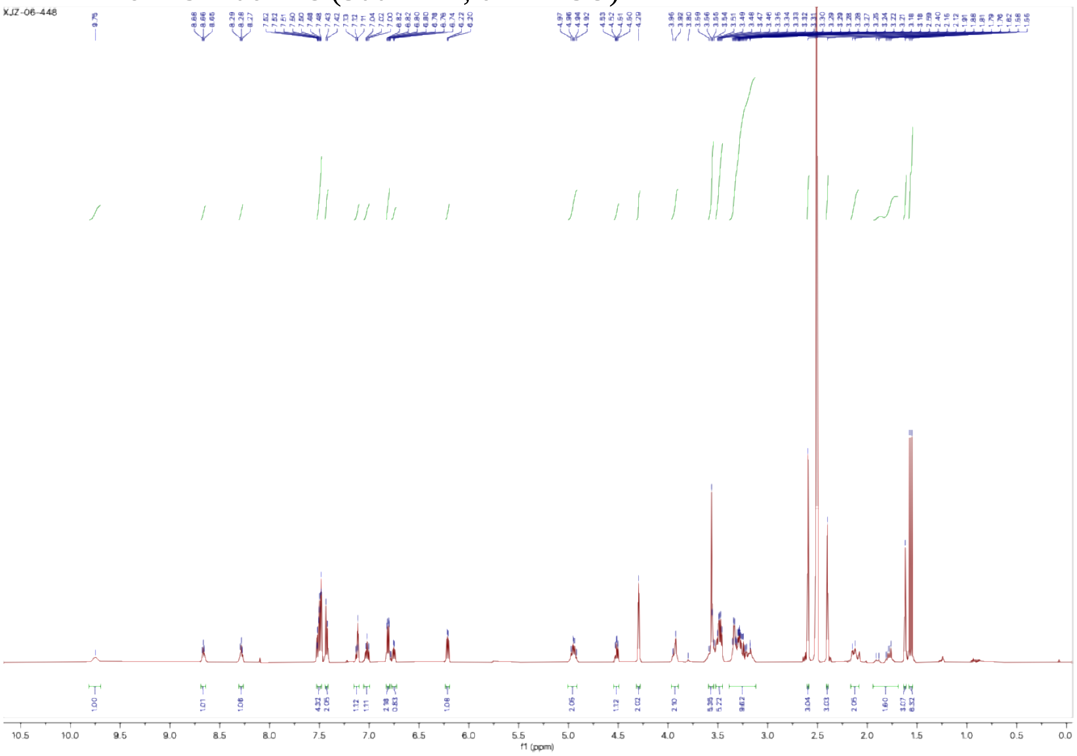

$^{1}$H NMR of **XJZ-06-449** (500 MHz, $d^6$-DMSO)

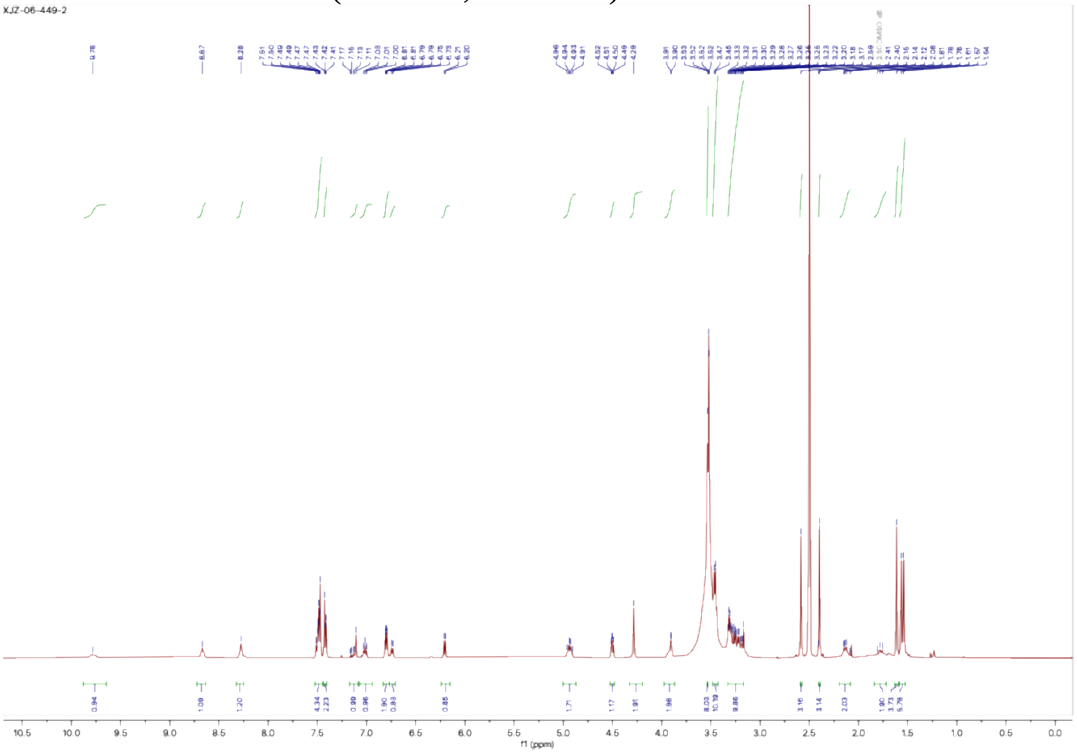


$^{1}$H NMR of **TRAP-1** (**XJZ-06-462**) (500 MHz, $d^6$-DMSO)

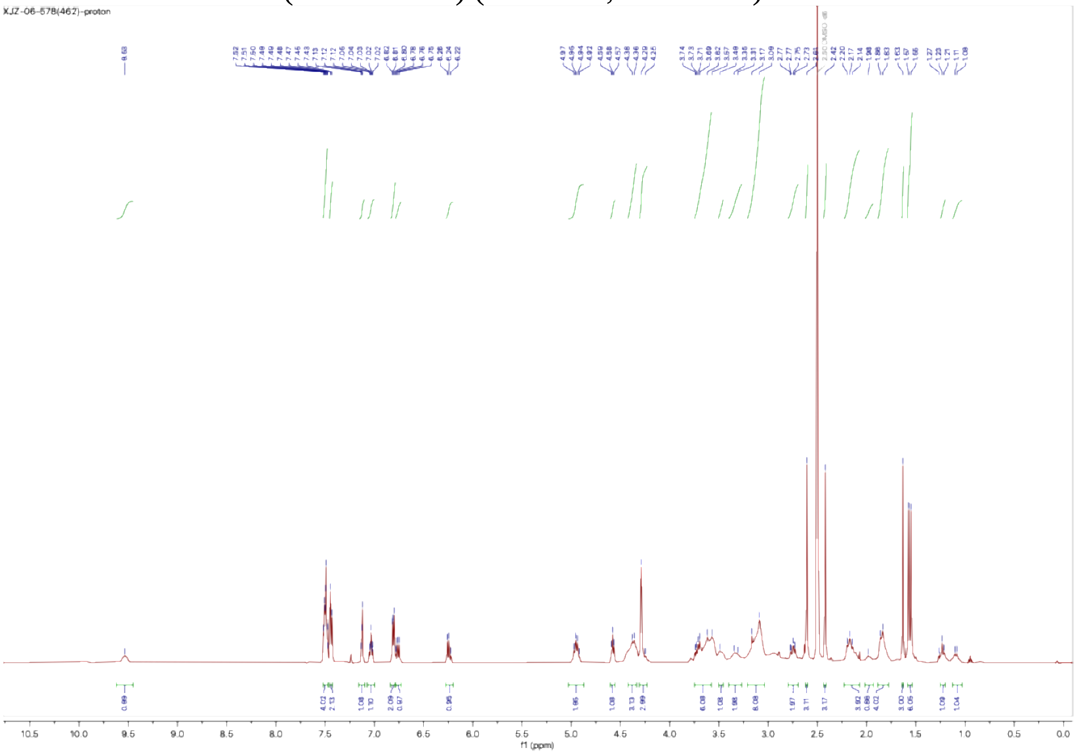

$^{1}$H NMR of **TRAP-2** (**XJZ-06-463**) (500 MHz, $d^6$-DMSO)

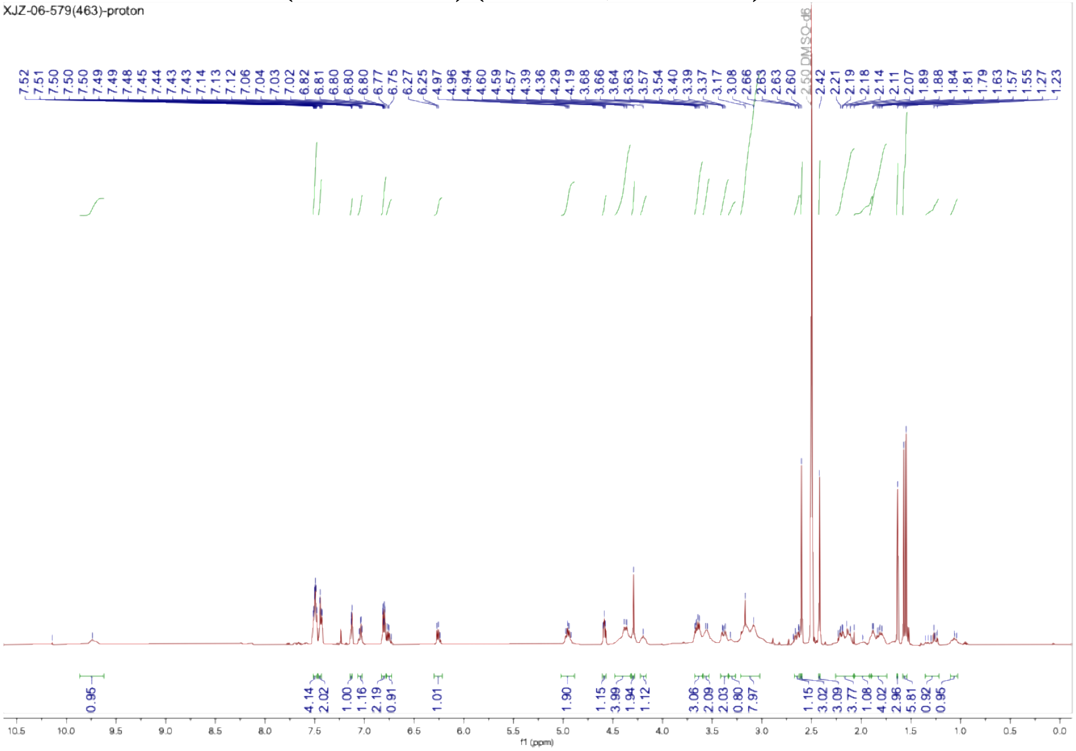

$^{1}$H NMR of **TRAP-3** (**XJZ-06-464**) (500 MHz, $d^6$-DMSO)

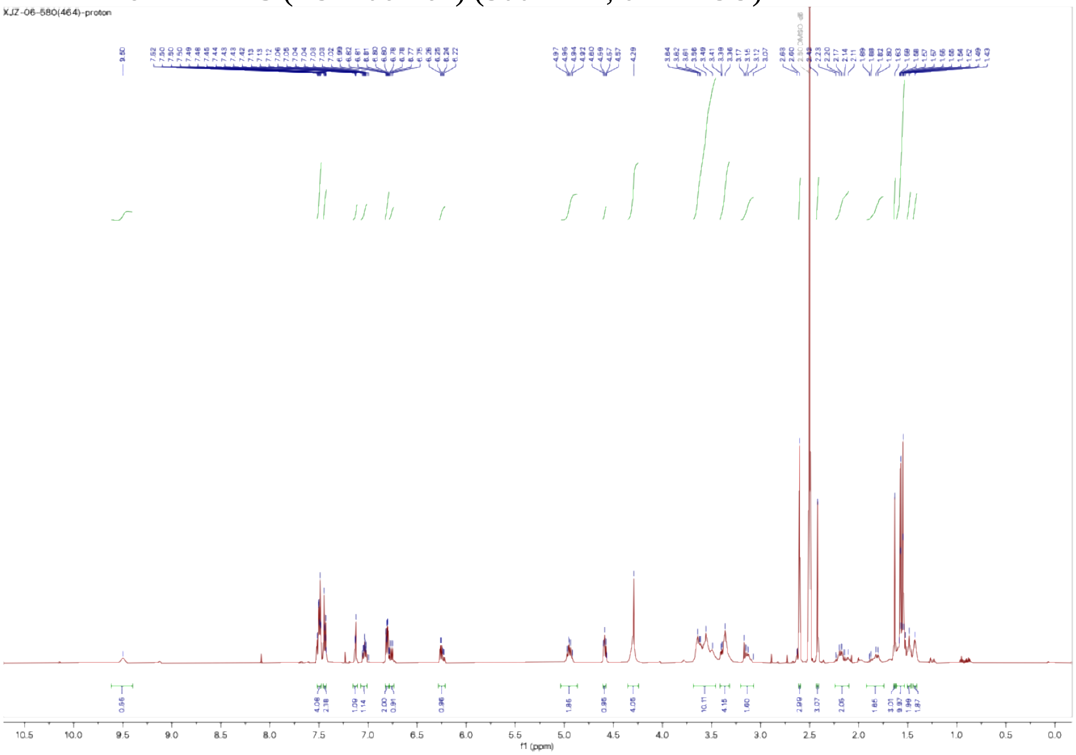

$^{1}$H NMR of **XJZ-06-465** (500 MHz, d$^{6}$-DMSO)

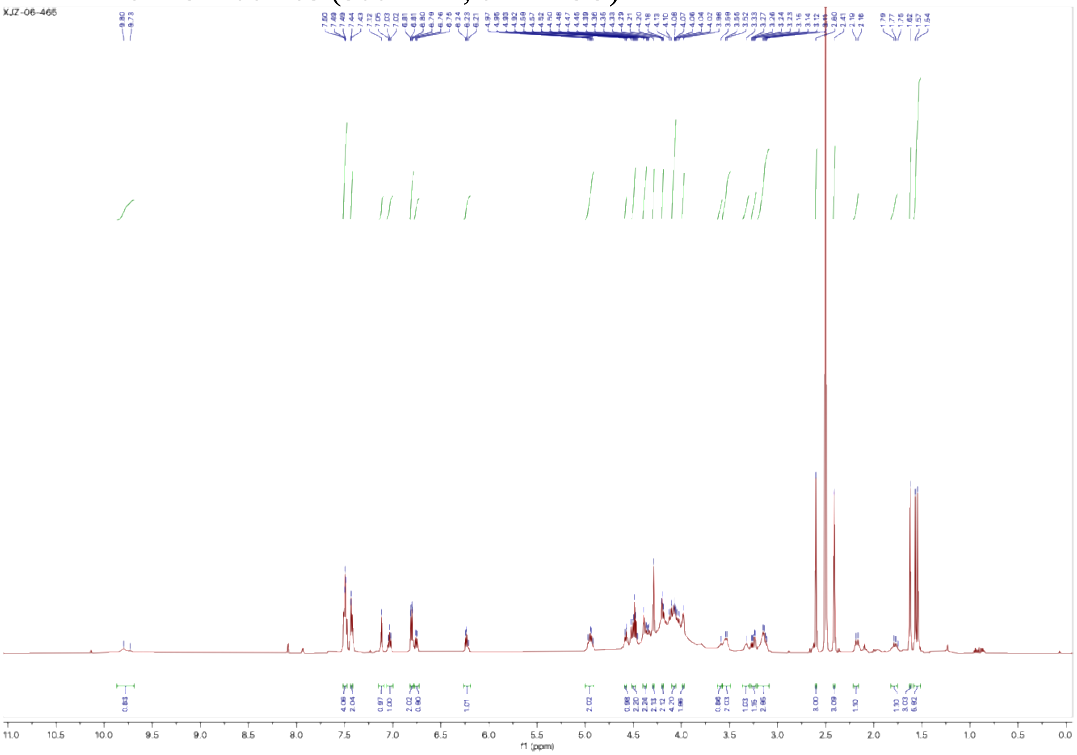

$^{1}$H NMR of **XJZ-06-471** (500 MHz, d$^{6}$-DMSO)

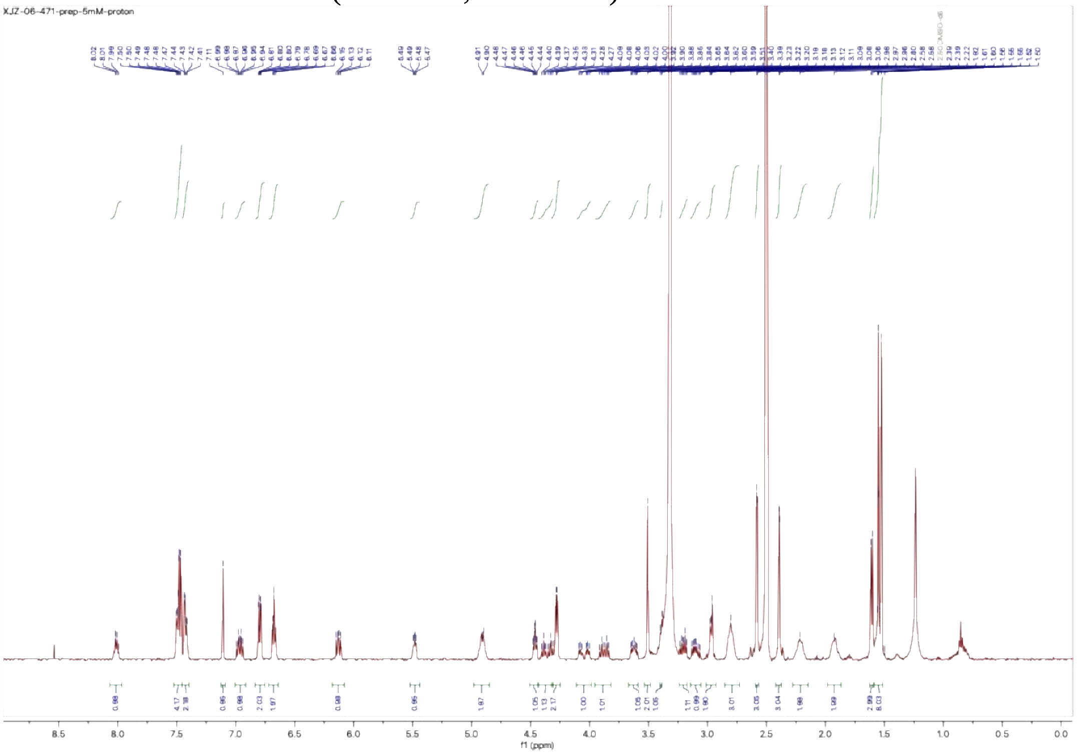

$^{1}$H NMR of **XJZ-06-537** (500 MHz, d$^{6}$-DMSO)

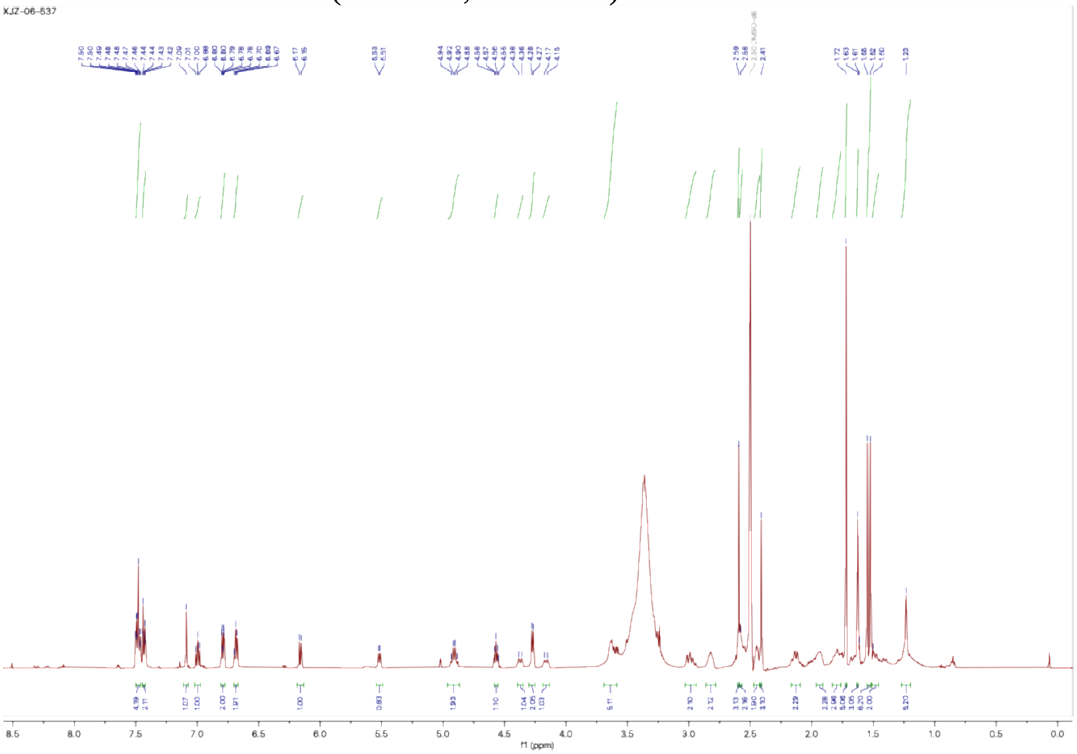


$^{1}$H NMR of **XJZ-06-540** (500 MHz, d$^{6}$-DMSO)

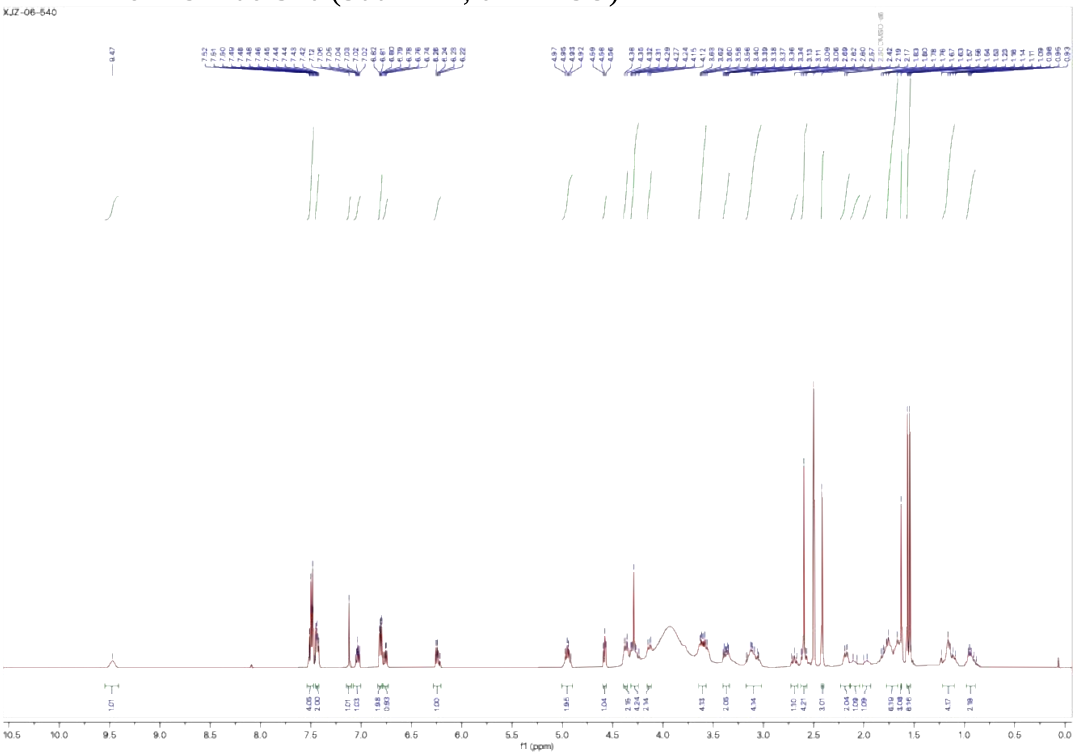

$^{1}$H NMR of **PK9328 free amine** (500 MHz, $d^6$-DMSO)

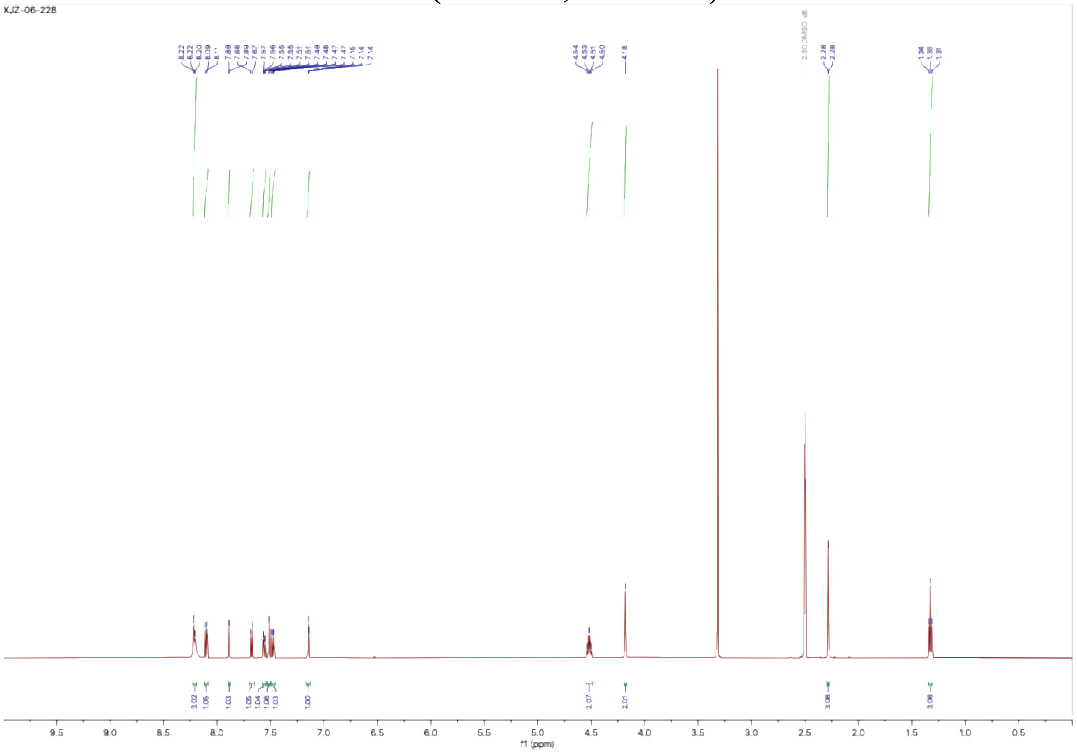

$^{1}$H NMR of **XJZ-06-408** (500 MHz, $d^6$-DMSO)

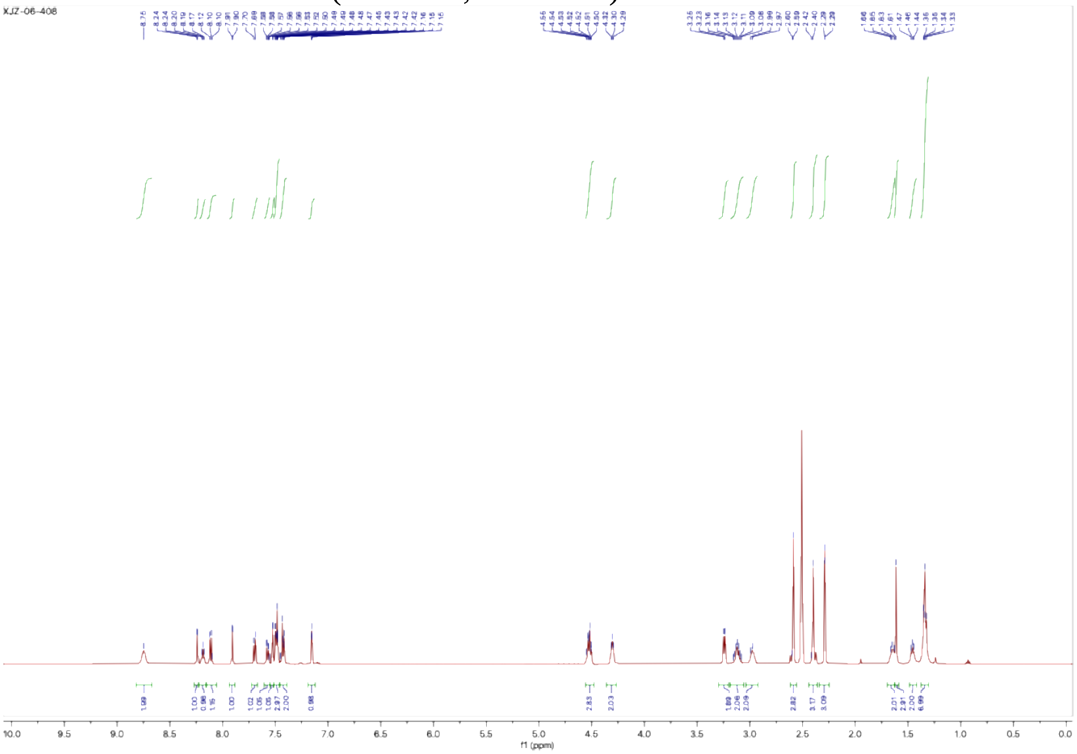

$^{1}$H NMR of **XJZ-06-409** (500 MHz, $d^6$-DMSO)

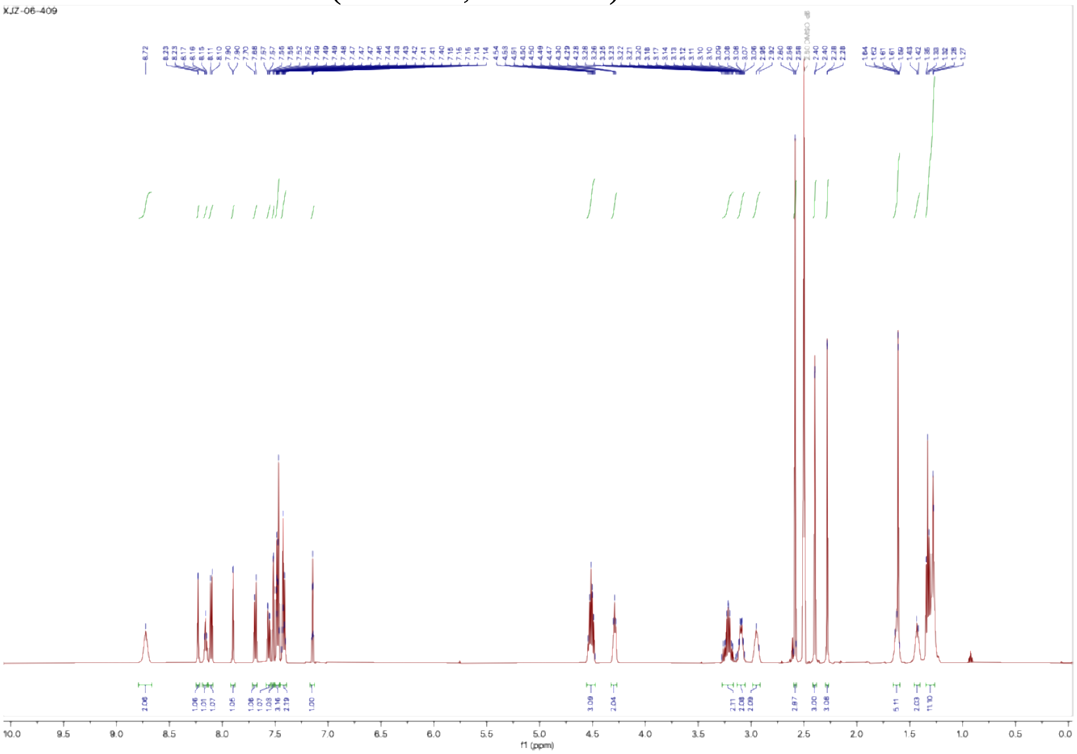

$^{1}$H NMR of **XJZ-06-411** (500 MHz, $d^6$-DMSO)

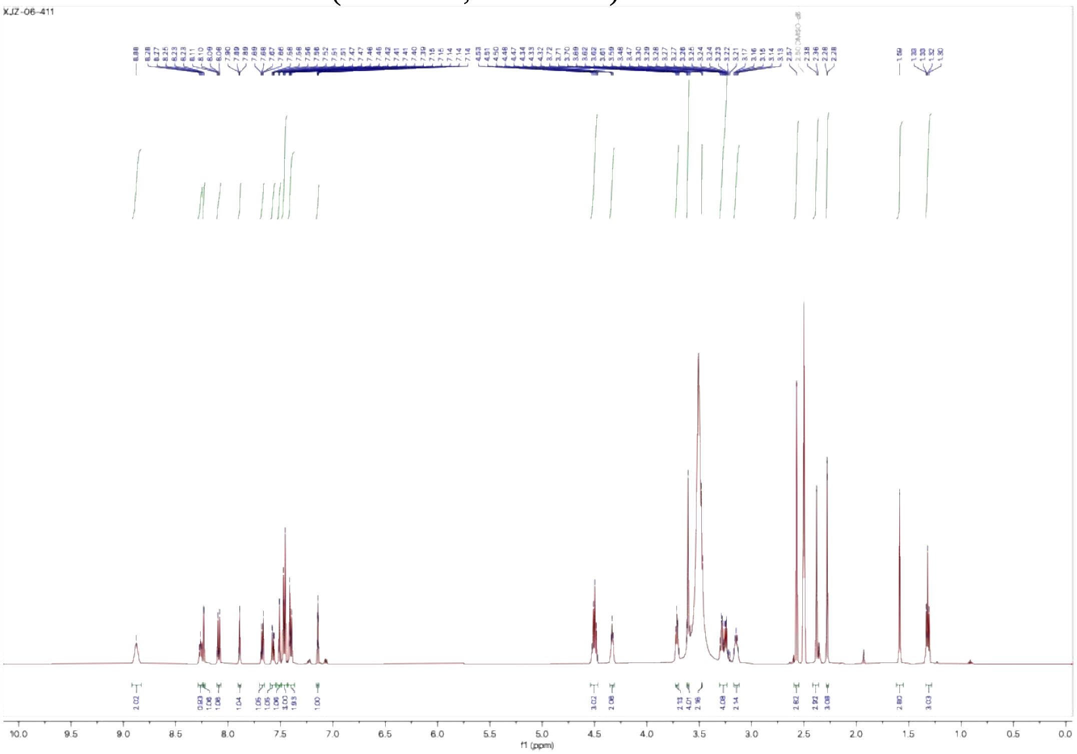

$^1$H NMR of **XJZ-06-392** (500 MHz, $d^6$-DMSO)

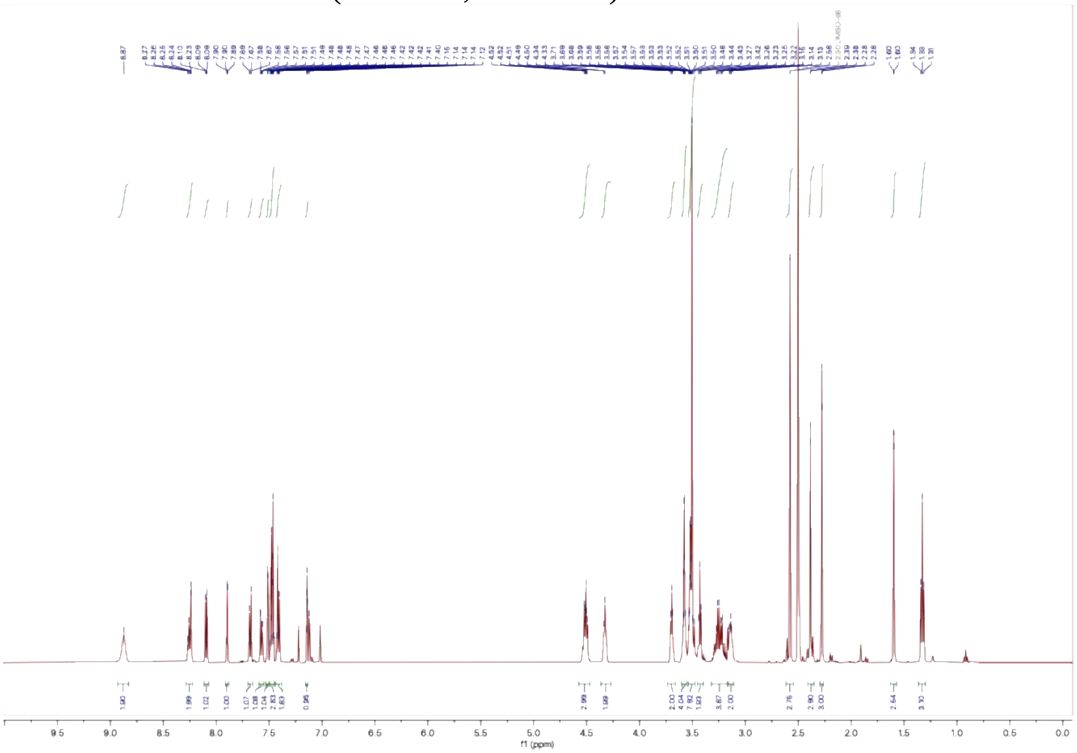

$^1$H NMR of **XJZ-06-429** (500 MHz, $d^6$-DMSO)

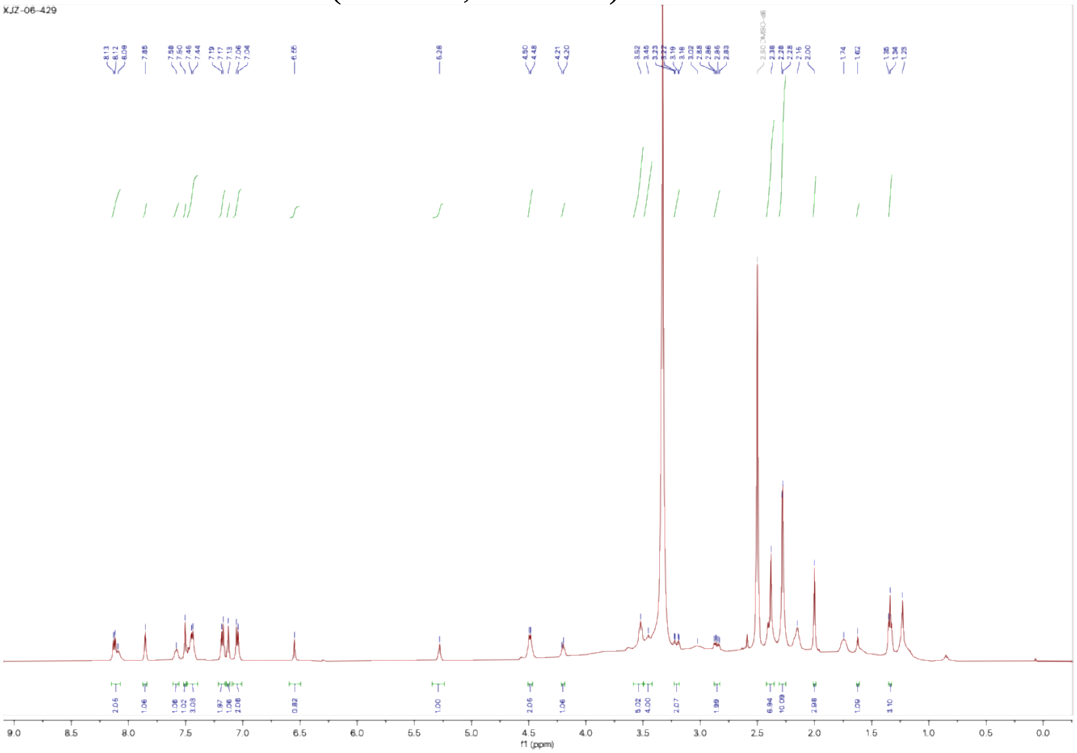

$^{1}$H NMR of **XJZ-06-430** (500 MHz, d$^{6}$-DMSO)

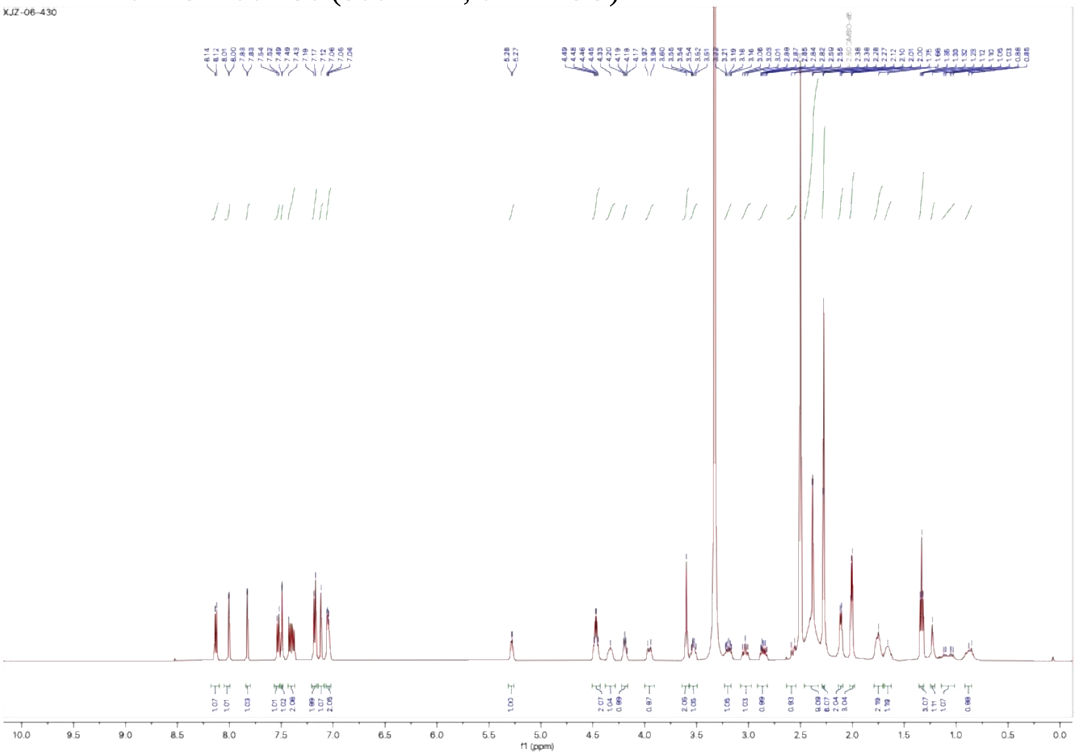

$^{1}$H NMR of **XJZ-06-431** (500 MHz, d$^{6}$-DMSO)

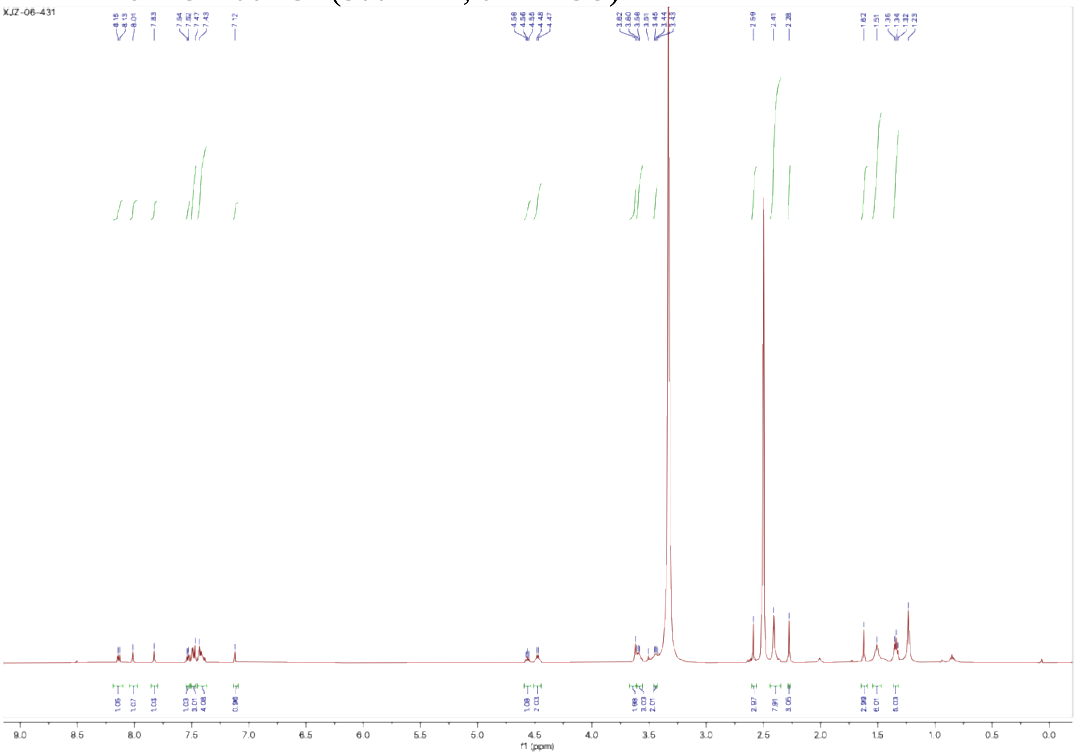

$^1$H NMR of **XJZ-06-432** (500 MHz, $d^6$-DMSO)

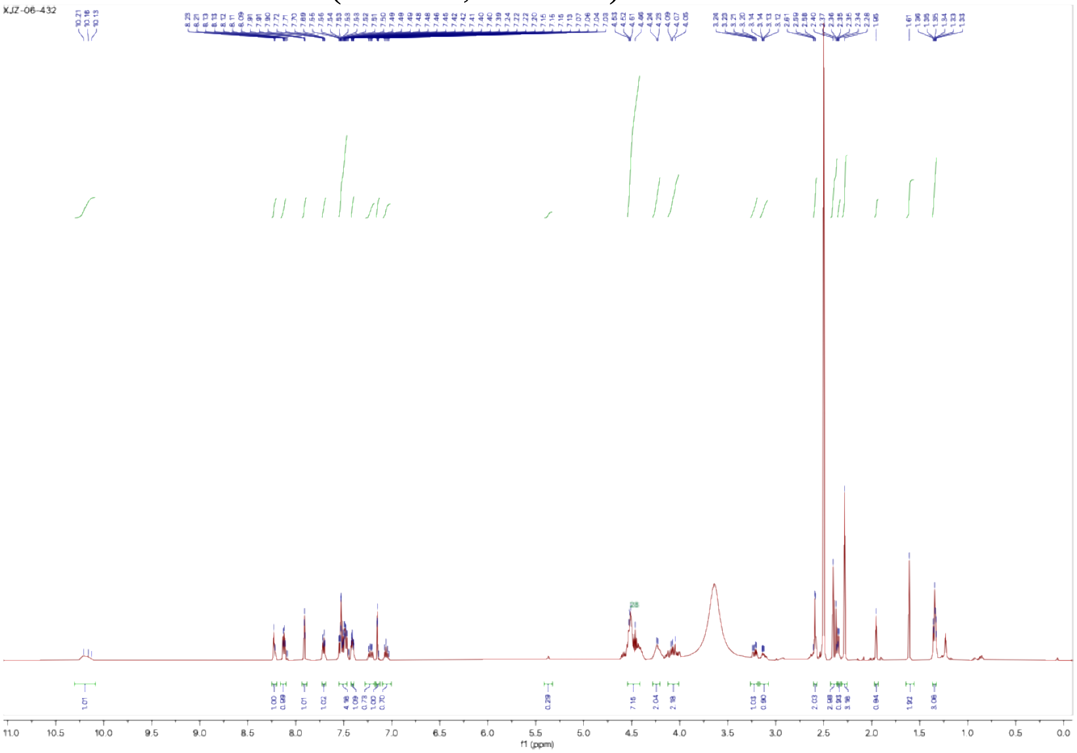

$^1$H NMR of **TRAP-1-Neg1** (**XJZ-06-483**) (500 MHz, $d^6$-DMSO)

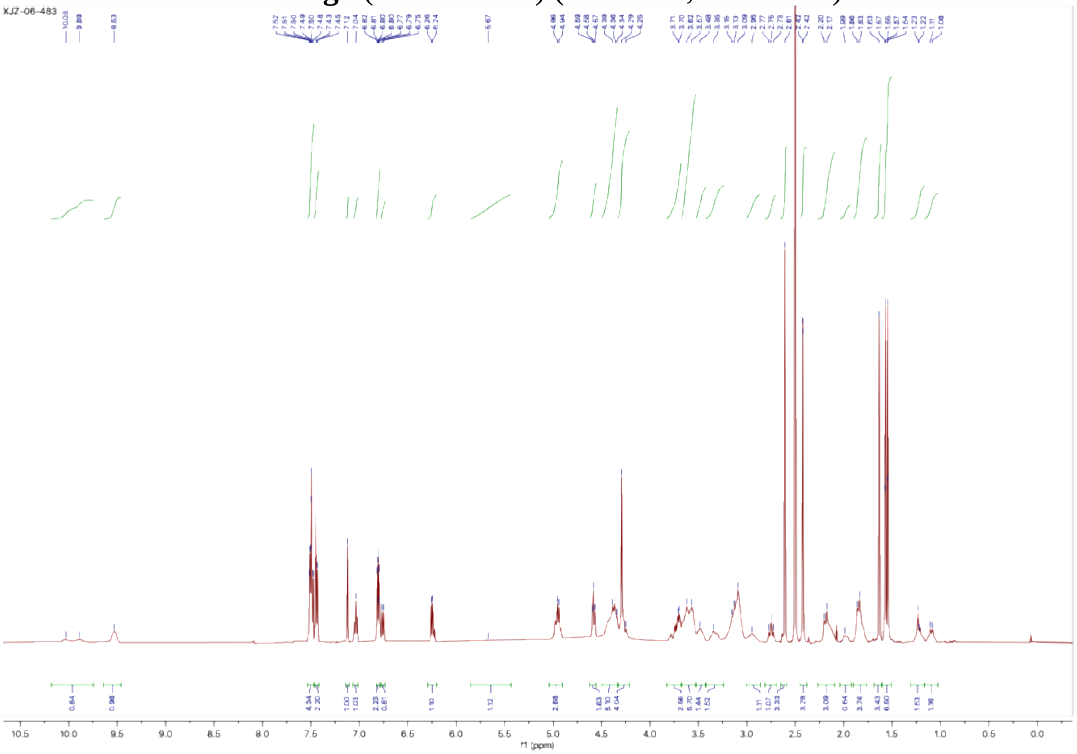

$^{1}H$ NMR of **TRAP-2-Neg1** (**XJZ-06-484**) (500 MHz, $d^6$-DMSO)

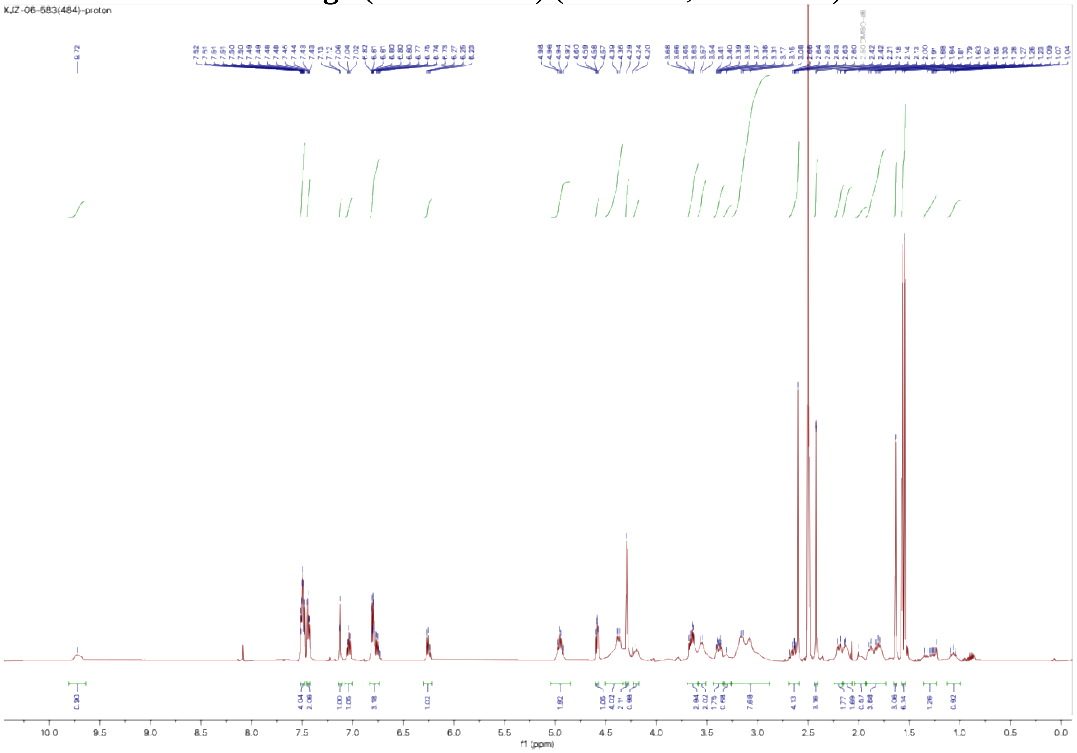

$^{1}H$ NMR of **TRAP-3-Neg1** (**XJZ-06-485**) (500 MHz, $d^6$-DMSO)

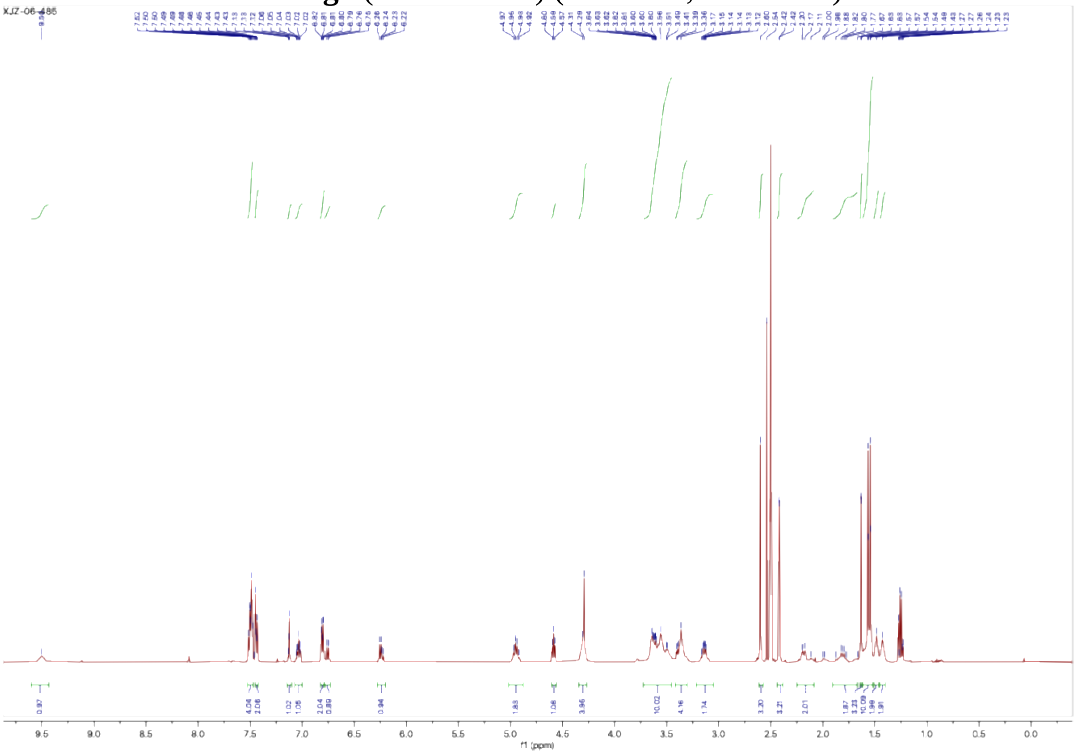

$^1$H NMR of **TRAP-1-Neg2** (**XJZ-06-488**) (500 MHz, d$^6$-DMSO)

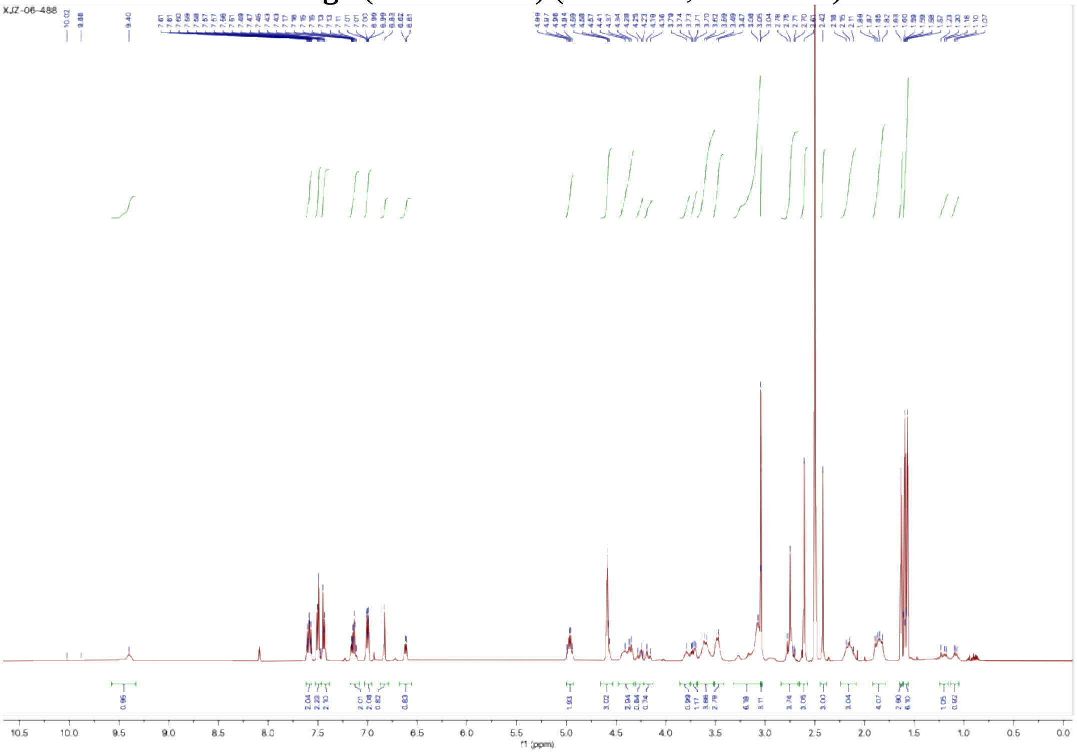

$^1$H NMR of **TRAP-2-Neg2** (**XJZ-06-489**) (500 MHz, d$^6$-DMSO)

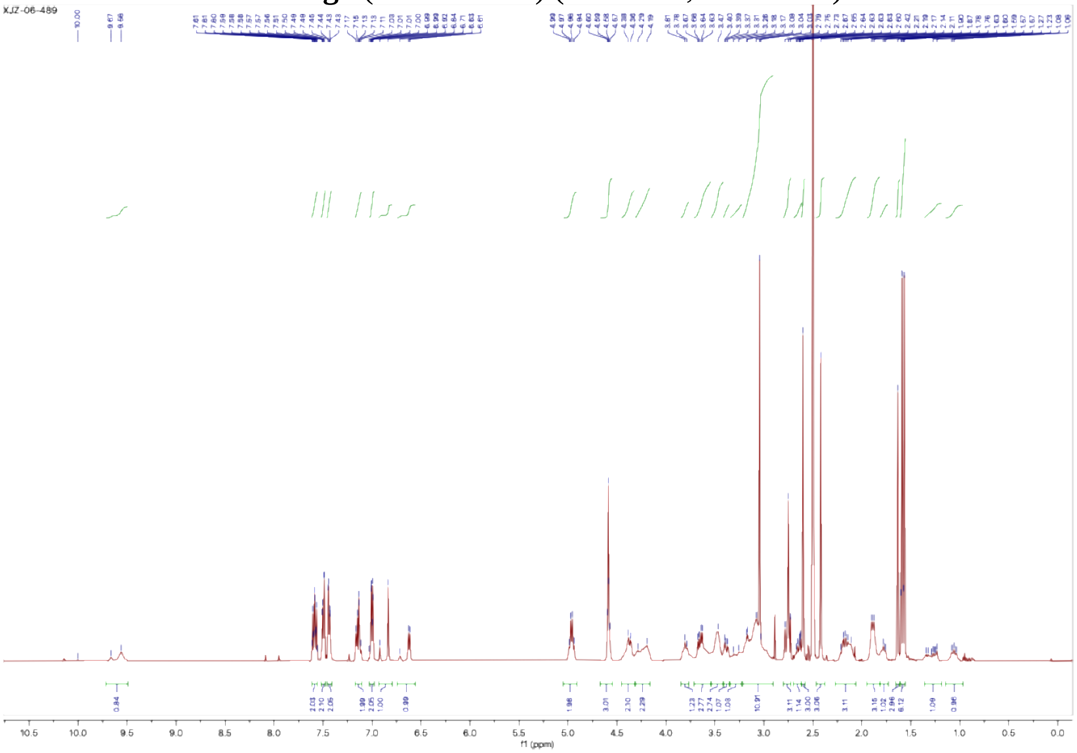

$^{1}H$ NMR of **TRAP-3-Neg2** (**XJZ-06-490**) (500 MHz, $d^6$-DMSO)

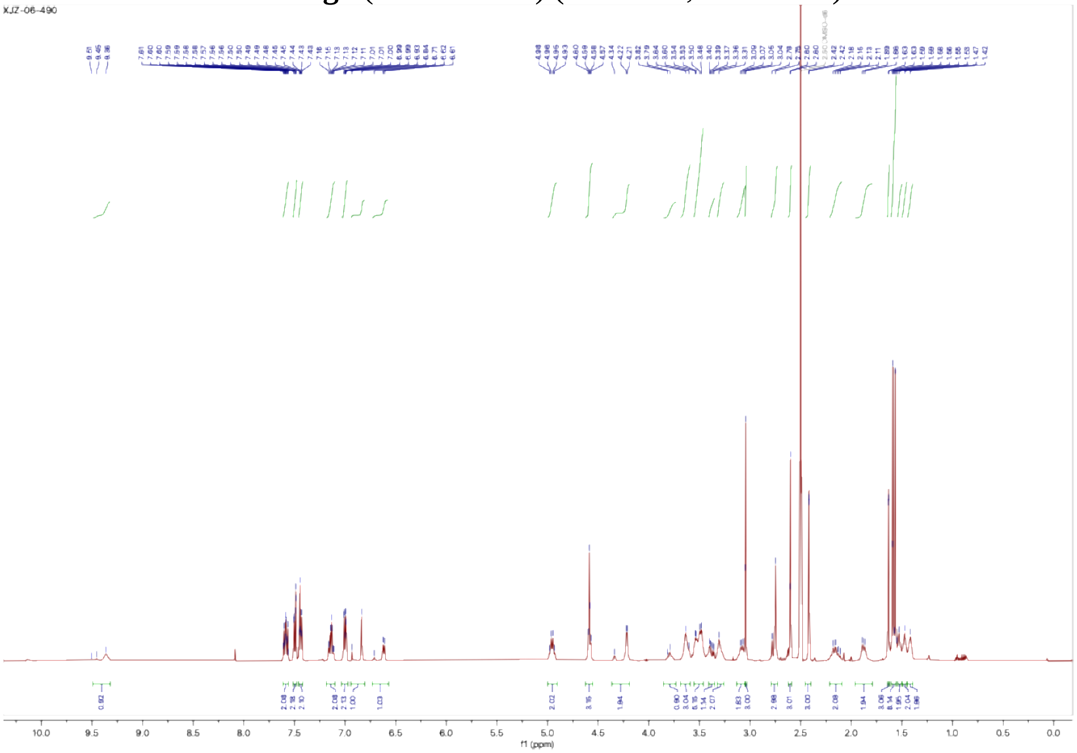

### 3. Uncropped Western Blots

Uncropped Blot for Figure 3D

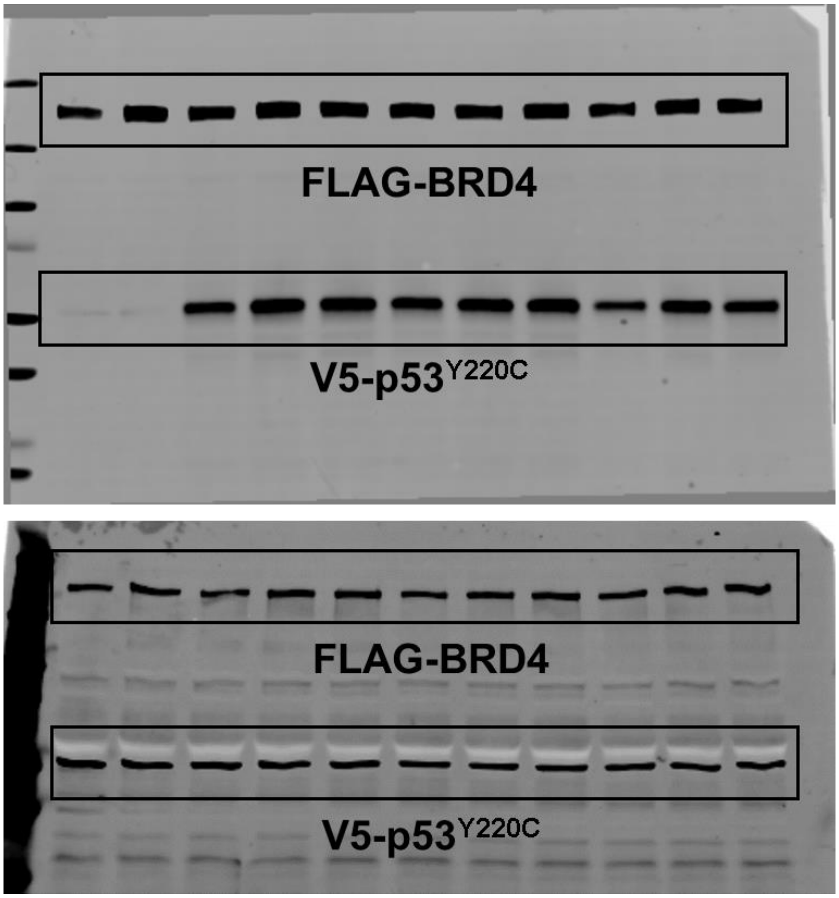


Uncropped Blot for Figure 4B

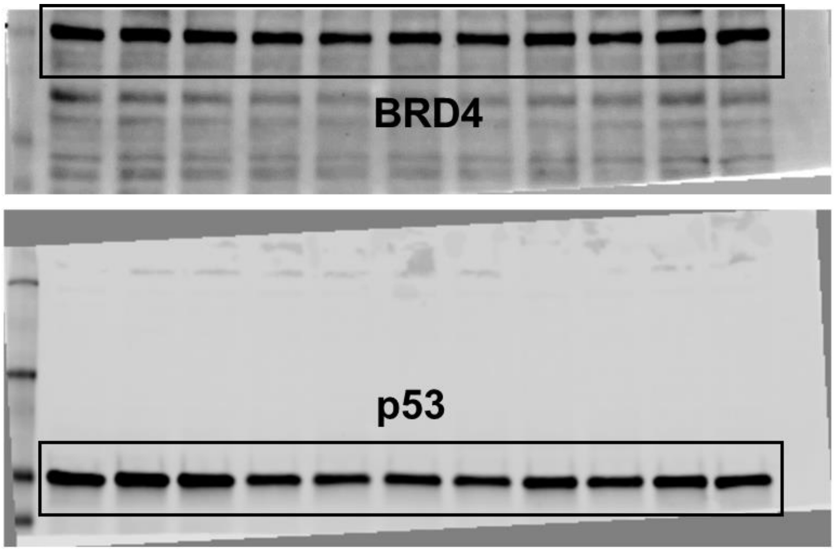

Uncropped Blot for Figure 4B (Continued)

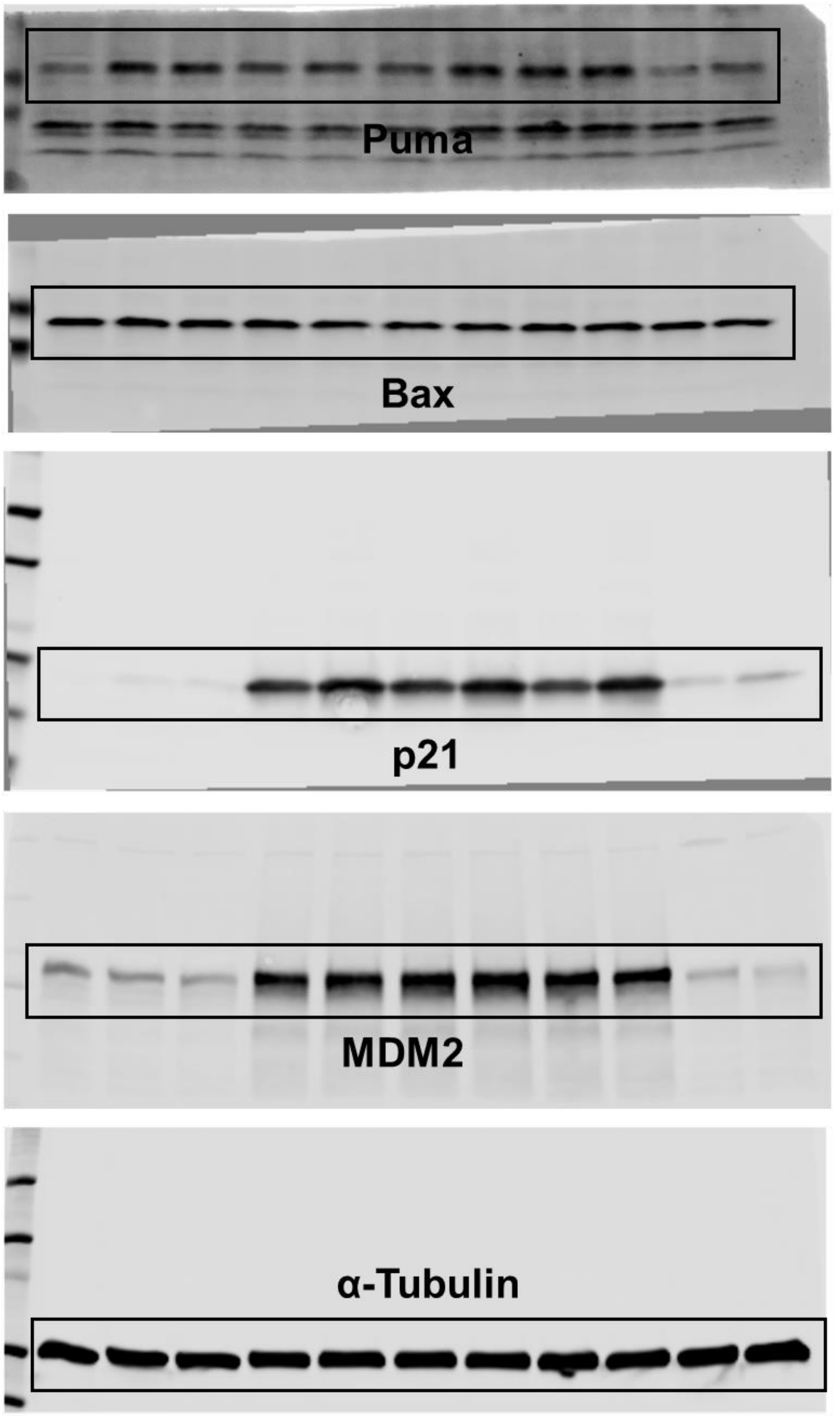

Uncropped Blot for Figure 4C

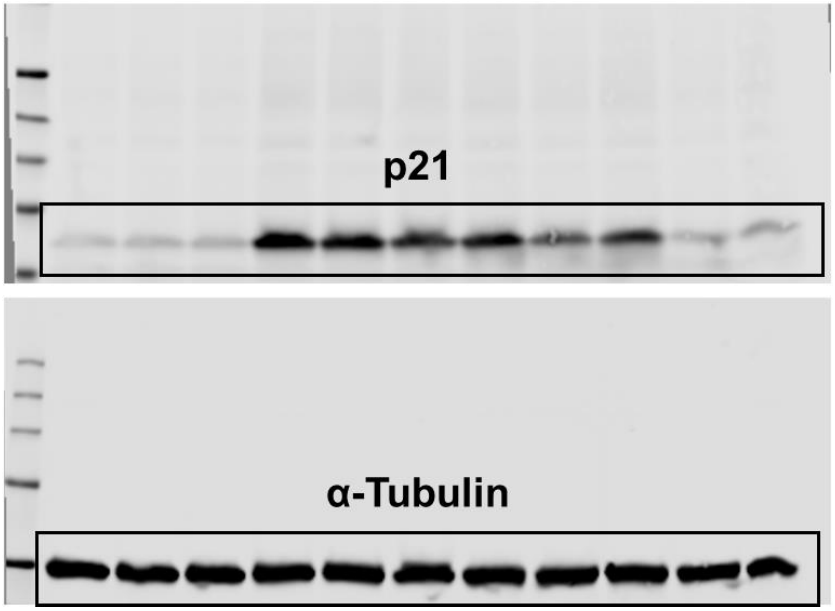
